# Coherence from Context – Two-Point Neuron Models for Contextual Integration in Visual Information Processing

**DOI:** 10.64898/2026.04.27.721094

**Authors:** Daniel Schmid, Heiko Neumann

## Abstract

The integration of contextual information is crucial for coherent perception and cognition. The morphology and conductance properties of cortical pyramidal cells suggest that they operate as “two-point neurons” (TPNs), asymmetrically combining driving sensory input at basal dendrites with modulating context at apical compartments. We present a mechanistic rate-based TPN model of layer V pyramidal cell populations and their local PV, SOM, and VIP interneuron circuits. The model rests on asymmetric feedforward-feedback integration and pooled inhibition to implement local competition and global cooperation supporting the selective amplification of coherent signals. Fitted to multi-compartment pyramidal cell simulations, it reproduces their basal-apical input-output map and, unlike static transfer functions, shows how apical amplification unfolds over time under interneuron control. Asymmetric apical-basal coincidence integrates context during processing and tags TPNs for reward-driven learning with a local four-factor rule. A pool of TPNs thereby learns to use flanker context to decide ambiguous orientations. Context-driven VIP cells gate apical integration and learning through dis-inhibition. We connect TPN populations with joint spatial and feature selectivity into networks. Contextual signals from lateral recurrence and top-down feedback yield coherence formation and evidence propagation. This propagation disambiguates visual motion and requires apical-basal coupling. For larger-scale simulations, we show how established simpler models follow from reducing the TPN’s complexity while retaining its core principles. Overall, context in the TPN amplifies sensory evidence but cannot create it without feedforward input. Finally, we discuss when dynamics and interneurons matter, the model’s testable predictions for VIP silencing, and implications for artificial neural networks.

**Graphical Abstract:** 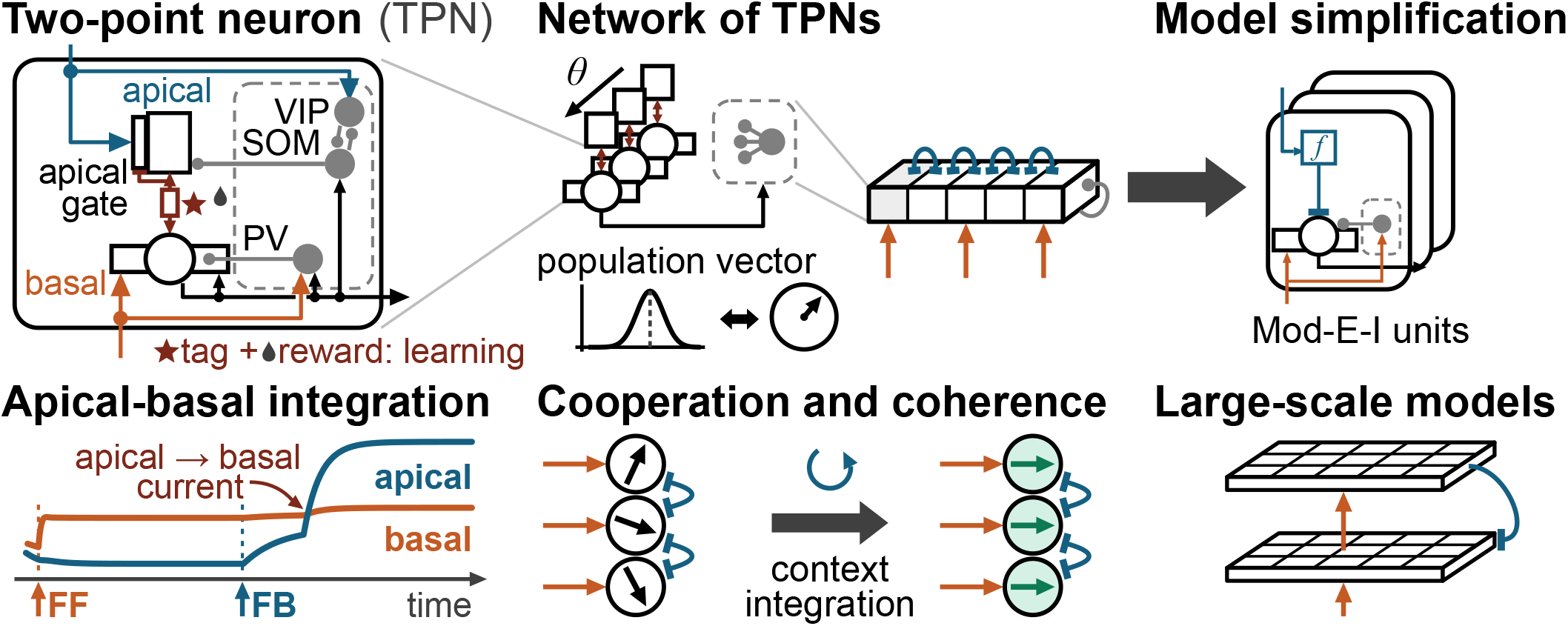

**Highlights:**

- Mechanistic TPN model with PV/SOM/VIP cells fits a multi-compartment L5 cell model.
- Asymmetric apical-basal coincidence underlies both context integration and learning.
- VIP disinhibition gates contextual amplification and reward-driven learning.
- Local competition and global cooperation via context make TPN networks coherent.
- Reduced models retain core computations for efficient large-scale vision networks.

## 1. Introduction

The neuron doctrine formulated by Ramón y Cajal forms the basis of modern neuroscience (Purves et al., 2018). It describes discrete neural cells and their synaptic connections as the constituent elements of brain function. Since then, much progress has been made at the cell level regarding anatomy, physiology and function, as well as at the network level regarding connectivity (Felleman and Van Essen, 1991) and functional motifs (Harris and Mrsic-Flogel, 2013). Investigations have identified many different neuron types in cortex whose neurotransmitters have excitatory effects, such as pyramidal cells and spiny stellate cells, or inhibitory effects. Among these neuron types, pyramidal cells have been found to account for the largest fraction of neurons in cortex (Purves et al., 2018). Furthermore, pyramidal cells possess a special morphology with electrically coupled but semi-independent basal and apical components (Larkum, 2013). Modeling investigations have tried to understand these cells and their function at different levels of detail ranging from microscopic investigations describing chemical and synaptic dynamics to meso- and macroscopic ones modeling whole populations and their interaction. These insights and levels of investigation become increasingly integrated with each other, and the proposal of two-point neurons (TPNs) (Phillips et al., 1995; Kay and Phillips, 1997) provides a useful conceptual framework for identifying concrete links between the cell and network level (Phillips, 2023).

The framework of TPNs revolves around thick-tufted cortical layer V pyramidal (L5Pyr) cells. It arrives at a level of description by which L5Pyr cells are captured by two cell compartments and their interaction. The proposed framework aims to integrate the cell-level properties of asymmetric interaction with the network-level properties of cooperation and competition. At the *cell level*, L5Pyr cells are a central cell type for understanding the brain’s information processing. Electrically coupled basal perisomatic and distal apical cell compartments connect with differing sources, integrate their input in separate zones and impact the cell’s output differently (Larkum, 2013; Kay and Phillips, 2020). An action potential that back-propagates into the apical dendrite and coincides with tuft input within a few milliseconds triggers a dendritic calcium spike, which turns single action potentials into bursts and enhances the cell’s firing rate (Larkum et al., 1999; Larkum, 2013; Fig. 1B). In awake animals, apical and somatic activity of L5Pyr cells are highly correlated (Beaulieu-Laroche et al., 2019; Francioni et al., 2019) and apical activity is causally linked to perceptual detection (Takahashi et al., 2016, 2020), whereas general anesthesia decouples the apical from the basal compartment (Suzuki and Larkum, 2020). Input to the apical tuft also controls associative learning, which is suppressed when dendritic activity is blocked (Doron et al., 2020), and feedback to the tuft has been proposed to tag synapses for reward-based plasticity (Roelfsema and Holtmaat, 2018). This apical-basal coupling is a candidate mechanism for understanding the integration of driving with contextual information (Kay and Phillips, 2020), learning (Sacramento et al., 2018; Maryada et al., 2025), and cooperation between cells (Phillips, 2023; Adeel et al., 2025), and it has even been linked to consciousness (Storm et al., 2024).

**Figure 1:**
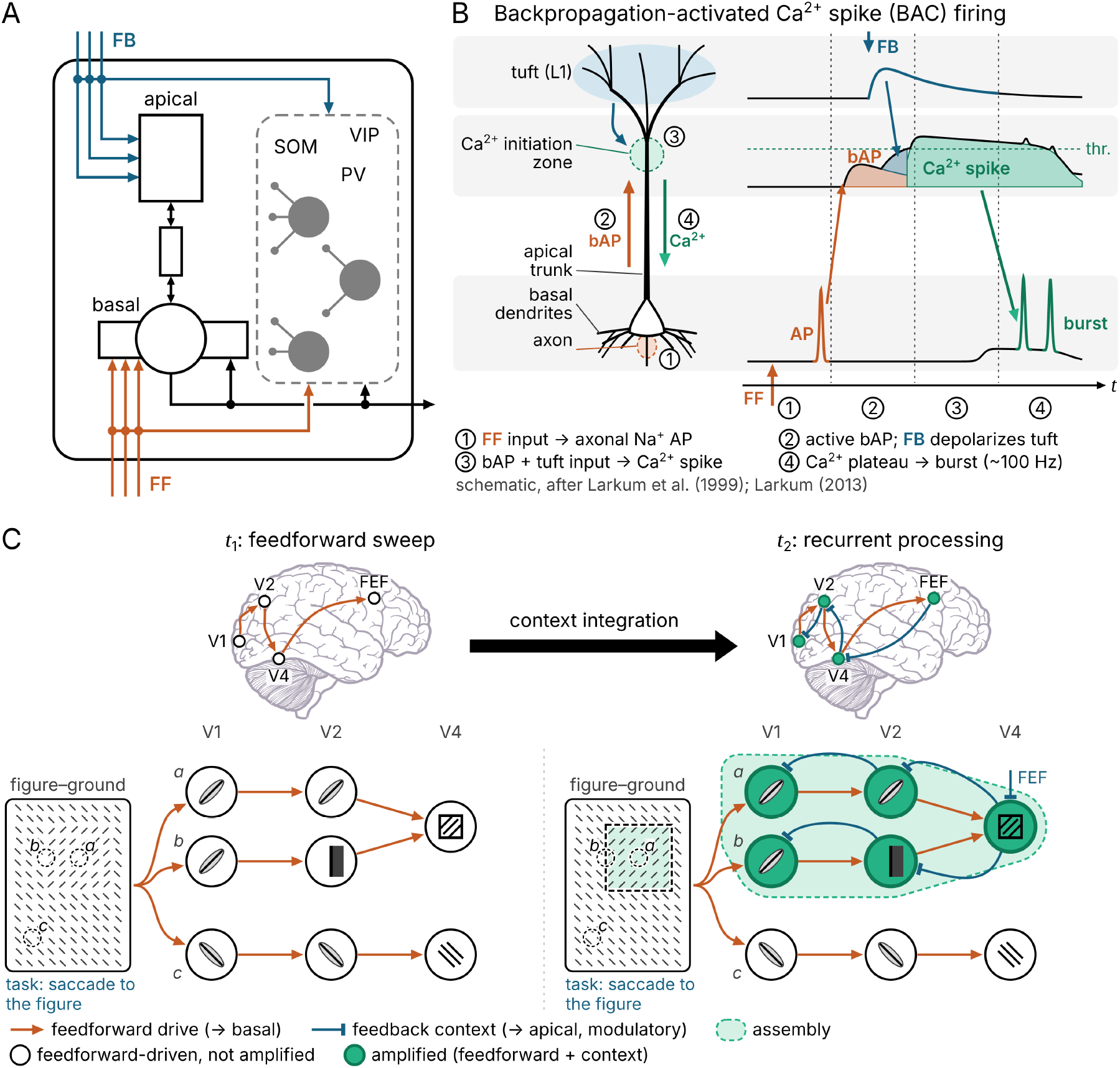
Pyramidal two-point neuron model, inhibitory interneurons, and cooperation among pyramidal cells. (A) Schematic of local pyramidal cell circuitry. Layer V pyramidal cells possess distinct basal and apical integration zones that are electrically coupled (rectangle and double arrows; Larkum, 2013). The integration zones receive distinct inputs arriving from basal feedforward (FF, orange) and apical lateral and feedback (FB, blue) pathways. The pyramidal cell dynamics are further regulated by a local network of inhibitory PV, SOM, and VIP interneurons (gray) that is driven by the FF and FB input and by the cell’s output (black) (Niell and Scanziani, 2021; Zhang et al., 2014; Campagnola et al., 2022), which also feeds back onto the basal compartment. (B) Backpropagation-activated calcium spike (BAC) firing in thick-tufted pyramidal cells (Larkum et al., 1999; Larkum, 2013). Firing rate enhancement from apical-basal coincidence is generated by a phasic causal chain of four steps. The steps are numbered on the cell (left) and on the time axis of the membrane potentials of tuft, Ca^2+^ initiation zone, and soma (right). Dotted lines delimit the steps in time. All arrows point in the direction in which the signals propagate, and the FF and FB markers on the traces denote input onset. First, FF input evokes an axonal Na^+^ action potential (AP). Second, the AP actively back-propagates into the apical dendrite (bAP, orange arrows), while FB input depolarizes the tuft, and this depolarization spreads to the Ca^2+^ initiation zone (blue arrows). Third, if both coincide within a few milliseconds, their summed depolarization (orange and blue areas) at the Ca^2+^ initiation zone crosses the threshold (thr.) of a voltage-gated calcium spike (green area), which the bAP alone does not reach. Fourth, the calcium spike spreads to the soma (green arrows) and produces an axonal spike burst, effectively increasing the cell’s firing rate. (C) Populations of pyramidal cells in cortex cooperate by different routes, as illustrated conceptually for figure-ground segregation across the visual areas V1, V2, and V4 and the frontal eye field (FEF). The stimulus contains a texture-defined figure, and dashed circles mark the receptive fields of neurons at its center (*a*), at its boundary (*b*), and in the background (*c*). These neurons integrate their input according to their feature tuning (symbols within circles). During the initial feedforward sweep (*t*_1_), driving excitatory routes (orange) carry signals forward across areas, and all neurons whose preferred feature is present respond without amplification. During subsequent recurrent processing (*t*_2_), long-range feedback projections from V2 and V4 and task-related feedback from FEF for a saccade to the figure provide modulatory contextual information (blue flat heads, denoting modulation). Those neurons in which forward signals and context coincide become amplified (green) and form an assembly that labels the figure and expresses a state of binding (Roelfsema, 2023), while background neurons remain unamplified. Brain outlines in (C) adapted from Chilton (2020) under CC BY 4.0 with added area markers and connections.

At the *network level*, pyramidal cells form intricate connectivity patterns within and across brain regions. Locally, the L5Pyr cells are regulated by a complex network of inhibitory interneurons (Fig. 1A), whose types are found to target distinct anatomical sites of the cell. The most relevant interneuron types are parvalbumin- (PV), somatostatin- (SOM), vasoactive intestinal polypeptide- (VIP), and neuron-derived neurotrophic factor-expressing (NDNF) interneurons (Tremblay et al., 2016; Schuman et al., 2019). Understanding their exact circuit motifs and their impact on L5Pyr cells is an active field of investigation. Multiple such circuit motifs have been proposed (Wang et al., 2004; Adesnik et al., 2012; Pfeffer et al., 2013; Pi et al., 2013; Harris and Mrsic-Flogel, 2013; Hangya et al., 2014; Bortone et al., 2014; Tremblay et al., 2016; Yang et al., 2016; Kirchberger et al., 2021; Niell and Scanziani, 2021; Campagnola et al., 2022; Wagatsuma et al., 2025). While partially differing interneuron couplings have been suggested, common connectivity and interaction patterns have also been identified. Examples are PV cells targeting L5Pyr cells perisomatically, SOM cells inhibiting the proximal and distal apical tuft, and VIP cells dis-inhibiting and NDNF cells inhibiting apical integration (Tremblay et al., 2016; Schuman et al., 2019; Kirchberger et al., 2021; Campagnola et al., 2022; Naumann et al., 2025). In addition to their local interneuron connections, L5Pyr cells form excitatory intra-area projection patterns with other pyramidal cells across layers of the laminar structure of cortex, and inter-area projection patterns with other cortical (Gilbert and Li, 2013) and sub-cortical areas, such as the thalamus (Halassa and Kastner, 2017; Halassa, 2022). These intra- and inter-area motifs give rise to functional hierarchies and recurrent feedforward-feedback (FF-FB) processing between populations of L5Pyr cells and excitatory-excitatory pyramidal and thalamic cell coupling. A first feedforward sweep establishes the feature selectivity of neurons, and recurrent processing binds those representing a common object into an assembly of enhanced firing rates (Lamme and Roelfsema, 2000; Roelfsema, 2023; Fig. 1C). The engaged L5Pyr cell populations are thought to yield joint but decentralized brain-wide states of information processing. These brain-wide states would form the relevant substrate for understanding cognition (Roelfsema, 2023) and even consciousness (Aru et al., 2019; Suzuki and Larkum, 2020; Storm et al., 2024).

In this paper, we propose a dynamical TPN model of L5Pyr cell populations and their local circuit of PV, SOM, and VIP interneurons that contributes to the understanding of cell- and network-level processes alike (Sects. 2, 3). Its generic processing motifs, asymmetric FF-FB integration and pooled inhibition, implement canonical operations, such as biased competition (Desimone and Duncan, 1995), and allow for application-dependent model simplifications. The work makes four contributions. First, the TPN with bidirectional apical-basal coupling and a voltage-dependent apical gate reproduces the asymmetric basal-apical input-output map of a multi-compartment L5Pyr model (Shai et al., 2015; Hay et al., 2011) and shows how apical amplification unfolds over time under interneuron control (Sect. 4.1). Second, the apical-to-basal current that amplifies the TPN also tags it for a local four-factor learning rule driven by reward. This coincidence tag establishes structural credit assignment across the network’s neurons, and VIP dis-inhibition gates whether learning takes place (Sect. 4.2). Third, networks of cooperating TPN populations with lateral apical coupling perform evidence integration and evidence propagation to arrive at coherent network states, which form a basis for perceptual integration (Sect. 5.1). In a model of the visual aperture problem (Pack and Born, 2001), disambiguating motion signals propagate at constant speed and fail to propagate when the basal and apical compartments of each TPN are decoupled (Sect. 5.2, Fig. 8). Fourth, a reduction relates the TPN to the modulated excitatory-inhibitory (Mod-E-I) point neurons (Brosch and Neumann, 2014a) and the simplified TPN of our earlier large-scale models of visual motion integration (Bayerl and Neumann, 2004; Bouecke et al., 2011) and incremental grouping (Schmid and Neumann, 2025). These models captured the asymmetric FF-FB integration by a modulatory interaction of the form FF · (1 + FB), which can also be derived from a firing-rate circuit with feedback-recruited dis-inhibition fitted to recordings of L5Pyr cells (Brosch and Neumann, 2014b). The reduction shows which properties of the TPN this form retains and which it loses (Appendix E). There, we also summarize published results of the simplified TPN on incremental grouping (Schmid and Neumann, 2025). Processing and learning are thus studied in the full TPN, and the reduced models carry its principles to large-scale simulations. We conclude with a discussion of the broader context of the model at the cell and network level, its relation to other approaches, and its testable predictions (Sect. 6).

## 2. Methods – Two-point neuron model for contextual processing and learning

Here we present a two-point neuron model that aims to capture the computational mechanisms for integrating driving sensory evidence with contextual signals during both processing and learning. The model TPN links the cell level to the network level and proposes a functional role of this integration mechanism. At the *cell level*, the proposed TPN model relates the inner working mechanisms of pyramidal cells and their interaction with circuits of local inhibitory interneurons to biophysical evidence. We develop the model within a framework that focuses on the components central to this integration mechanism and that links it with earlier work on simplified Mod-E-I models (Bayerl and Neumann, 2004; Bouecke et al., 2011; Brosch and Neumann, 2014a; Schmid and Neumann, 2025). The framework describes how local processing and connectivity give rise to motifs of asymmetric FF-FB integration and inhibitory regulation of the TPN’s operating regime, and how the same integration mechanism provides the signal for learning.

### 2.1. Processing dynamics in the two-point neuron model

The presented TPN model is a phenomenological rate-based model that subsumes a group of spiking pyramidal neurons and their processing dynamics. It is formulated for thick-tufted layer V pyramidal (L5Pyr) cells. Each TPN is described by two conductance-based compartments that integrate signals arriving at different anatomical sites of L5Pyr cells (Fig. 2A). A basal compartment aggregates perisomatic input and provides the signal for firing rate computation, while an apical compartment aggregates input to distal apical dendrites and the apical tuft. Inputs arriving at the two compartments play distinct roles. The basal compartment receives driving input from feedforward (FF) projections, while the apical compartment receives modulating contextual input from lateral or feedback (FB) and other inter-areal projections. Both compartments are coupled with each other to perform coincidence detection. In the model, this coincidence detection mechanism operates asymmetrically: basal FF input can drive the cell into firing on its own, while apical FB input can only amplify existing basal activity upon coincidence. The mechanism is modeled to reproduce the rate-level consequences of the apical-basal coupling that is mediated by backpropagation-activated calcium spike (BAC) firing in L5Pyr cells (Larkum et al., 1999; Larkum, 2013; cf. Fig. 1B), without modeling the underlying voltage-gated calcium channels, action potentials, and bursts separately. The basal and apical patterns that yield coincidence themselves need to be learned. In the presented model, this local synaptic learning process is likewise dependent on the same signaling logic.

**Figure 2:**
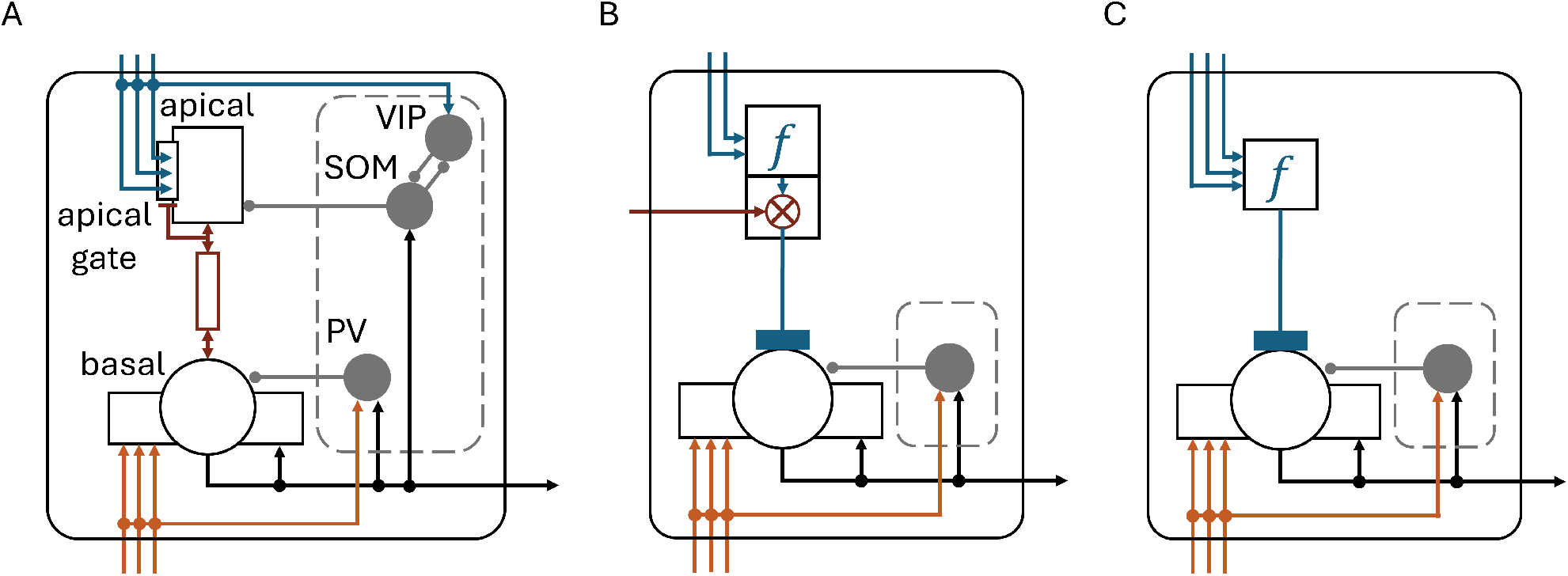
Two-point neuron model and simplified variants. (A) Proposed TPN model. The model specifies a pyramidal cell with a basal compartment (circle with perisomatic dendrites) and an apical compartment (rectangle) together with a local pool of PV, SOM, and VIP interneurons (gray discs within the dashed frame). The two compartments exchange current through the dendritic trunk (dark red resistor symbol with double arrows, Eq. 8). Feedforward (FF) input (orange) projects onto the basal compartment and onto PV cells, whereas feedback (FB) input (blue) projects onto the apical compartment and onto VIP cells. The FB input reaches the apical compartment through a voltage-dependent apical gate (Eq. 6), which opens only once the apical compartment has been depolarized through the exchange with the basal compartment (dark red line). The pyramidal cell’s output (black) projects recurrently to its own basal compartment, to PV and SOM cells, and to other pyramidal cells. Arrowheads denote excitatory and round heads inhibitory connections (cf. Tab. 1). PV cells inhibit the basal compartment, while SOM cells inhibit the apical compartment and, thus, regulate the apical integration. VIP cells, in turn, inhibit SOM cells, which yields a dis-inhibition of the apical integration. Conversely, SOM cells also inhibit VIP cells. (B,C) Simplified circuit variants used in earlier works that are contextualized within the current framework. Both models contain the main processing motifs of the TPN model, but lead to reduced neuronal complexity with a single type of inhibitory cells driven by FF input and the cell’s output. Earlier descriptions utilized point neurons with a reduction to modulated excitatory-inhibitory (Mod-E-I) interaction of the form FF · (1 + *f* (FB)) with a modulation function *f* (box in C; Bayerl and Neumann, 2004; Bouecke et al., 2011; Brosch and Neumann, 2014a), while a more recent version extended the Mod-E-I description to a TPN with external gating of apical input (dark red arrow onto the multiplication node *⊗*; B; Schmid and Neumann, 2025). The flat-headed link of the simplified variants maps a scalar unidirectionally from the apical input onto the basal compartment, unlike the dynamical bidirectional coupling of the full TPN model. As for the contextual feedback in Fig. 1C, flat heads denote modulatory influences. The reduction of the TPN model to these variants is derived in Appendix E.

**Table 1:** Connectivity matrix between components of the two-point neuron (TPN) model. Rows represent source elements (*from, j*), columns represent target elements (*to, i*), and entries list the channel type *s* by which elements are connected, namely excitatory (**Exc**), inhibitory (**Inh**), or compartmental exchange (**Comp**). Rows and columns **b** and **a** denote the basal and the apical compartment, with the TPN output rate as the source of the excitatory projections of b.

|  |  | To |  |  |  |  |
| --- | --- | --- | --- | --- | --- | --- |
|  |  | b | a | PV | SOM | VIP |
| From | **FF** | Exc | – | Exc | – | – |
|  | **FB** | – | Exc | – | – | Exc |
|  | **b** | Exc | Comp | Exc | Exc | – |
|  | **a** | Comp | – | – | – | – |
|  | **PV** | Inh | – | – | – | – |
|  | **SOM** | – | Inh | – | – | Inh |
|  | **VIP** | – | – | – | Inh | – |

The TPN is embedded within a local circuitry of inhibitory interneurons, which further regulate the TPN’s activity and integration behavior during processing and learning. The inhibitory interneurons are modeled by single compartments and are categorized into three different types, namely, PV, SOM, and VIP cells. Each cell type follows specific connectivity patterns and receives its inputs from different sources (Tab. 1). The TPN model synthesizes different sources of evidence, as circuit motifs of inhibitory interneurons are still an active field of research (cf. Sect. 1; Wang et al., 2004; Adesnik et al., 2012; Pfeffer et al., 2013; Pi et al., 2013; Harris and Mrsic-Flogel, 2013; Hangya et al., 2014; Bortone et al., 2014; Tremblay et al., 2016; Yang et al., 2016; Kirchberger et al., 2021; Niell and Scanziani, 2021; Campagnola et al., 2022). PV interneurons receive their input from FF sources and recurrent output from the TPN and target the basal compartment inhibitorily (Niell and Scanziani, 2021; Williams and Holtmaat, 2019; Campagnola et al., 2022). SOM interneurons receive their input from recurrent TPN output and project back onto the apical compartment. As a result, they regulate the apical integration and, thus, apical-basal coupling (Kirchberger et al., 2021; Murayama et al., 2009; Roelfsema and Holtmaat, 2018). VIP interneurons, in turn, receive their input from FB sources and inhibit SOM interneurons (Zhang et al., 2014; Williams and Holtmaat, 2019). Together with SOM interneurons, this yields a dis-inhibition of the apical integration (Pi et al., 2013; Kirchberger et al., 2021; Roelfsema and Holtmaat, 2018). Conversely, SOM interneurons also inhibit VIP interneurons (Pfeffer et al., 2013; Niell and Scanziani, 2021).

#### 2.1.1. Basic processing motifs and mechanisms

The presented model implements a distinct set of processing motifs to form the basis of its function. The set of motifs stems from a framework of generic neural processing principles (Schmid et al., 2023) and is formed by feature extraction, feedback modulation, (dis-)inhibition and pool competition, and nonlinear firing rate computation. Here we contextualize the proposed TPN model within this framework and describe the relation to each of the motifs.

##### Feature extraction

Neurons exhibit differing basic tuning properties toward features of their upstream input. These basic tuning properties are attributed to the afferent synaptic connections and can be conceptualized by the connectivity pattern and their strength. In the model, these basic tuning properties are encoded in the weight vector, or kernel, of a cell’s feature selectivity. Any synaptic input is weighted and summed, which is expressed by a local dot product of pattern matching^1^.

##### Activity modulation by feedback and other inter-areal signals

At the heart of the TPN model lies the coincidence detection mechanism described above. It leads to an asymmetric combination of basal FF and apical contextual FB signals. As a result, FF input can drive the TPN into firing at moderately growing output rates for growing inputs, while coinciding FF and FB input can gain-modulate the TPN into a regime of overall higher firing rates, also for weaker FF inputs (cf. Fig. 1B). At an abstract level, this asymmetric FF-FB integration can be described by an equation of the form FF · (1 + FB), which has been used in earlier modeling investigations (Bayerl and Neumann, 2004; Bouecke et al., 2011; Brosch and Neumann, 2014a) and was shown to be derivable from a dis-inhibitory firing-rate circuit that accounts for the firing rates of L5Pyr cells (Brosch and Neumann, 2014b). Here, we propose a TPN model with an apical compartment and explicit PV, SOM, and VIP cells to implement dynamical phenomena of this asymmetric FF-FB integration mechanism. It combines a symmetric, conductance-based exchange between the basal and apical compartments with a voltage-dependent gating of apical FB input to reproduce the causal ordering of BAC firing at the level of firing rates (Larkum et al., 1999; Larkum, 2013; cf. Sect. 1 and Fig. 1B).

##### (Dis-)inhibition & pool competition

The asymmetric FF-FB mechanism is paired with a local circuit motif between TPNs and inhibitory neurons forming inhibitory as well as mutual excitatory-inhibitory and inhibitory-inhibitory circuit interactions. These mainly serve two functions: (i) to control the operating regime and state of individual TPNs, and (ii) to put different TPNs into competition with each other. *Controlling the operating regime* is achieved by local projection patterns between the TPN and its associated PV, SOM, and VIP interneurons (see projection patterns described above; Fig. 2A). PV neurons keep the overall TPN activity within an operating regime, as inhibition from PV cells grows with FF input and recurrent TPN output. The interaction between SOM and VIP cells gates the apical-basal integration and depends on FB input and the recurrent TPN output. *Competition* between different TPNs is achieved by pooling activity from multiple TPNs to form an additional component to the inhibitory signaling. This pool competition puts the activities of different TPNs into relation with each other, and, thus, normalizes their overall activity. Within the TPN model, this pooling is performed by PV cells. Additionally, a secondary motif of competition is established within the TPN model. FB processes compete for apical-basal integration through SOM cells that inhibit VIP cells of the associated TPN as well as of other competing TPNs.

##### Nonlinear firing rate computation

The somatic output response of TPNs communicates their current state to other neurons. A transfer function is applied to map the basal membrane potential to a firing rate representing the average state of the local group of spiking neurons. The transfer function is nonlinear such that membrane potentials exceeding a threshold generate monotonically increasing firing rates.

These basic processing motifs have also been used in predecessors to the TPN model presented here. The earlier model descriptions were based on point neurons that reduced the local neural processing to Mod-E-I interaction (Fig. 2C; Bayerl and Neumann, 2004; Bouecke et al., 2011). The most recent model then extended the Mod-E-I description to that of a TPN with external gating of apical input while retaining a single type of inhibitory cells (Fig. 2B; Schmid and Neumann, 2025; see Appendix E for further details).

The model TPN and its local circuitry of inhibitory neurons are described by a system of first-order ordinary differential equations (ODEs). These equations capture the dynamical computational properties for simultaneous learning and processing in the model.

#### 2.1.2. Equations

The system of ODEs consists of five state variables that describe the temporal evolution of the membrane potentials of the TPN’s basal and apical compartments (*v*_*b*_ and *v*_*a*_) and of the local inhibitory interneurons (*v*_*PV*_ , *v*_*S OM*_, *v*_*VIP*_). The rate of change 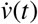 of individual cell types or cell compartments is defined by time constants *τ*_*i*_. *Excitatory pyramidal cells* with separate basal and apical compartments are described by

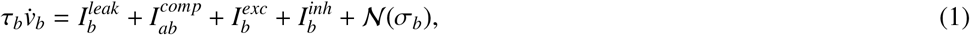

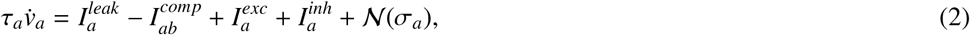

and *inhibitory interneurons* (PV, SOM, VIP cells) are modeled as single-compartment units

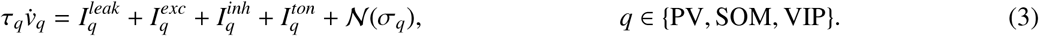

Input currents 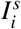 capture the effective impact on the membrane potential from different groups of ionic channels, namely excitatory and inhibitory synaptic input (*exc, inh*), leakage currents (*leak*), and ionic exchange between cell compartments (*comp*). SOM and VIP cells receive an additional tonic input 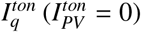. Optional additive Gaussian white noise *N*(*σ*_*i*_) is applied to each state variable to account for neural stochasticity. Since the state variables describe the mean activity of a local group of neurons, the time constants are those of rate units. The basal compartment relaxes with *τ*_*b*_ = 10 ms, on the order of the response times of cortical neurons (Koch et al., 1996), while the apical compartment integrates its input more slowly with *τ*_*a*_ = 35 ms, in line with NMDA spikes in tuft dendrites (Larkum et al., 2009) that last about 50 to 100 ms (Antic et al., 2010), and the interneurons follow their input faster than both compartments with *τ*_*q*_ = 1 ms, close to the 2 ms used in rate models of the PV-SOM-VIP circuit (Hertäg and Sprekeler, 2020). All times are therefore given in milliseconds (Tab. A.1 and Appendix A.1).

The input currents to the apical and basal TPN compartments follow a conductance-based description

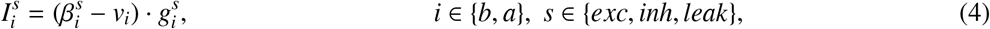

with an input-dependent conductance 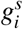 and a force term 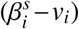 that shunts off the input’s impact once the membrane potential *v*_*i*_ equilibrates to the channel’s constant reversal potential 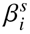^2^.

The conductances of the synaptic inputs (*exc, inh*) of all cells are described by weighting the firing rate *r*_*j*_ of projecting cells *j* with a synaptic efficacy 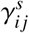.

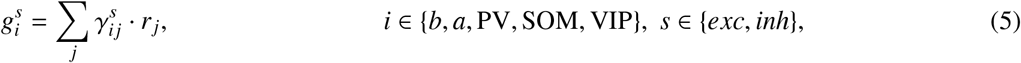

while the leak conductances 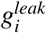 are constants. External FF and FB inputs enter as rectified sums 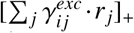, which keep the conductances non-negative, with [*x*]_+_ = max(*x*, 0). The connectivity of an input node *j* that projects to each channel *s* of a node *i* is structured according to Tab. 1 (cf. Fig. 2A).

The synaptic efficacies 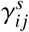 are scalar weights with the exception of the excitatory apical input. Its efficacy captures a *voltage-dependent gating* of apical input by

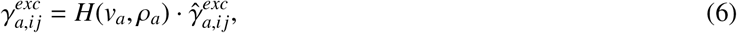

which gates the maximally possible synaptic efficacy 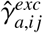 with a voltage-dependent Heaviside function *H*(*v*_*a*_, *ρ*_*a*_)

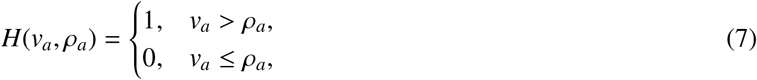

with constant threshold *ρ*_*a*_. This apical gate is a single threshold nonlinearity, which is motivated by NMDA-receptor-mediated tuft spikes (Larkum et al., 2009) and represents an equivalent threshold of the dendritic calcium spike in BAC firing (Larkum et al., 1999). As voltage-gated calcium channels are not modeled separately, the gate captures the consequences of BAC firing at the level of firing rates rather than its millisecond coincidence window, calcium-spike kinetics, or bursts (see Appendix A.3 for a variant with sodium- and calcium-like exchange terms).

The conductance-based equation for the *exchange current* between the basal and apical compartments is defined by a conductance-gated potential difference between the two compartments, *v*_*b*_ and *v*_*a*_,

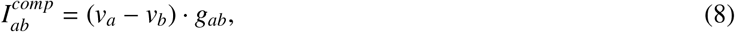

where the conductance *g*_*ab*_ is constant, and positive values of 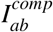 denote a current from the apical to the basal compartment. This exchange current, together with the voltage-dependent gating of apical input above, forms the basis of the model’s apical-basal coincidence detection mechanism and captures the computational logic underlying BAC firing dynamics at a phenomenological level (cf. Fig. 1B; see Appendix A.3 for an extended model variant and discussion).

The input currents to the inhibitory interneurons are modeled by simpler equations with static synaptic efficacies, in which excitatory inputs are current-based without a force term, while leak and inhibitory inputs shunt the membrane potential toward the reversal potential 0, i.e.,

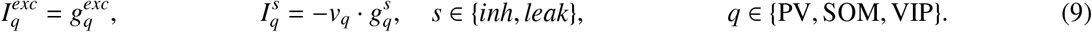

The tonic input 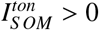 sets the resting activity of SOM cells, and 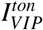 shifts the balance of the VIP-SOM interaction.

The rate-based output *r*_*i*_ of each cell is a nonlinear mapping of its membrane potential to a firing rate. In the presented model, this mapping is described by threshold-linear functions

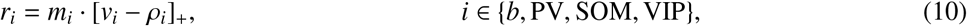

with a threshold *ρ*_*i*_ and a scaling factor *m*_*i*_, where *r*_*b*_ is the output rate of the TPN.

### 2.2. Learning in the two-point neuron model

#### 2.2.1. Learning objectives

The described apical-basal coincidence detection mechanism signals a match between basal driving sensory evidence and apical context. Which patterns are extracted at the basal and apical compartments, and that they belong together, is determined by synaptic long-term adaptation, i.e., learning. In turn, apical-basal coincidence itself is a driver for learning (Kampa et al., 2006; Takahashi and Magee, 2009; Gambino et al., 2014; Bittner et al., 2017; Williams and Holtmaat, 2019; Wilmerding et al., 2026). The presented model captures how apical-basal coincidence detection is involved in learning and how it depends on the same signals used during processing (Fig. 3A).

**Figure 3:**
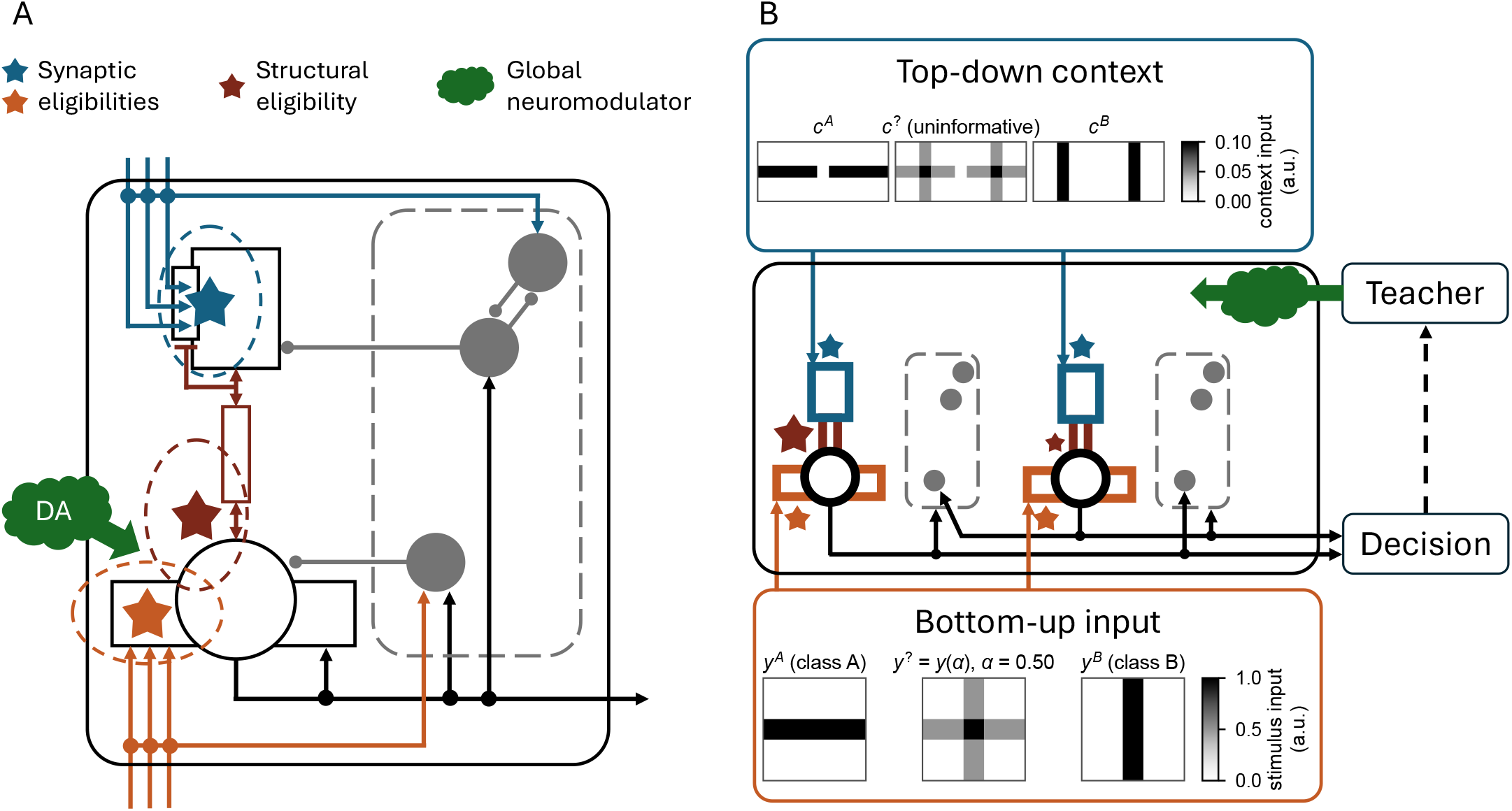
Learning to discriminate orientations in a pool of two-point neurons (TPNs). (A) Learning factors in the TPN microcircuit. The TPN and its PV, SOM, and VIP cells (gray discs within the dashed frame) are drawn as in Fig. 2A. The basal learning rule combines synaptic eligibilities at the basal synapses (orange star) with a coincidence tag based on the occurrence of apical-basal coincidence (dark red star) and with a global neuromodulatory signal (dopamine, DA). The coincidence tag establishes the structural eligibility of the TPN for learning. The apical learning rule, in contrast, is purely local and relies on the synaptic eligibilities at the apical synapses (blue star). (B) Setup of the orientation discrimination task. Two TPNs, each with its own PV, SOM, and VIP cells, compete via pooled inhibition through their PV cells for the representation of sensory evidence. Oriented contrast stimuli of 5 × 5 pixels are presented as basal input (orange) and show either a horizontal bar *y*^*A*^, a vertical bar *y*^*B*^, or a mixture *y*^?^ of both (bottom). Flanker stimuli in a left and a right patch are presented as apical context (blue) and show either horizontal bars *c*^*A*^, vertical bars *c*^*B*^, or their uninformative superposition *c*^?^ (top). The mixture *y*^?^ is shown for *α* = 0.50, the point of subjective equivalence of the trained networks. The output rates of both neurons are fed into a decision function. The decision function provides the discrimination response to a teacher module, which releases a positive or negative reward based on the correctness of the response. The teacher’s reward signal is delivered as volume-transmitted DA, and its uptake by each TPN is set by the strength of its coincidence tag (differently sized dark red stars). See Sect. 2.3.2 and Appendix C.2 for details.

##### Overall goal – participation in coherent representations

Basal and apical synapses are trained by separate specific objectives. Yet, both help the TPN to achieve one overarching goal best described as “maximize the chance of contributing to a coherent representation at the network level”. The learning objectives for basal and apical synaptic plasticity in the TPN model follow local learning rules.

##### Basal objective – extract relevant input patterns

Basal synapses extract useful evidence about driving upstream sensory input. The basal learning objective should therefore help to extract patterns that are relevant for behavioral task performance. Such pattern selectivity helps the neuron to become part of a coherent representation in the network. The basal objective therefore needs to learn about *driving input patterns*, but in combination with their *utility* for participating in the global network.

##### Apical objective – find co-active context patterns

Apical synapses are fed by a broad range of lateral, FB and other inter-areal signals and need to tell apart relevant context that typically coincides with basal activity patterns from nuisance signals. The apical learning objective needs to learn which of the different arriving context signals coincide and, thus, correlate with the TPN’s own representation. Matching these contextual patterns will let the TPN reach an amplified activity state from coincidence during processing. This, in turn, helps again with participating in a coherent representation at the network level. The apical objective therefore is designed to learn about *context co-occurrences with the neuron’s own state*.

##### Processing interacts with learning locally – coincidence and local competition

The TPNs form pools locally competing for representation of the sensory evidence at each location (see Sect. 2.1.1). Thus, if bottom-up input from a sensory location is ambiguous, it could be treated as evidence toward several competing alternatives represented by different neurons within the pool. The neuron that integrates the input better with the context than the rest gains an advantage in this competition for participating in a coherent representation across locations. Processing, thus, yields three consequences for learning, namely (i) different neurons within a pool should specialize in extracting different patterns to sometimes win the competition, (ii) the same apical-basal coincidence mechanism that helps the TPN to reach amplified levels of activity is also a useful learning signal indicating the *utility* of the neuron, and (iii) learning is further regulated by the surrounding circuit of interneurons via VIP-SOM (dis-)inhibition through their gating of apical depolarization.

#### 2.2.2. Equations

An additional set of ODEs captures the dynamics of basal and apical synaptic plasticity in the model. These equations describe local learning rules and are driven by the model quantities described above. They adapt the FF and FB efficacies of a pool of TPNs, in which the presynaptic rates *r*_*j*_ are the pixel values of stimulus and context (Fig. 3B; Appendix C.2).

##### Learning in the basal compartment

The basal synaptic plasticity implements a four-factor rule (Frémaux and Gerstner, 2016; Gerstner et al., 2018; Roelfsema and van Ooyen, 2005; Roelfsema and Holtmaat, 2018) of the form

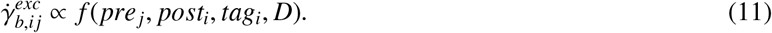

The four-factor rule on the right-hand side is defined by pre- and postsynaptic synapse-specific factors *pre*_*j*_, *post*_*i*_ between neurons *j* and *i*, a neuron-specific structural credit factor *tag*_*i*_ for the postsynaptic neuron, and a scalar global neuromodulator-derived factor *D* (Fig. 3A). Hebb-like synapse-specific factors allow for specificity and pattern formation, the neuron-specific factor enables structural credit assignment for contribution to the network’s computation, and the global scalar factor steers the direction of adaptation. Concretely, we choose an eligibility-based description for the factors (Izhikevich, 2007; Gerstner et al., 2018), which allows for temporal integration of adaptation signals from a stream of ongoing computation

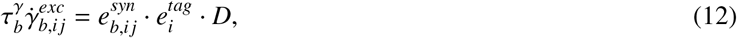

with eligibility terms 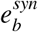 combining pre- and postsynaptic factors, *e*^*tag*^ for the structural credit factor and time constant 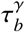

The *synapse-specific eligibility* is given by an instar rule (Grossberg, 1976)

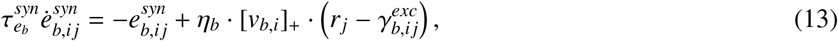

with presynaptic firing rate *r*_*j*_ and learning rate *η*_*b*_. As postsynaptic factor, the rule reads the rectified basal potential [*v*_*b,i*_]_+_ instead of the output rate *r*_*b,i*_ (Eq. 10).

The neuron-specific eligibility to adapt the local synaptic weights forms a *coincidence tag* based on apical-basal coincidence detection

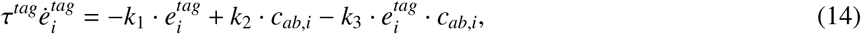

with constants *k*_1_, *k*_2_, *k*_3_ parametrizing the speed and decay of the integration and its saturation level, and with the apical-basal coincidence term

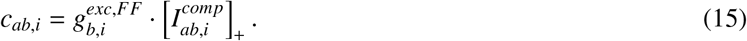

Tag formation for structural credit assignment happens if basal feedforward input 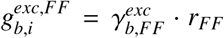(cf. Eq. 5) meets an apical-to-basal current via the rectified exchange term 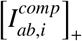 . Because 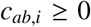, the tag stays between 0 and *k*_2_*/k*_3_ (see Fig. C.1 in the Appendix).

The *coincidence tag* steers the neuron’s uptake of global volume-transmitted neuromodulators and enables learning. A neuron without coincidence forms no tag, so that its basal efficacies do not change. The coincidence tag thereby establishes the structural eligibility of each neuron for learning, i.e., a structural credit assignment across the network’s neurons.

The *neuromodulatory* factor *D* derives from the amount of released neuromodulators, which is determined by specific structures in the network, such as the cortico-basal ganglia loop (Brown et al., 1999). It is modeled as a dopamine-like reward signal (Schultz et al., 1997), whose level *D* accumulates the dopamine release *D*^*\**^(*t*) of a trial at time point *t*,

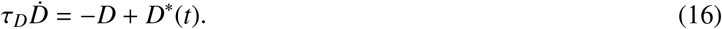

The release is positive for correct and negative for incorrect decisions (Appendix C.2).

##### Learning in the apical compartment

The apical synaptic plasticity is given by a purely local stabilized covariance rule of the form *f* (*pre*_*j*_, *post*_*i*_), defined by

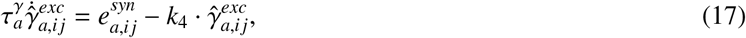

with a decay term with constant *k*_4_ that stabilizes the efficacies, and with the apical synaptic eligibility

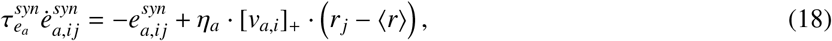

with learning rate *η*_*a*_ and the rectified apical potential [*v*_*a,i*_]_+_ as postsynaptic factor. The term *r*_*j*_ − ⟨*r*⟩ expresses the synapse-specific contribution of input *r*_*j*_ to the apical dendritic tree relative to the average input across all the neuron’s apical synapses *⟨r⟩*.

The efficacies are not bounded and can become negative, which can be read as net disynaptic inhibition, while the rectified input sums keep the conductances non-negative (Sect. 2.1.2). The parameters of the learning rules are listed in Tab. C.1.

### 2.3. Comparison models, evaluation and metrics

This section summarizes how the TPN is compared with a multi-compartment neuron model at the level of single cells (Fig. 5) and how the learned pool of TPNs is evaluated. Further details are given in Appendices B.1 and C.2.

**Figure 4:**
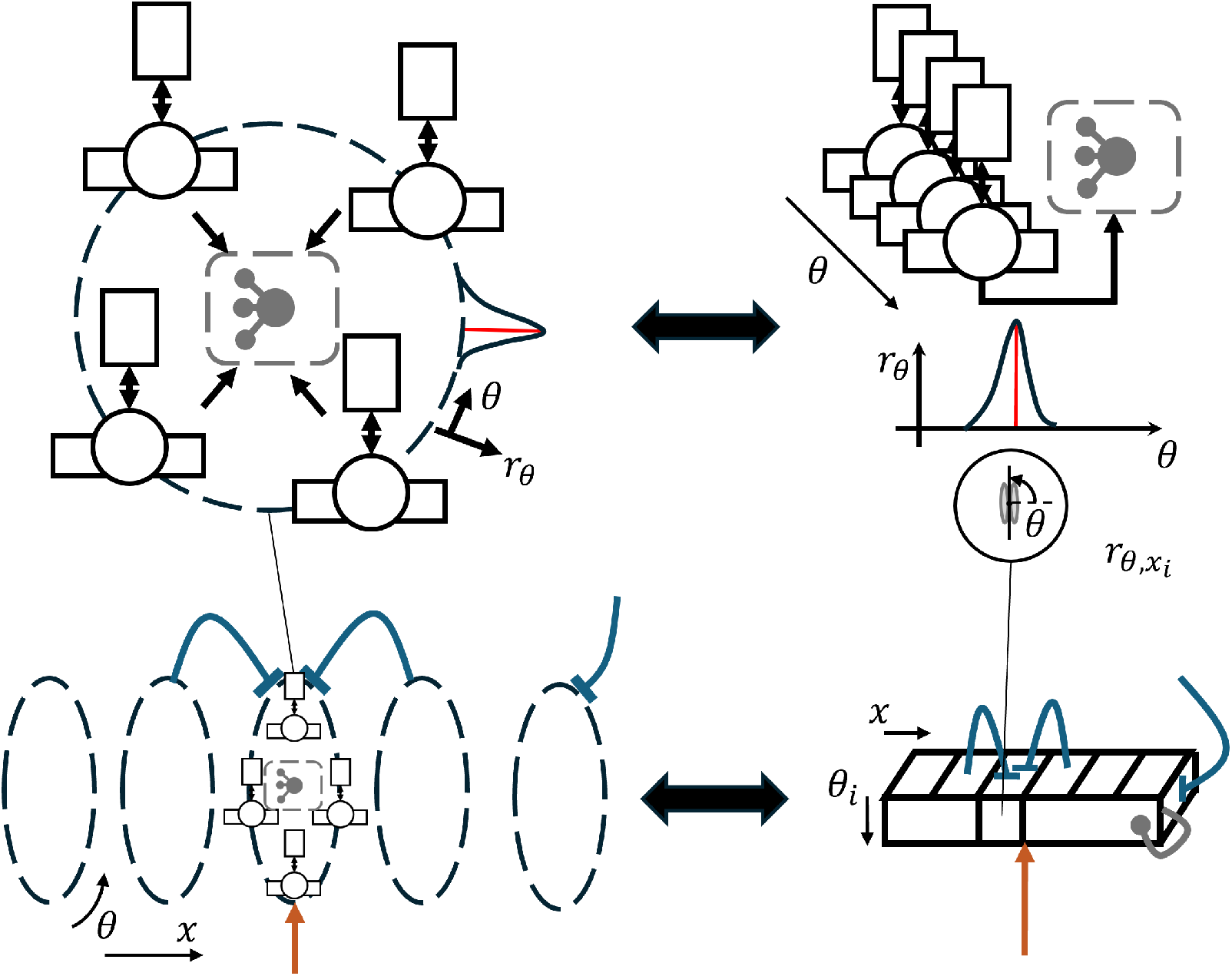
Canonical network model of cooperating TPNs. (Top, left) The TPNs of a local population, drawn with their basal (circle with horizontal bar) and apical (rectangle) compartments, are tuned to different values *θ* within a 1-D feature domain. As in ring attractor networks, they are arranged on a ring (dashed circle) according to their distance within the feature dimension. TPN signals are pooled by inhibitory PV cells (gray) to yield competition within the population. The curve along the ring sketches the resulting rate profile *r*_*θ*_ with its peak (red line). (Top, right) Equivalently, the ring can be understood as a population of TPNs linearly organized along *θ* with circular boundary conditions. (Bottom, left) Within a spatially organized network, such rings encode the feature values at different locations along the spatial dimension *x*. FF input (orange arrow) drives the basal compartments. Between neighboring rings, contextual projections (blue lines) reach the apical compartments (and the VIP cells, not drawn) of TPNs with the same preferred feature value and provide a means of cooperation. Their flat heads indicate that this context modulates the TPNs without driving them. The line at the right border denotes external FB from outside the network. (Bottom, right) Likewise, the spatially coupled rings can be interpreted as a neural network layer with a spatial and a feature dimension. Its TPN at location *x*_*k*_ with preferred value *θ*_*n*_ has the rate *r*(*x*_*k*_, *θ*_*n*_), labeled 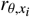, and the inset depicts *θ* as the orientation of a bar. The gray loop with a round head indicates the inhibitory interactions along *θ* within a location. For clarity, only four TPNs are drawn per ring, whereas the simulated rings consist of *N* = 200 TPNs.

**Figure 5:**
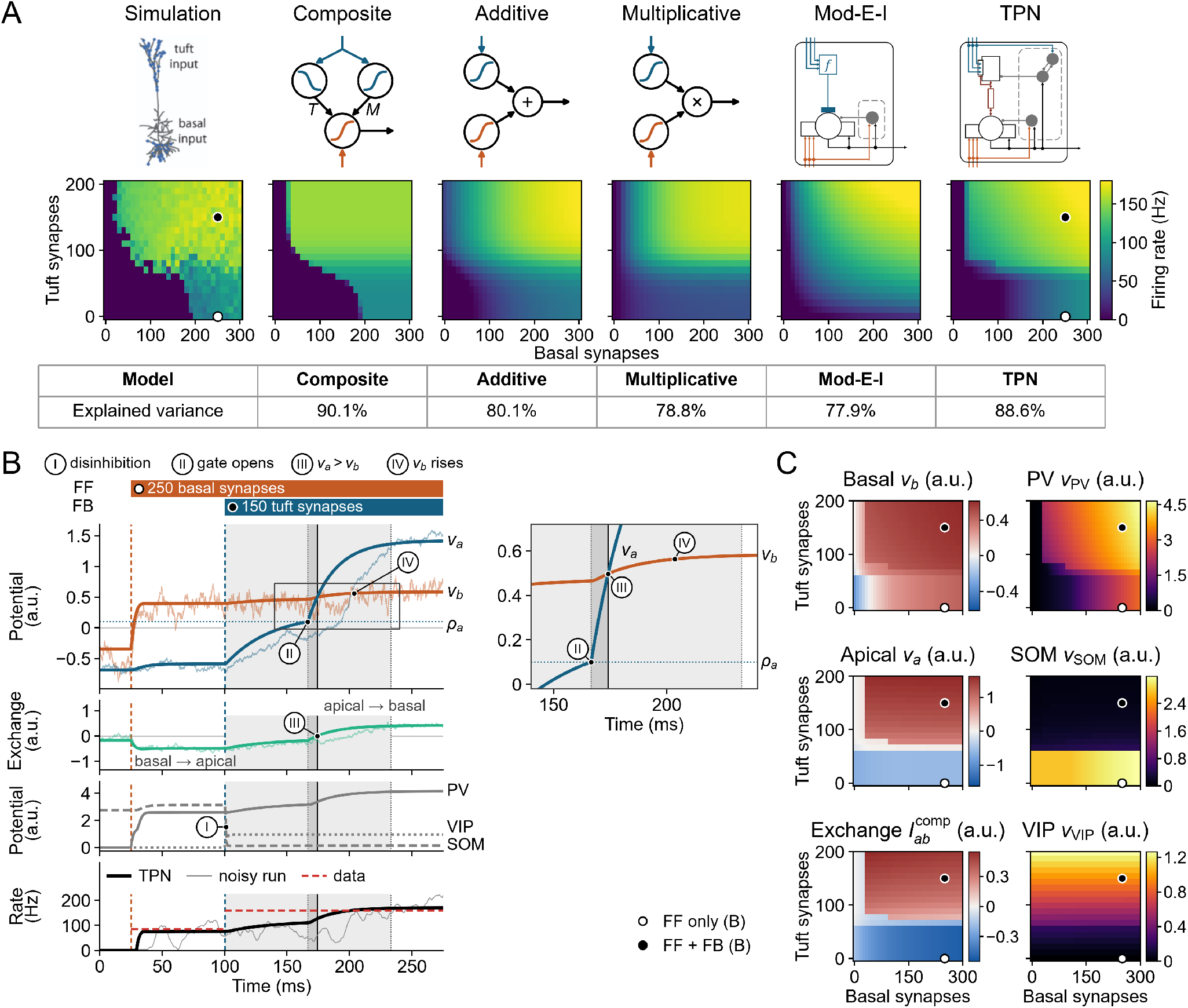
Pyramidal cell response and TPN model dynamics. (A) Asymmetric feedforward-feedback computation in pyramidal cell models. Models of varying detail for describing pyramidal cell integration (first row) can be grouped into a biophysical multi-compartment model (left), abstract functional descriptions (middle), and dynamical models (right). The reference data in the left column stem from the *multi-compartment model* of Shai et al. (2015), which receives input through tuft and basal synapses (blue dots). In the *composite model* of Shai et al., the tuft input (blue) sets the threshold *T* and the maximum *M* of a sigmoidal function of the basal input (orange). The *additive* and the *multiplicative model*, in turn, add or multiply a sigmoidal function of the tuft input and one of the basal input. The *Mod-E-I model* is the steady state of the TPN reduced to a point neuron (Fig. 2C), and the *TPN model* proposed here gives the TPN output at equilibrium under constant inputs (Fig. 2A). The firing rates of all models (second row, color scale in Hz) are shown over the number of active basal and tuft synapses in steps of 10 (651 input conditions). Conditions in which the multi-compartment model fires no or a single spike, as well as negative predictions of the fitted models, are shown at 0 Hz. All other models were (re)fitted to these conditions by least squares (Sect. 2.3.1), and the table gives their explained variance *R*^2^ (third row). Open and filled dots on the maps of the multi-compartment model and the TPN model mark the two input conditions of panel B. (B) TPN model dynamics. The TPN receives a step of FF input at 25 ms (250 basal synapses, *r*_*FF*_ = 1.63), followed by a step of FB input at 100 ms (150 tuft synapses, *r*_*FB*_ = 0.72), mapped by the TPN fit of panel A. From top to bottom, the rows show the input timing, the membrane potentials *v*_*b*_ and *v*_*a*_ of the basal and the apical compartment, the exchange current 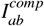 (Eq. 8), and the membrane potentials of the PV, SOM, and VIP cells. The exchange current is positive from the apical to the basal compartment. The bottom row shows the firing rate of the TPN (Eq. 23) together with the rates of the multi-compartment model for the two input conditions (red dashed). Thick lines show the deterministic simulation from the resting state of Tab. A.1 with *τ*_*b*_ = 10 ms, *τ*_*a*_ = 35 ms, and *τ*_*q*_ = 1 ms. Thin lines show one simulation with additive noise on *v*_*b*_ and *v*_*a*_ (*σ*_*b*_ = *σ*_*a*_ = 1), whose rate is smoothed over 10 ms. The dotted horizontal line marks the threshold *ρ*_*a*_ of the apical gate (Eq. 6), and the detail on the right shows *v*_*b*_ and *v*_*a*_ between 140 and 240 ms (gray frame). Shaded intervals and a solid line mark the four phases described in the text, and circled numerals mark the events that define them. Phase I starts at FB onset, when SOM cells are silenced, and ends when *v*_*a*_ reaches *ρ*_*a*_ at 167 ms and the apical gate opens (phase II). Phase III marks the reversal of the exchange current, when *v*_*a*_ exceeds *v*_*b*_ at 174 ms (solid line), and phase IV lasts until *v*_*b*_ has completed 95 % of its rise after 174 ms, i.e., until 233 ms. Figure 5(C) TPN state at equilibrium for the different combinations of basal and apical input strengths of panel A under constant FF and FB input. The left column shows *v*_*b*_, *v*_*a*_, and 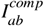 on a diverging color scale that is light gray at zero, and the right column shows the membrane potentials of the PV, SOM, and VIP cells, all in model units (a.u.). Dots mark the two input conditions of panel B. Multi-compartment pyramidal cell schematic in A reproduced from Shai et al. (2015) under the CC BY 4.0 license (https://creativecommons.org/licenses/by/4.0/).

#### 2.3.1. Comparison with a multi-compartment neuron model

The reference data are the somatic output rates of the multi-compartment model of a layer 5b pyramidal cell of Shai et al. (2015), in which *n*_*b*_ *∈* {0, 10, … , 300} basal and *n*_*t*_ *∈* {0, 10, … , 200} tuft synapses were activated at random times within 100 ms (651 input conditions, ModelDB accession 180373, control condition). The output rate is the inverse of the shortest inter-spike interval of the simulated response (Appendix B.1). We denote the rate for *n*_*b*_ basal and *n*_*t*_ tuft synapses by *f* (*n*_*b*_, *n*_*t*_) and the prediction of a model by 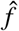, with a subscript that names the model. The logistic function is denoted by *σ*(*x*) = 1*/*(1 + *e*^−*x*^).

Five models were fitted to these data. In the *composite model* of Shai et al., the tuft input sets the maximum *M* and the threshold *T* of a sigmoidal function of the basal input,

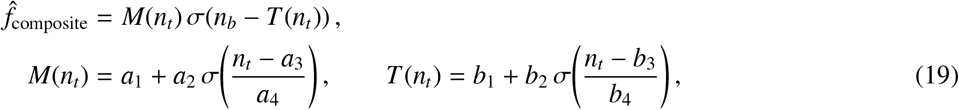

with *k* = 8 parameters. The *additive* and *multiplicative models* described by Shai et al. add or multiply the outputs of two separate sigmoids of the tuft and the basal input,

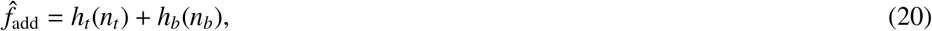

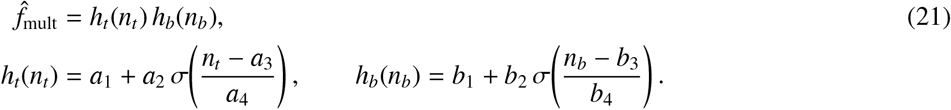

Both models have *k* = 7 identifiable parameters, because *a*_1_ and *b*_1_ enter the additive model only through their sum, and a common factor can be moved between *h*_*t*_ and *h*_*b*_ in the multiplicative model. The *Mod-E-I model* is the steady state of the TPN reduced to a point neuron (Eqs. E.1–E.4 and Fig. 2C) with feedforward inhibition,

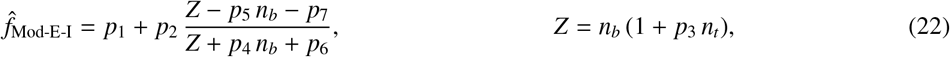

where the term *Z* implements the modulation FF·(1+FB) of the basal drive introduced in Sect. 2.1.1, with *p*_2_, … , *p*_7_ ≥ 0 and *k* = 7 parameters (derivation in Appendix B.1). The *TPN* keeps its internal parameters, which were tuned during model development, in part by a non-exhaustive search against the same data (Appendix B.1), and which are not counted in *k*. It was fitted only through a linear map of the synapse numbers onto its inputs, *r*_*FF*_ = *s*_*b*_ *n*_*b*_ and *r*_*FB*_ = *s*_*t*_ *n*_*t*_, and an affine map of its output (Eq. 10) onto the firing rate,

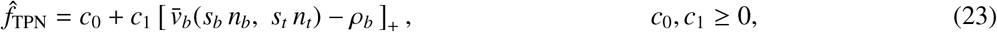

with *k* = 4 parameters. Here, 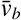 is the basal potential at equilibrium under constant input. It was obtained by integrating the TPN equations (Eqs. 1–10) from the resting state of Tab. A.1, i.e., the equilibrium without input, with the forward Euler method (time step 0.05 ms) until the state settled, on a grid of 77 × 77 inputs *r*_*FF*_, *r*_*FB*_ *∈* [0, 4]. Under the fitted map, the largest synapse numbers of the data, 300 basal and 200 tuft synapses, correspond to the inputs *r*_*FF*_ = 1.96 and *r*_*FB*_ = 0.95. The input-output maps of Fig. 5A,C show the grid up to these inputs. The time courses of Fig. 5B were integrated from the same resting state with a time step of 0.1 ms.

All models were fitted to the same data by minimizing the sum of squared errors (SSE) over the 651 conditions. The models of Shai et al. and the Mod-E-I model were fitted by nonlinear least squares from 500 starting points, which include the published parameters of the composite model. The TPN map was fitted by a search over a regular grid of input scales *s*_*b*_ and *s*_*t*_ followed by a local refinement, in which offset and gain follow from linear least squares for every pair of scales (Appendix B.1). The goodness of fit is the coefficient of determination *R*^2^ = 1 − SSE*/*SST, where SST is the sum of squared deviations of the data from their mean. *R*^2^ is the fraction of the variance of the data that a model explains and is reported in percent in the main text.

#### 2.3.2. Evaluation of the learned TPN pool

The learned pool of two TPNs (Fig. 3) is evaluated in test trials with frozen synaptic efficacies and without dopamine. Its decision function assigns class A (horizontal bar, *y*^*A*^) to TPN 1 and class B (vertical bar, *y*^*B*^) to TPN 0. A trial is decided for class A if the margin *m* = *r*_1_ − *r*_0_ between the output rates of the two TPNs (Eq. 10) at the end of the presentation is positive, and for class B if it is negative. During training, the teacher applies the same comparison to set the sign of the dopamine release (Sect. 2.2.2).

Each test condition, i.e., each combination of a stimulus and a context, is tested in repeated trials that start from different random initial states, and *P*(*A*) denotes the fraction of trials that are decided for class A. The *accuracy* is the fraction of correct decisions on the four stimulus-context pairings of the training, with 10 trials per pairing, where undecided trials with |*m*| ≤ 0.05 count as errors (Fig. 6A,E). *Ambiguous stimuli* are pixel-wise mixtures *y*(*α*) = (1 − *α*) *y*^*A*^ + *α y*^*B*^ of the two classes with *α ∈* [0, 1]. Each network has its own point of subjective equivalence 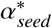 , at which the margin vanishes for silenced context, and its own ambiguous stimulus 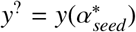 . *Neurometric curves* (Newsome et al., 1989; Britten et al., 1992) give *P*(*A*) for 21 mixtures, with *α* from 0 to 1 in steps of 0.05 and 40 trials per condition. Each curve is measured under one of four contexts, i.e., horizontal flanker bars *c*^*A*^, vertical flanker bars *c*^*B*^, their uninformative superposition *c*^?^, or silenced context. The neurometric curves are aligned at 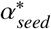 before they are averaged across networks (Fig. 6C). The *context effect* Δ = *P*(*A* | *y*^?^, *c*^*A*^) − *P*(*A* | *y*^?^, *c*^*B*^), again with 40 trials per condition, measures how strongly informative context shifts the decision on the ambiguous stimulus (Fig. 6A,E). To test the role of the tag and of the neuromodulatory factor, either or both were replaced by the constant 1 in Eq. 12. This yields ablated three- and two-factor variants of the rule, with everything else unchanged (Fig. 6A). All measures are averaged over 30 independently initialized networks, and additional details are given in Appendix C.2.

**Figure 6:**
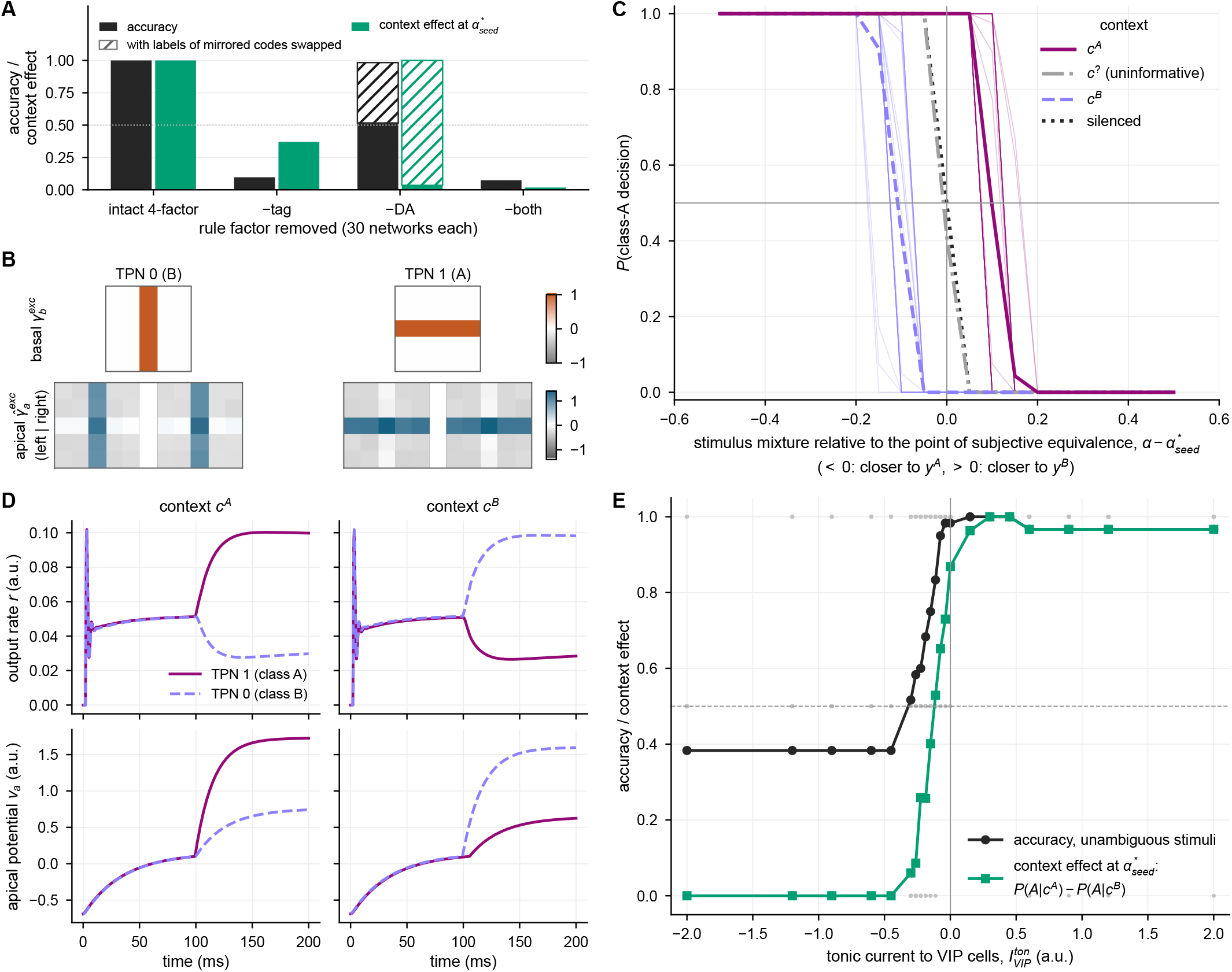
Learning in the two-point neuron (TPN) microcircuit via apical-basal coincidence and VIP dis-inhibition. Panels A–D show networks that were trained with a tonic input 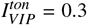 to VIP neurons. (A) Accuracy on the unambiguous stimuli (black) and context effect Δ at each network’s point of subjective equivalence (green) after training, each for 30 differently seeded training runs. The four-factor rule (Eq. 12) is shown intact and with its coincidence tag (−tag), its dopamine signal (−DA) or both removed. Without dopamine, about half of the networks learn a mirrored code, in which each TPN prefers the stimulus and the context of the other class. The hatched bars show the accuracy and the context effect with the class labels of these networks swapped (Appendix C.3). (B) Basal efficacies 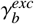 from the 5 × 5 stimulus pixels (top) and apical efficacies 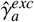 from the units of the left and the right flanker patch (bottom) after learning the orientation discrimination task. The efficacies of TPN 0 (left) and TPN 1 (right) are shown for one example training run out of 30, with positive values in orange (basal) or blue (apical) and negative values in gray. The decision function assigns the vertical bar (class B) to TPN 0 and the horizontal bar (class A) to TPN 1. (C) Neurometric response curves for 30 differently seeded training runs, i.e., the fraction *P*(*A*) of decisions for class A over the stimulus mixture, for decisive context *c*^*A*^ (magenta) or *c*^*B*^ (lavender, dashed), uninformative context *c*^?^ (gray, dash-dotted), and silenced apical input (black, dotted). Response curves are obtained as mean response per input across 40 test trials with different initial system states and are plotted relative to each network’s point of subjective equivalence 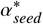, such that negative values of 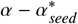 denote mixtures closer to *y*^*A*^ and positive values mixtures closer to *y*^*B*^. Average response curves are displayed in saturated hue and individual response curves for *c*^*A*^ and *c*^*B*^ in light hue. (D) Example trial after learning, started from the resting state, at the point of subjective equivalence with decisive context *c*^*A*^ (left) or *c*^*B*^ (right) for the same example training run. The top row shows the output rates *r* that the decision function reads, and the bottom row shows the apical potentials *v*_*a*_, each for TPN 1 (magenta, solid) and TPN 0 (lavender, dashed). (E) Effect of tonic input 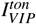 to VIP neurons, applied during training and testing, on the learned accuracy for unambiguous stimuli (black circles) and on the context effect Δ at each network’s point of subjective equivalence (green squares, Sect. 2.3.2). The solid lines mark the averages across 30 differently seeded training runs per value of 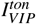, and gray dots the accuracies of individual runs. The average context effect includes only the runs that have a point of subjective equivalence (Appendix C.3). The dashed line marks an accuracy of 0.5, and the vertical line neutral tonic input.

## 3. Methods – Network-level motifs of two-point neuron coupling

So far, the proposed two-point neuron mechanism has defined a zero-dimensional (0-D) model with a single layer V pyramidal (L5Pyr) cell and its adjacent inhibitory interneurons. The previous section explained how the model’s cell-level motifs capture biological mechanisms important for both processing and learning. At the *network level*, the model proposes how interaction between TPNs gives rise to competition and cooperation required for processing and learning to unfold. Related networks have been studied before with the simplified units of Fig. 2B,C (Bayerl and Neumann, 2004; Bouecke et al., 2011; Schmid and Neumann, 2025). Here, every network node is a TPN with its own PV, SOM, and VIP cells, and context reaches it only through its apical compartment and its VIP cell.

### 3.1. Network-level motifs

Here, we describe how interactions between pools of TPNs are structured at the network level. The proposed network architecture is broadly applicable to scenarios where neural population codes are organized along space and feature dimensions.

#### 3.1.1. Connectivity schemes

The network is based on canonical structuring principles and distinguishes between local populations and global network topology. *Local populations* of neurons consist of interconnected TPNs that share tuning properties toward the same region in the input domain. Such interconnected TPNs form a one-dimensional (1-D) neural field, in which interactions depend on the distance between neurons along the feature dimension (Amari, 1977). Here, the TPNs of a population form a ring of laterally coupled neurons with circular boundaries (Fig. 4, top). Excitation between similarly tuned neurons and broad inhibition define the dynamics of such ring attractors, whose activity forms a localized bump on a ring-shaped, low-dimensional manifold (Ben-Yishai et al., 1995; Khona and Fiete, 2022). The bump represents the population’s estimate of the feature value at its location^3^. Neurons within a ring are interconnected by three connectivity patterns:

- Pyr-Pyr: TPNs project onto the basal compartments of TPNs with similar preferred feature values, with weights that decrease with the difference between the preferred values, as found, e.g., for layer 2/3 pyramidal cells of mouse V1 (Ko et al., 2011; Cossell et al., 2015). This strengthens the evidence for similar feature values.
- Pyr-PV: TPNs uniformly project to the PV cells of all TPNs within the ring, including their own. This establishes pooled inhibition, as each PV cell receives the summed activity of all TPNs within the ring, and, in turn, inhibits the basal activity of its TPN (Kerlin et al., 2010; Carandini and Heeger, 2012).
- SOM-VIP: SOM cells project inhibitorily onto the VIP cells of TPNs with similar preferred feature values, including their own (Pfeffer et al., 2013). This puts the dis-inhibition of apical integration in competition with that of neighbors along the feature dimension. The feature-specific spread of this projection is a model assumption.

The *global network topology* arises in the model from the interconnection of the local populations. For simplicity, we here assume a generic line topology that organizes populations as neighbors along a 1-D spatial domain (Fig. 4, bottom). Two projection types determine the network’s information flow:

- forward projections: external input is forward projected into the network and excites the basal TPN compartments and PV cells of each population. From this driving input each TPN extracts features according to its tuning properties.
- contextual projections: each TPN projects to the apical compartment as well as to the VIP cell of the TPN with the same preferred feature value in each of the two neighboring populations. These inter-population projections serve as modulating context and yield cooperation among populations. Additionally, further context can enter the network via the same projection targets as external FB.

All network connections are fixed, and the synaptic learning of Sect. 2.2 is not applied in the network. Overall, the model proposes how TPNs are organized into local populations and how connectivity among populations forms a structured network. This connectivity provides the means to turn the TPN’s cell-level processing motifs into network-level computational capabilities of evidence integration, competition, and cooperation.

#### 3.1.2. Cell-level computational motifs at the network level

The network model consists of the same model TPNs with their generic processing motifs (Sect. 2.1.1). Their equations and cell parameters are unchanged, and the network only adds connections between TPNs, including a dendritic threshold on the contextual projections onto the apical compartments. Their PV and VIP cells now pool input over several TPNs. The gains of these pooled projections were adjusted accordingly (Tab. A.1). When paired with the connection structure described above, the TPN’s generic processing motifs give rise to computational function at the network level.

##### Feature extraction

We here assume that TPNs are tuned toward a feature of a certain location within a topological domain, e.g., retinotopic space. TPNs within a population encode the same location but differ in their preferred feature value, while TPNs at different locations may share the same preferred value. The feature value at the population’s location can then be decoded from its population vector (Georgopoulos et al., 1986) (Sect. 3.3.2).

##### Pool competition

Each population establishes pool competition by its Pyr-PV projections (Sect. 3.1.1). Because PV cells inhibit the basal compartment through a conductance with a reversal potential below rest (Sect. 2), pooled inhibition implements a divisive-like normalization of the TPNs’ activities by the summed activity of the pool (Grossberg, 1973; Carandini and Heeger, 2012; Brosch and Neumann, 2014a). As a result, the stronger a neuron fires, the stronger it contributes to the pool and the stronger it suppresses its competitors.

##### Feedback modulation

Combining pool competition with asymmetric FF-FB integration yields an implementation of *biased competition* (Desimone and Duncan, 1995). Biased competition proposes how neuron populations can integrate contextual FB evidence with the existing FF evidence about the presence of a feature. Neurons compete for representing the input through mutual suppression, and FB signals bias this competition in favor of those neurons whose FF evidence is consistent with the context. If only FF signals are available, the firing rate output of the TPN population resembles the pattern of the FF signals filtered by the input weights to the basal compartment. If additional FB signals provide matching contextual input, they amplify TPNs with coinciding FF signals and effectively tip the balance of the competition toward these neurons in the population.

##### Cooperation

Contextual projections couple TPNs with the same preferred feature value at neighboring locations, and through the VIP cells of these TPNs they release the apical compartments from SOM inhibition. Only afferent rates above a dendritic threshold contribute to apical integration (Sect. 3.2). Weakly driven populations therefore do not bias each other, whereas a strongly active population biases the competition of its neighbors where its context coincides with their FF input. In this way, evidence can propagate from population to population across the network. In sum, the TPN’s generic processing motifs establish competition and cooperation at the network level. TPN populations form local pools of biased competition to integrate evidence about input features from the forward projections with the context of neighboring populations.

### 3.2. Neural field equations

The neural field of TPNs extends the model of a single TPN unit (Sect. 2) to a network of *R* ring populations at the locations *x*_*k*_, *k* = 0, … , *R* − 1, each with *N* TPNs with the preferred feature values *θ*_*n*_ = 2*πn/N, n* = 0, … , *N* − 1. The former 0-D model with scalar equations (Eqs. 1–10) now becomes two-dimensional (2-D) over locations *x*_*k*_ and feature values *θ*_*n*_, i.e., *v*_*i*_ turns into *v*_*i*_(*x*_*k*_, *θ*_*n*_) with *i ∈* {*b, a*, PV, SOM, VIP}, and likewise for the rates *r*_*i*_. The connection patterns and their parametrization are the only differences between the 0-D TPN model and the 2-D model. The former scalar rates *r*_*j*_ and conductances 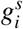 now become 2-D fields **r**_*j*_ and 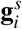 over all locations and feature values, and the former scalar synaptic efficacies 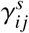 (Eq. 5) become weight matrices 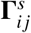,

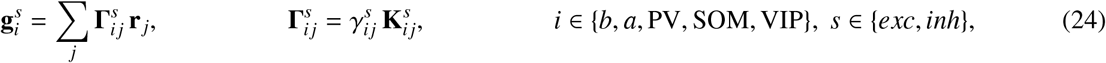

where the kernel 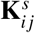 specifies how the projection from element *j* to element *i* spreads over locations and feature values.

Synaptic weights 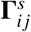 are the structuring element of the 2-D network. They organize TPNs locally into ring populations along the feature dimension *θ* and ring populations globally into a network along the locations *x*. The network is *x*-*θ* separable. For a source at (*x*_*l*_, *θ*_*m*_) and a target at (*x*_*k*_, *θ*_*n*_), every kernel weight is the product of a spatial and a feature weight, 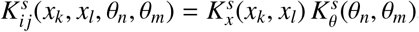, where the indices *i j* of the factors are omitted. With the TPNs ordered by location and, within each location, by feature value, the kernel is thus the Kronecker product 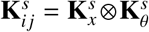 of two sub-matrices that depend on only one dimension. For the given network, each projection from *j* to *i* is formed along one dimension only, similar to the separate stages of spatial and orientational competition of Grossberg and Mingolla (1985). Such one-dimensional projections further reduce the complexity of synaptic connectivity (either 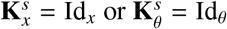, with identity matrix Id). Furthermore, apart from the ends of the line, the kernels only depend on the circular distance *d*(*θ*_*n*_, *θ*_*m*_) between preferred feature values and on the distance |*k* − *l*| between locations. Hence, they can be described as vectors used as kernels in a convolutional filtering operation. Four types of kernels exist in the network:

- local 1:1 (0-D) projections 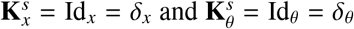 with Kronecker delta *δ*, i.e., the Dirac impulse of the discrete field,
- uniform projections 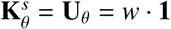 with the all-ones matrix **1** and the same weight *w* for all TPNs of a ring,
- Gaussian projections 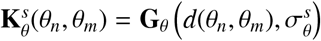 with standard deviation 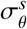, centered at the target’s feature value *θ*_*n*_,
- a surround kernel **S**_*x*_ with equal weights for the two immediate neighbors (|*k*−*l*| = 1) and zero weights otherwise. Since the end rings of the line have only one neighbor, they receive twice this weight from it, so that all rings receive the same total weight of contextual input.

The resulting connectivity structure can be found in Tab. 2, and the widths and weights of the kernels in Tab. A.1.

**Table 2:** 2-D network connectivity. Rows represent source elements (*from, j*), columns represent target elements (*to, i*), and entries list the kernel 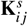 of the projection (Eq. 24) with the channel types *s* of Tab. 1. All connections span either the space dimension *x* or the feature dimension *θ* only. Apical TPN compartments *a* are electrically coupled to their own basal compartments *b* and receive firing rate input from neighboring TPNs. Projections with **S**_*x*_ onto the apical compartments pass the dendritic threshold of Eq. 25. Kernel widths and weights are listed in Tab. A.1. The kernels are the identity 1:1 mapping Id, the Gaussian kernel **G**_*θ*_, the uniform kernel **U**_*θ*_, and the surround neighborhood kernel **S**_*x*_, and **Comp** denotes compartmental exchange.

|  |  | To |  |  |  |  |
| --- | --- | --- | --- | --- | --- | --- |
|  |  | b | a | PV | SOM | VIP |
| From | **FF** | Id | – | Id | – | – |
|  | **FB** | – | Id | – | – | Id |
| | **b** | $\mathbf{G}_\theta$ | Comp, $\mathbf{S}_x$ | $\mathbf{U}_\theta$ | Id | $\mathbf{S}_x$ |
|  | **a** | Comp | – | – | – | – |
|  | **PV** | Id | – | – | – | – |
| | **SOM** | – | Id | – | – | $\mathbf{G}_\theta$ |
|  | **VIP** | – | – | – | Id | – |

The contextual projections (**S**_*x*_) target the arborized dendrites of the apical tuft and the VIP cells. For the projections onto the apical dendrites, the afferent rates **r**_*j*_ in Eq. 24 are replaced by

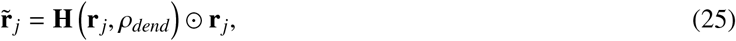

where **H**(**r**_*j*_, *ρ*_*dend*_) applies the Heaviside function (cf. Eq. 7) element-wise with a dendritic threshold *ρ*_*dend*_ on the afferent rates, and *⊙* denotes element-wise multiplication. With this mechanism, only afferents with a rate above the threshold contribute to the weighted sum, so that weak afferents cannot add up to the input of a strongly active one. The VIP cells receive the contextual projections without this threshold, and external FB input enters both targets without it. The voltage-dependent gating of the summed apical input (Eq. 6) remains in place.

### 3.3. Network simulations, readouts and evaluation

#### 3.3.1. Simulations

The simulated networks consist of *R* = 5 ring populations for the line of Fig. 7 and of *R ∈* {10, 20, 40, 100} ring populations for the bars of Fig. 8, each with *N* = 200 TPNs. FF and FB inputs are Gaussian profiles over *θ* with additive Gaussian noise, which is drawn independently for every TPN (Eq. D.1, Tab. D.1). The noise is held constant within time bins of 1 ms, or of 5 ms for bars of 40 and 100 rings, so that the noisy inputs can be tabulated in advance for the adaptive Runge–Kutta solver (Tab. A.2). Apart from this input noise, the simulations are deterministic (N(*σ*_*i*_) = 0 in Eqs. 1–3). The simulations start from the resting state of the network with small random offsets of the basal potentials, the input noise is drawn from seeded generators, and each configuration of the line and each bar length is simulated once (Appendix D.1).

**Figure 7:**
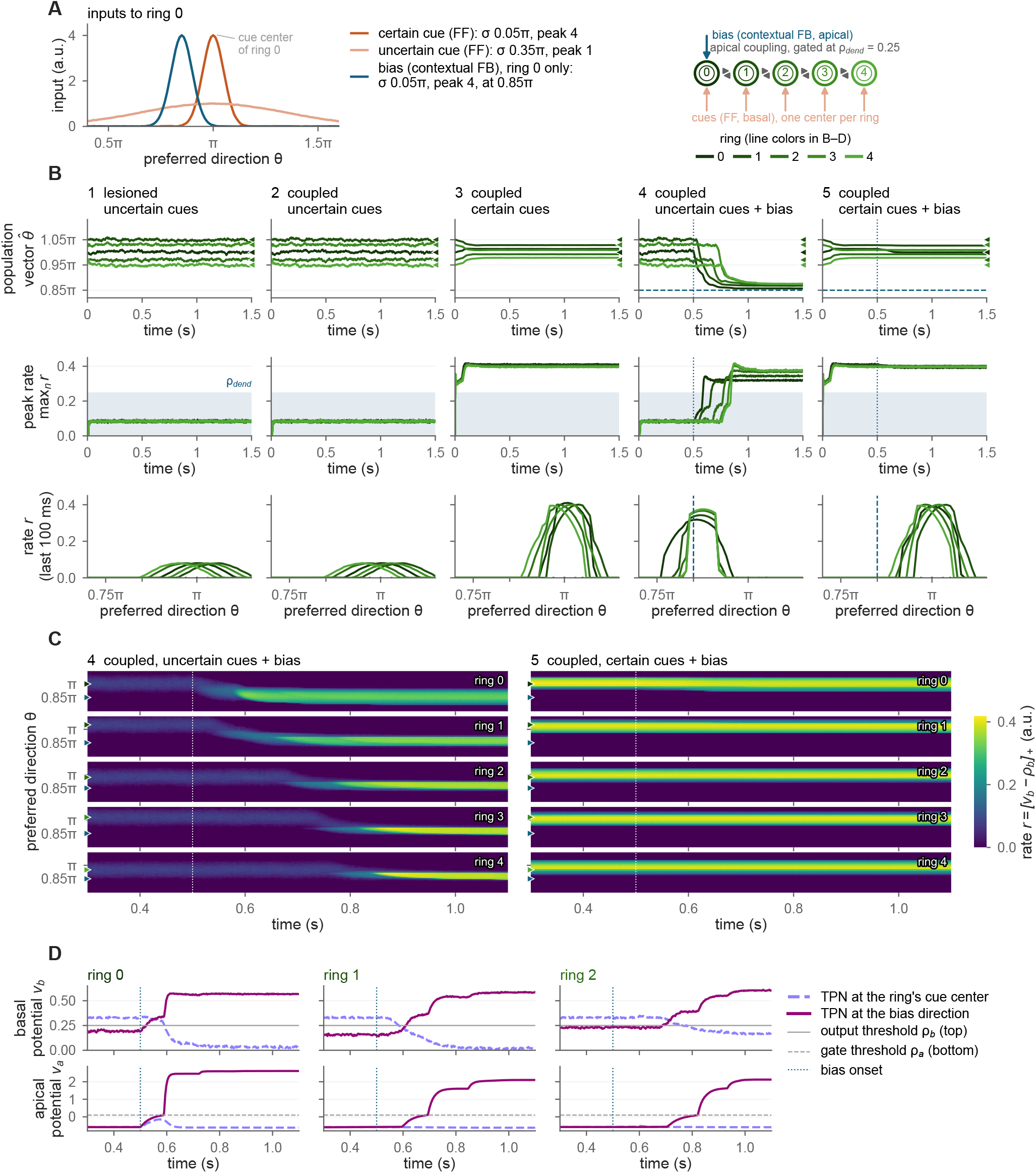
Competition and cooperation for evidence integration among a network of TPNs. (A) Configuration of a TPN network model with five ring attractor populations of *N* = 200 TPNs organized into a line (right; cf. Fig. 4) and its basal and apical input distributions among a set of simulations (left). The inputs are shown for ring 0 and averaged over their noise samples. Feedforward input excites each population and is either uncertain and broadly tuned (*σ* = 0.35*π*, peak amplitude 1) or certain and narrowly tuned (*σ* = 0.05*π*, peak amplitude 4). Its center lies at 1.00*π*, 1.05*π*, 0.97*π*, 1.03*π* and 0.95*π* in rings 0 to 4. Biasing contextual feedback is either absent or excites ring 0 from *t* = 0.5 s on with a narrowly tuned input (*σ* = 0.05*π*, peak amplitude 4) centered at 0.85*π*. Neighboring rings are coupled by contextual projections onto the apical compartments, which pass only afferent rates above the dendritic threshold *ρ*_*dend*_ = 0.25 (Eq. 25), and onto the VIP cells. (B) Network simulations with different input combinations. Line colors from dark to light green denote rings 0 to 4 (as in A). From left to right, the columns show the lesioned network with uncertain cues (1) and the coupled network with uncertain cues (2), certain cues (3), uncertain cues and bias (4), and certain cues and bias (5). The top row shows the population vector 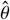 of each ring (Eq. 26) with the cue centers as triangles on the right and the bias direction as a dashed blue line. The middle row shows the peak rate max_*n*_ *r* of each ring with the range below *ρ*_*dend*_ shaded in light blue. Both rows show the first 1.5 s of the 3-s simulations, after which population vectors and peak rates stay nearly constant. The bottom row shows the rate profile *r* (Eq. 10) of each ring over the preferred direction, averaged over the last 100 ms of the simulation. (C) Rate *r* of the TPNs with preferred directions between 0.7*π* and 1.15*π* in each ring (rows) over time between 0.3 and 1.1 s for the two diagnostic cases of biasing feedback (4 and 5). Green and blue triangles on the left mark the ring’s cue center and the bias direction. (D) Example time courses of the basal potential *v*_*b*_ (top) and the apical potential *v*_*a*_ (bottom) in rings 0 to 2 for the case of uncertain input and biasing feedback (4). Each panel shows the TPN closest to the ring’s cue center (lavender, dashed) and the TPN closest to the bias direction (magenta, solid). Gray lines denote the output threshold *ρ*_*b*_ = 0.25 (top) and the apical gate threshold *ρ*_*a*_ = 0.1 (bottom, dashed; Eq. 6). Vertical dotted lines in B to D denote the onset of biasing contextual feedback. Rates and potentials are given in model units (a.u.).

**Figure 8:**
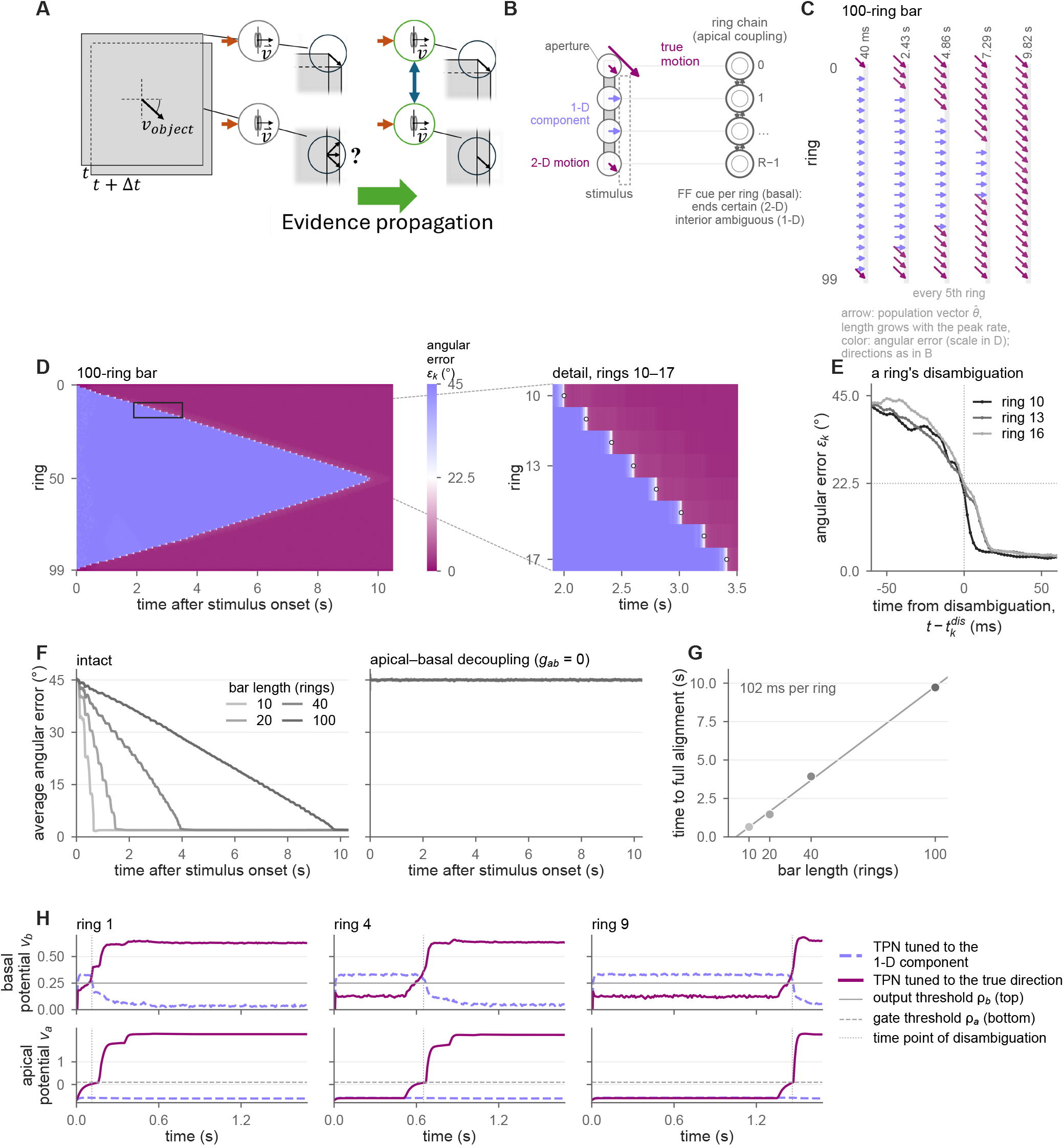
Visual motion integration in a TPN network. (A) Visual motion integration and the aperture problem. The aperture problem arises from the restriction of receptive fields to locally available motion cues (black circles on the object outline). Local motion cues are ambiguous for receptive fields covering edges of objects, while unambiguous cues can only be found at corners. (B) TPN network setup. The simulated TPN network consists of a line of interconnected ring populations (cf. Fig. 4). It instantiates the aperture problem by receiving motion information similar to that of neurons with receptive fields positioned along a moving bar. Each of the *R* locations along the bar is represented by a ring population of *N* = 200 TPNs (rings 0 to *R* − 1), and neighboring rings are coupled by contextual projections onto their apical compartments as in Fig. 7. Unambiguous cues are provided at end positions and ambiguous cues at interior positions along the outline. The two end rings receive a certain FF cue centered at the true motion direction 0.75*π* (*σ* = 0.05*π*, peak amplitude 4), whereas the interior rings receive an uncertain FF cue centered at the 1-D motion component *π* (*σ* = 0.35*π*, peak amplitude 1). (C) Formation of a coherent representation. The motion direction encoded by local ring populations dynamically changes over time. Arrows show the population vector 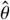 (Eq. 26) of every fifth ring of a bar of 100 rings at five points in time after stimulus onset. As in B, they point right for the 1-D motion component (*π*) and down-right for the true motion direction (0.75*π*), while their length grows linearly with the ring’s peak rate and their color codes the angular error (color scale in D). At first only the end points of the bar outline encode the true motion direction. (D) Localized angular error over time. The angular error *ε*_*k*_ (Eq. 27) of every ring of the 100-ring bar is color-coded over time (left). A detail of rings 10 to 17 at the output resolution of 2 ms (rectangle in the left map) marks their time points of disambiguation by circles (right; Eq. 29). Positions receiving ambiguous motion cues represent 1-D motion components at first with an absolute angular error of 45^°^. (E) Prototypical disambiguation behavior. The angular errors of rings 10, 13 and 16 of the 100-ring bar are aligned to their time points of disambiguation, and dotted lines mark this time point and the 22.5^°^ criterion. (F) Average angular error for different bar lengths and for a decoupled variant. The average angular error over the interior rings (Eq. 28) is plotted over time for bars of 10, 20, 40 and 100 rings in the intact network (left) and in a decoupled variant (right). In the decoupled variant, the exchange conductance between the basal and apical compartments of every TPN is set to zero (*g*_*ab*_ = 0, Eq. 8). (G) Time to full alignment. The time point at which the last interior ring of each bar disambiguates its estimate is plotted over the bar length together with a least-squares line, whose slope is 102 ms per ring of bar length. (H) Integration dynamics among individual rings. Basal potential *v*_*b*_ (top) and apical potential *v*_*a*_ (bottom) of the TPN tuned to the 1-D motion component (lavender, dashed) and of the TPN tuned to the true motion direction (magenta, solid) are shown for rings 1, 4 and 9 of a bar of 20 rings. Gray lines denote the output threshold *ρ*_*b*_ = 0.25 (top) and the apical gate threshold *ρ*_*a*_ = 0.1 (bottom, dashed), and dotted vertical lines denote each ring’s time point of disambiguation. Potentials are given in model units (a.u.).

#### 3.3.2. Population readouts

Three quantities describe the state of a ring population, all computed from the output rates *r*(*x*_*k*_, *θ*_*n*_, *t*) = *r*_*b*_(*x*_*k*_, *θ*_*n*_, *t*) of its TPNs (Eq. 10). The feature value that the ring at location *x*_*k*_ represents is read out by its *population vector* (Georgopoulos et al., 1986), the direction of the rate-weighted circular mean (imaginary unit i),

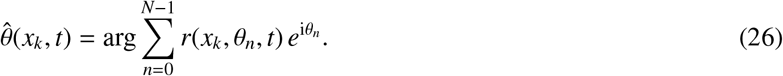

It is undefined while all TPNs of the ring are silent, which happens only during the first milliseconds after stimulus onset. The activity of the ring is additionally summarized by its *peak rate* max_*n*_ *r*(*x*_*k*_, *θ*_*n*_, *t*), and its *rate profile r*(*x*_*k*_, *θ*_*n*_, *t*) over the preferred feature values shows the whole distribution of activity. Population vector and peak rate are evaluated at every stored time step (2 ms) without temporal averaging.

#### 3.3.3. Line and bar simulations

In the simulations of the line (Fig. 7), the FF inputs are presented alone for 0.5 s. They are followed by 2.5 s with or without the biasing contextual FB to the ring at *x*_0_. The rate profiles of Fig. 7B are averaged over the last 100 ms of the simulation. Peak rates and population vectors of the end state are averaged over the last 0.5 s. The shift of the ring at *x*_0_ toward the bias is measured relative to the circular mean of its population vector over 0.4 to 0.5 s. A ring takes up the bias when its peak rate first exceeds *ρ*_*dend*_ after the bias onset. The steps from ring to ring are the differences of these times.

In the simulations of the bars (Fig. 8), all inputs are present from stimulus onset for 12 s. The two end rings receive a certain FF input at the true motion direction *θ*_*true*_ = 0.75*π*, and the interior rings receive an uncertain FF input at the 1-D component *π*. The *angular error* of a ring is the absolute difference between its population vector and the true direction,

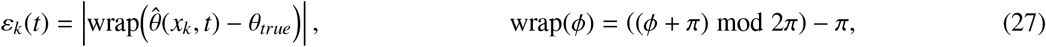

given in degrees between 0^°^ and 180^°^, so that the 1-D component has an error of 45^°^. The *average angular error* of a bar (Fig. 8F) is the mean error of its *R* − 2 interior rings,

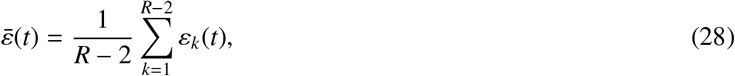

because the end rings receive the true direction directly. It is computed once all rings of the bar respond, since a silent ring has no population vector. A ring has *disambiguated* its estimate once its error stays below 22.5^°^, the midpoint between the two candidate directions, for the rest of the simulation, i.e., at the time point of disambiguation

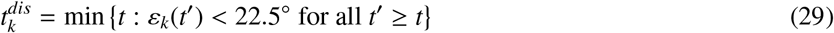

(Fig. 8D,E,H). The time to full alignment of a bar is the time point at which its last interior ring disambiguates its estimate, and a least-squares line relates it to the bar length (Fig. 8G). The slope of the time points of disambiguation over the distance of the interior rings from the nearest end gives the time in which each of the two fronts advances by one ring.

## 4. Results – Simulation of two-point neuron local circuitry

The following subsections investigate the TPN model’s behavior in simulation with respect to its local processing and learning. The simulations demonstrate how its computational motifs account for biologically detailed data, as well as display its learning capabilities and how these are rooted in the same motifs and mechanisms.

### 4.1. Explanatory power for multi-compartment data

We simulate the dynamics of the TPN model and its local inhibitory interneurons to relate the model to biological evidence and to make the interplay of its processing motifs visible. The model has been run with different combinations of basal and apical input and monitored for its response behavior (Fig. 5). A similar investigation by Shai et al. (2015) employed a biophysically detailed spike-based multi-compartment model. This model was adjusted to patch-clamp recordings from L5Pyr cells in slices of mouse primary visual cortex (see Shai et al., 2015 for details). Hence, it can serve as a reference to fit the TPN, the simplified Mod-E-I model from earlier investigations (Brosch and Neumann, 2014a), as well as the abstract functional models from Shai et al. (2015) against the multi-compartment model data provided by them^4^ (Fig. 5A; see Sect. 2.3.1 for further details). The data of the multi-compartment model are visualized in a two-dimensional map of basal and apical tuft input dimensions. In this map two input-dependent regions can be distinguished. One region with intermediate activations appears for moderate to strong basal input combined with weak apical tuft input. Another region shows elevated activity for moderate to strong apical input over most of the range of basal input. To fit the data, Shai et al. (2015) specified descriptive models in which the basal and apical output functions are combined differently (Fig. 5A). Among the proposed mechanisms, the composite model best explains the data (90.1 % explained variance after refitting; 87.8 % with the parameters published by Shai et al. 2015). The TPN model proposed here ranks second in explaining the data (88.6 %), followed by the additive model (80.1 %). The TPN model and the composite model also qualitatively capture the existence of two input-dependent regions. The multiplicative model and the Mod-E-I model fit the data with the lowest explanatory power (78.8 % and 77.9 %, respectively). Qualitatively, the additive, the multiplicative, and the Mod-E-I model exhibit the main feature of coincidence detection (higher activity in the presence of strong basal and apical input) but do not account for the existence of the two separate regions of activity in the basal-tuft input space. The additive and the multiplicative model rise smoothly with tuft input and are already active for moderate basal input without tuft input. The Mod-E-I model, in turn, lowers its basal threshold with tuft input but responds to basal input alone with at most 26 Hz instead of about 70 to 100 Hz. Except for the additive model, which also responds to tuft input alone, common to all models is the main feature of asymmetric coincidence detection, i.e., the absence of activity for missing basal input.

Unlike the class of abstract functional models from Shai et al. (2015), the TPN model is a dynamical model. Thus, it allows for a time-resolved investigation of its computation and provides a mechanistic view on how the two distinct interacting compartments generate input-dependent activity patterns. Two main aspects of the TPN model’s processing appear to be important here, namely the apical-basal integration mechanism and the interplay between the TPN and the inhibitory interneurons. We investigate the *apical-basal integration mechanism* computationally in simulations with time-dependent pairing of feedforward (FF) and feedback (FB) input (Fig. 5B). A time-resolved visualization of the basal and apical membrane potentials shows the phasic convergence to specific operating regimes. After FF input becomes available, the basal membrane potential starts to rise and settles to an equilibrium. While FB input is missing, the apical compartment remains hyperpolarized and the basal compartment remains in a regime of intermediate depolarization. When coinciding FB input is present, a cascade of mechanisms is set into motion in a sequence of four phases (compare to the BAC schematic in Fig. 1B):

I. I Apical inhibition is lifted and basal-to-apical influx occurs depolarizing the apical compartment. Overall, the cell becomes more depolarized in the absence of apical inhibition.
II. Once the apical depolarization reaches the threshold *ρ*_*a*_, the voltage-dependent apical gate opens (Eq. 6), allowing for a strongly depolarizing influx of direct FB input to the compartment.
III. After a first steep rise the apical compartment becomes more depolarized than the basal compartment, and the exchange current changes direction.

#### IV A strong apical-to-basal influx now drives the basal compartment to settle into a regime of high depolarization

In sum, this phasic interplay of basal and apical depolarization and exchange captures the main aspects of BAC firing at the level of firing rates, namely the causal order of basal-apical coincidence detection and the amplifying impact on the output firing rate. It explains the strong enhancement of the TPN’s activity in the presence of coinciding contextual FB input. The phases can be mapped onto the steps of BAC firing (Fig. 1B), as phase I corresponds to the depolarization of the apical dendrite by the back-propagating action potential, phase II to the crossing of the calcium-spike threshold with coinciding tuft input, and phases III and IV to the calcium plateau that drives the soma into high-rate firing. Yet, the millisecond coincidence window, the calcium-spike kinetics, and the bursts are not resolved, and the time course reflects the population and membrane time constants of the rate model^5^.

The *interaction between the TPN and the inhibitory interneurons* becomes understandable by comparing the respective activities for the different FF and FB input strengths (Fig. 5C). Three distinct regions of qualitatively different joint activity become visible, namely a low apical input regime, a low basal input regime, and a coinciding input regime:

- For low apical input, VIP cells remain silent, while PV and SOM cells become increasingly active with increasing basal input. This increase is matched by an increase in basal, but not apical, activity, as SOM inhibition keeps the apical compartment hyperpolarized despite a growing basal-to-apical exchange current.
- For low basal input, the apical compartment is released from SOM inhibition but stays below the threshold *ρ*_*a*_ of the apical gate. Beyond a certain level of apical input, VIP cells become increasingly active and inhibit SOM cells. Yet, the basal compartment is not depolarized enough to provide the first influx that would raise the apical membrane potential to *ρ*_*a*_ and open the gates of the voltage-dependent synaptic FB channels. As a result, the basal activity stays low, and the TPN remains silent.
- For coinciding basal and apical input, such initial depolarization of the apical compartment from basal influx becomes possible. While the activation pattern of the VIP and SOM interplay stays the same, now the apical compartment becomes highly active, too, and, in turn, drives the basal compartment into the high-activity regime.

In sum, these simulations show that the proposed TPN model reproduces the input-output map of a multi-compartment L5Pyr model and, thus, support its biological plausibility. The simulations further provide a dynamical computational account of a mechanistic interpretation of the underlying processes for asymmetric apical-basal coincidence detection and firing rate enhancement.

### 4.2. Learning in the TPN microcircuit via apical-basal coincidence and VIP dis-inhibition

#### 4.2.1. Context integration helps decide ambiguous input

Contextual input to a pool of TPNs influences its evidence integration via biased competition (Sect. 3). Those neurons receiving coinciding basal driving sensory input and apical context will gain an advantage in the competition and more strongly steer the pool’s population response. Yet, for the coincidence detection, learning first needs to discover which features need to be represented and with which context they frequently co-occur.

An instructive example of learning driving and contextual patterns can be drawn from experiments in visual psychophysics of orientation discrimination in the presence of flanker stimuli. There, a decision about the orientation of a central stimulus has to be made, and the orientation of laterally displaced flanker stimuli provides context for this decision. If flankers are in good continuation, i.e., placed collinearly, they facilitate the perception of a weak stimulus at the center (Polat and Sagi, 1993; Kapadia et al., 1995). In contrast, placing the flankers incongruent to the center of the configuration, i.e., rotated by 90 degrees, does not yield such facilitation (Kapadia et al., 1995).

In the present learning task a small pool of two TPNs is competing for representation of the central stimulus (Fig. 3; see Appendix C.2 for details). The two TPNs have the same cell parameters as the single TPN of Fig. 5 (Tab. A.1). Both neurons receive the same bottom-up image, as well as an image representing the contextual flanker information. Congruent flankers share the orientation of the central bar, which places them collinear with the horizontal bar but parallel to the vertical bar, so that the model has to learn from their co-occurrence which flanker orientation belongs to which bar. Kay and Phillips (1997, Experiment 2) studied a simpler task with horizontal and vertical bars of 5 × 5 pixels, in which two local processors provided each other with context (Sect. 6.2.1). Neurons enter a competition with each other via mutual recurrent input to PV interneurons. After initialization with random basal and apical synaptic efficacies (cf. Sect. 2.2.2), the model is presented with trials of different stimulus-flanker pairings, three quarters of which are congruent. During each trial a decision function evaluates the pool’s representation and decides on the network’s response to the stimulus, i.e., either “horizontal” or “vertical”, based on reading which of the two neurons responds more strongly (Sect. 2.3.2). A teacher then provides either positive or negative reward based on the correctness of the response. The reward signal becomes available to the whole local circuit by means of a volume-transmitted neuromodulator (Sect. 2.2.2). Yet, how strongly each TPN takes up this modulator is set by its coincidence tag, which grows with the apical-to-basal current during the trial. The tag thereby also sets the extent of the longterm facilitation or depression of the TPN’s synapses. Likewise, which synaptic pattern is learned depends on the neo-Hebbian factors of coinciding pre- and postsynaptic quantities as defined above (Sect. 2.2.2).

Across trials, differentiation between the TPNs happens and each neuron becomes selective to a different basal input pattern and favors different coinciding apical context (Fig. 6B). After 320 training trials, all 30 networks discriminate the unambiguous stimuli, with an average accuracy of 1.00 compared to 0.41 before training (see Fig. C.2 in the Appendix). Both the coincidence tag and the dopamine signal are needed (Fig. 6A). Without the tag, both TPNs take up the reward signal alike and approach the same basal pattern, so that 28 of the 30 networks fail to discriminate the stimuli. Without dopamine, the tag alone still builds a consistent code of stimulus and context in 29 networks, but which TPN reports which class is left to chance, with 15 correct and 14 mirrored codes (Appendix C.3). Without both factors, the networks do not learn the task. A neurometric curve can be obtained to test how the learned circuit handles different levels of uncertainty (Fig. 6C; Newsome et al., 1989; Britten et al., 1992). To this end, the model is presented with blended stimuli paired with different types of contextual evidence (similar to networks in a study by Islah et al. 2025). In the case of uninformative context, as with silenced apical input, the decisions of the model switch abruptly at the point of subjective equivalence. If decisive context for either of the learned patterns is provided, the model integrates this additional source of evidence and two regions of distinct response profiles emerge around the former point of subjective equivalence (original chance level response). For the region where context matches the predominantly represented pattern in the basal input (congruent case), the decisions become certain up to the former point of subjective equivalence (Fig. 6C). After an initial onset phase, the apical potentials reach the threshold *ρ*_*a*_ of the apical gate (Eq. 6), and contextual evidence affects the basal compartment (Fig. 6D). The activity of the TPN with a congruent representation is then amplified and wins the competition. In contrast, for the region where context mismatches the predominantly represented pattern in the basal input, the decisions follow the context up to mixtures about 0.1 away from the former point of subjective equivalence, beyond which the basal evidence prevails. Mixtures close to the point of subjective equivalence contain features that match the learned basal patterns of both TPNs, so that context is a relevant source of evidence for their disambiguation. Further away, the bottom-up evidence becomes clearer and strongly mismatches the context, which then no longer reverses the decision.

Overall, this learning task showcases how the TPN model is able to learn from purely local rules to adapt its basal and apical synapses. Learning and processing are intimately linked in the model. On the one hand, learning requires coincidence detection to form coincidence tags determining the uptake of volume-transmitted neuromodulators. On the other hand, learned patterns help with coincidence detection and biased competition to form more decisive response patterns. The model learns to perform weighted apical-basal evidence integration and generalizes its response properties to unseen ambiguous stimulus-context configurations.

#### 4.2.2. Apical-basal integration learning depends on VIP dis-inhibition

Learning in the TPN model is not just a pure function of basal and apical integration. The surrounding microcircuit of inhibitory interneurons plays a crucial role in modulating the TPN’s operating regime and response behavior. Mutual inhibition between the pool’s TPNs via their PV interneurons establishes a motif of competition that weighs the evidence for different features against each other (Sect. 3). The VIP-SOM circuit of inhibitory interneurons, in turn, regulates apical (hyper-/de-)polarization via (dis-)inhibition and, thus, indirectly impacts apical-basal coincidence detection (Fig. 5C).

In the first experiment, the static tonic input to VIP neurons was fixed at 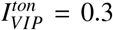(Appendix C.2). A second experiment investigates the effect of the VIP-SOM dis-inhibition on learning the orientation discrimination task. To this end, separate sets of the same 30 initial networks were trained and tested, each set at its own constant level of tonic VIP excitation or inhibition. Tonic excitation of this kind could be supplied by sub-cortical sources (Sect. 6.1.3).

Depending on the tonic input to the VIP cells, the model’s learning behavior changes substantially (Fig. 6E). Two regimes can be identified, one where learning is enabled and one where it is shut off. The transition between them spans tonic inputs from about −0.45 to 0.3, and within it, first the accuracy and then the context effect saturate (see Fig. C.6 in the Appendix). If VIP cells are depolarized, SOM inhibition on the TPN’s apical compartment is lifted and apical-basal coincidence detection can in principle take place. In this case, both TPNs could possibly establish a coincidence tag on individual trials. Yet, all networks but one still establish their overall selectivity and the response behavior of the first experiment is preserved (cf. Sect. 4.2.1 and Fig. 6E; Appendix C.3). Neutral tonic input already lies within this transition, where the networks still learn to discriminate the stimuli but the context effect at the point of subjective equivalence is weaker (0.87 instead of 1.00). A context whose initial apical efficacies are weak can then fail to open the apical gate of its TPN, so that the apical learning rule does not strengthen these efficacies and their decay erases them (Appendix C.2). If VIP cells are inhibited more strongly and shifted into a regime of hyperpolarization 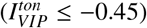 , learning is effectively stopped. In this regime, contextual input can no longer drive VIP cells enough to establish dis-inhibition of the TPN’s apical compartment. As a result, apical-basal coincidence does not take place and, hence, no coincidence tag is formed. Despite continued presentation of congruent trials, no clear selectivity emerges and response behavior stays at the level of untrained networks, with an average accuracy of 0.38 that lies below 0.5 because undecided trials count as errors (Sect. 2.3.2)^6^.

In sum, learning and the formation of selective patterns rely on the model TPN’s apical-basal coincidence mechanism and local learning terms, while the surrounding microcircuit of inhibitory interneurons further regulates and gates this process. Specific interneuron circuits, such as the VIP-SOM circuit for regulating apical (dis-)inhibition, can alter the prerequisites for coincidence detection and, thus, shut off or allow learning in the model.

## 5. Results – Simulations with the two-point neuron network model

Competition within and cooperation among TPN populations jointly give rise to processing capabilities for evidence integration in the network. We conduct model simulations that show how TPN populations cooperate to establish coherence and to propagate evidence to other parts of the network. Critically, these capabilities depend on the contextual projections between neighboring TPN populations. Yet these projection patterns can take effect far beyond the local neighborhood by means of recurrent processing.

### 5.1. Context integration and evidence propagation

In total, five network simulations with different basal and apical input configurations were conducted to understand the impact of context on a local population’s processing (Fig. 7). They use synthetic inputs to isolate the network mechanisms, as each configuration differs from another one in a single factor only. The network consists of a line of five interconnected ring attractor populations (Fig. 7A, right; cf. Fig. 4, bottom). Neurons in each ring were driven by Gaussian FF input with time-varying noise and a mode that differs per ring between 0.95*π* and 1.05*π* (Fig. 7A, left; see Tab. D.1 in the Appendix). For the different simulations, FF input was configured to represent varying degrees of ambiguity about the presence of an input feature. The Gaussian input was either parametrized as an uncertain cue (*σ* = 0.35*π*, peak amplitude 1; configurations 1, 2, 4) or as a certain cue (*σ* = 0.05*π*, peak amplitude 4; configurations 3, 5). Additionally, simulations differed depending on whether external biasing context was provided to one of the rings (configurations 4, 5). Finally, we define a *lesioned network variant* in which the contextual projections between neighboring rings are removed, i.e., their input to both the apical compartments and the VIP cells (configuration 1). This network serves as a baseline to evaluate the computational impact contributed by a network with intact contextual input integrated in the apical compartments.

#### Configuration 1: lesioned, uncertain cues

If the TPN populations compute independently of each other, each TPN population forms, by competition among its members, a bump of activity (Fig. 7B, 1). This bump represents the integrated evidence of the population’s TPNs, and its population vector encodes the most likely feature value (direction feature along the ring structure). Because bottom-up evidence is uncertain, each output peak remains of moderate activity (peak rates of about 0.08, averaged over the last 500 ms). Since TPN populations remain segregated, the evidence of each population is not integrated with that of the others. As a consequence, the population vector readouts remain noisy and incoherent.

#### Configuration 2: coupled, uncertain cues

When neurons become coupled, but activity per ring is still uncertain, the baseline pattern persists (Fig. 7B, 2). TPNs do not reach a high enough level of activity (cf. Fig. 5); hence, their output to the apical dendrites of the neighboring rings stays below the dendritic threshold *ρ*_*dend*_ required for integration (Eq. 25).

#### Configuration 3: coupled, certain cues

Once the network is presented with enough bottom-up evidence, dynamical lateral coupling sets in (Fig. 7B, 3). TPNs reach elevated levels of activity above *ρ*_*dend*_, which enables apical dendritic integration. The contextual projections excite the apical compartments of equally tuned TPNs in the neighboring rings and, via their VIP cells, release them from SOM inhibition (Sect. 3.1.2). As a consequence, apical-basal coincidence detection occurs and basal and apical evidence becomes locally integrated per ring. TPNs whose FF input coincides with the context of their neighbors are amplified and bias the competition within each ring toward the preferred directions of the neighbors. Evidence propagates among neighboring rings and their activity profiles become more similar.

#### Configuration 4: coupled, uncertain cues and bias

External contextual bias to one of the rings can also result in dynamical coupling, if paired with weak uncertain cues (Fig. 7B, 4). Contextual FB coincides with existing TPN activity and biases the uncertain competition toward the shared evidence. This coincidence again yields elevated states of TPN firing above *ρ*_*dend*_, which become apically integrated by the next neighboring ring. In turn, the neighbor’s uncertain competition is biased and the process repeats, ring by ring in steps of 22 to 112 ms, 64 ms on average (Fig. 7C, 4). Over time, the evidence among populations becomes integrated and the population vector readouts become stable and coherent. Along the coupled rings the evidence propagation results in rotating population vectors from the unbiased winner to the TPN that best represents the integrated evidence. The contextual input first releases the apical compartment of this TPN from SOM inhibition via its VIP cell, and its apical potential crosses the gate threshold *ρ*_*a*_ only after the population vector has passed the midpoint between the ring’s cue center and the bias direction (Fig. 7D; see Fig. D.3 in the Appendix).

#### Configuration 5: coupled, certain cues and bias

If biasing context is met by a network that already receives strong, certain bottom-up evidence, the picture differs (Fig. 7B, 5). The ring that locally receives both external context and context from its neighbor keeps its population vector close to its cue center and moves it by only about 8 % of the distance to the bias direction (Fig. 7C, 5). The certain bottom-up evidence to all five locations outweighs the single additional bias, and the other ring populations remain stable and centered.

In sum, these simulations show how model TPNs can (co-)operate within an interconnected network to establish meaningful contextual processing on a low-dimensional manifold and how this processing has different operating regimes depending on the sources and ambiguity of evidence. The neural fields consisting of laterally coupled TPN populations perform contextual evidence integration, establish coherence, and propagate integrated evidence about the input among neighbors. If the driving evidence remains uncertain, biasing context can help steer the population response toward directions supported by FF evidence (see Appendix D.2 for context opposite to the FF input), while certain input remains stable and largely unbiased.

### 5.2. Motion disambiguation through contextual integration

The previous sections established how the TPN model both processes information and learns from coincidence detection and how TPNs integrate evidence once they compete within pools and cooperate with neighboring pools via apical coupling. This section demonstrates how the TPN mechanisms establish evidence integration and propagation in larger networks.

Again, each location in these networks is represented by a ring population that spans the dimension of motion direction. Models with 10, 20, 40 and 100 such locations were simulated under varying conditions of a visual motion integration paradigm (Fig. 8). Visual motion integration describes the problem of integrating locally available motion signals into a coherent object-level motion estimate, a task relevant for biological vision and computer vision alike. To form useful motion estimates for visual shapes with extended outline boundaries, neural systems need to solve the *aperture problem* (Fig. 8A; Adelson and Movshon, 1982; Born and Bradley, 2005). The aperture problem arises from the nature of distributed and localized information processing due to the small receptive field (RF) sizes of cells at early processing stages in the brain. Such RFs can only measure local motion signals. The quality of such local motion signals depends on the spatio-temporal luminance gradient, and, consequently, on the structural information of the moving visual object. Two-dimensional image structures that determine an intrinsic object feature (corners) carry unambiguous motion signals about the true motion, while one-dimensional structures (edges) carry only ambiguous motion signals. Neurons within the network need to disambiguate and integrate these local motion signals to form a coherent motion representation of the object.

To investigate motion integration in light of the aperture problem, a line of TPN ring populations, one per location along the outline, was simulated with input resembling (un-)ambiguous cues from moving bars of different lengths (Fig. 8B). The network received certain, unambiguous input at the true motion direction only at the bar’s corners and uncertain, ambiguous input at the 1-D motion component along its interior. As the broad uncertain input also drives the TPNs tuned to the true motion direction, the true motion is already contained in the local signal and only needs to be disambiguated. The model shows how motion integration and disambiguation happen dynamically by propagating evidence from corner to interior locations connected along the bar’s outline (Fig. 8C,D). This process is sequential and time-consuming with the evidence propagation occurring at constant speed, such that it takes longer the further the interior locations are away from corners (Fig. 8F,G). Each of the two fronts that start at the ends of the bar advances by one ring every 167 to 220 ms, so that the time to full alignment grows by about 100 ms per ring of bar length (Fig. 8G). The fronts thus advance more slowly than the steps along the line of rings (Sect. 5.1), where the bias lies closer to the cue centers and receives more FF drive. A qualitatively similar time-dependent resolution, from the 1-D motion component to the true motion direction, has also been found in the responses of MT neurons and in smooth pursuit eye movements of monkeys viewing tilted moving bars (Pack and Born, 2001).

Locally, the same evidence propagation among neighbors is found as in the previous experiment (cf. Sect. 5.1 and Fig. 7). TPNs within each local pool compete for representation of the motion information and disambiguate their representation based on the available evidence from neighboring context (Fig. 8D). Each ring population integrates and disambiguates its representation locally but follows a stereotypical pattern (Fig. 8E) with constant propagation speed along the bar representation (Fig. 8G). At first, the ambiguous motion signal resolves the competition toward the 1-D motion component with an absolute angular error of 45^°^ against the true motion direction (Fig. 8H). Once the context provides clear enough evidence to resolve the local ambiguity, it releases the apical compartment of the neuron coding for the true motion direction from SOM inhibition via the VIP cells. The exchange current then drains less from the basal compartment of this neuron, so that the biased competition shifts toward the true motion direction until the local pool disambiguates its estimate. Shortly afterward, the neuron integrates the context apically (*v*_*a*_ *> ρ*_*a*_, Eq. 6). The resulting amplification further biases the competition, and the output of the neuron rises enough to pass the evidence on to the next ring (see Fig. D.3 in the Appendix). Eventually, motion information is disambiguated along the complete bar and yields a coherent, integrated representation (Fig. 8C).

A decoupled network variant tests whether this propagation requires the coupling between the apical and basal compartments within each TPN. In this *decoupled variant*, the exchange conductance of every TPN is set to zero (*g*_*ab*_ = 0 in Eq. 8), which removes the only route by which apical input acts on the basal compartment, while the contextual projections remain in place. No interior ring then disambiguates its representation in any of the bars, and the average angular error stays at 45^°^ (Fig. 8F, right). The local motion signals are still represented but no longer integrated into a coherent representation of object motion.

Overall, these simulations show how the local processing motifs of the TPN model yield evidence integration and propagation at the network level. Both rest on biased competition within the pools and on apical-basal coincidence detection within the TPNs. The local propagation of context yields evidence integration and a coherent representation far beyond the local interaction between immediate neighbors in the network.

## 6. Discussion

In this contribution, we present a dynamical two-point neuron (TPN) model of layer V pyramidal (L5Pyr) cell populations and their local circuit of PV, SOM, and VIP interneurons, and study it at the cell and the network level. The work makes four contributions. First, the conductance-based TPN with bidirectional apical-basal coupling reproduces the basal-apical input-output map of a multi-compartment L5Pyr model (Shai et al., 2015; Hay et al., 2011) and shows how apical amplification unfolds over time under interneuron control (Sect. 4 and Fig. 5). Second, the same coincidence mechanism serves as a structural credit signal in a local four-factor learning rule, with VIP dis-inhibition that gates whether learning takes place (Sect. 4.2 and Fig. 6). Third, networks of TPN populations with lateral apical coupling integrate context, form coherent states, and propagate evidence at constant speed, as shown for the motion aperture problem (Figs. 7, 8). This propagation fails when the basal and apical compartments of each TPN are decoupled (Fig. 8F). Fourth, a reduction relates the TPN to our earlier reduced models, i.e., modulated excitatory-inhibitory (Mod-E-I) point neurons (Bayerl and Neumann, 2004; Bouecke et al., 2011; Brosch and Neumann, 2014a,b) and a simplified TPN with thalamic gating of apical-basal integration (Schmid and Neumann, 2025), whose large-scale results on incremental grouping are summarized in Appendix E.4. The model is developed along a progression of scale, from a single TPN with its interneurons (0-D), via pools and rings of TPNs that span a feature dimension (1-D), to networks of rings that span space and feature (2-D), and to the reduced models for large-scale simulations. Rings of TPNs provide a compact population code of a feature dimension, and lines of rings a minimal spatial arrangement, so that the mechanisms of evidence integration and propagation can be studied in isolation. The model equations extend to input and feature spaces of higher dimension and more complex organization. We have so far studied two-dimensional retinotopic maps with intersecting contours in the simplified TPN (Schmid and Neumann, 2025; Appendix E.4).

### 6.1. Cell level – what the dynamical TPN microcircuit adds

Models of pyramidal cell computation span several levels of description (Appendix A.3; see Fig. A.1 in the Appendix), from biophysically detailed multi-compartment models with reconstructed (Hay et al., 2011; Shai et al., 2015) or simplified morphology (Graham et al., 2025; Graham and Kay, 2026) via reduced spiking models (Pastorelli et al., 2025; Capone et al., 2023) to abstract transfer functions (Shai et al., 2015; Kay and Phillips, 2020). Further TPN models have been formulated at the rate level, from the original local processors with contextual guidance (Phillips et al., 1995; Kay and Phillips, 1997) to units of artificial neural networks (Adeel et al., 2026, 2025; Islah et al., 2025), and at the spike level (Naud and Sprekeler, 2018; Payeur et al., 2021; Munn et al., 2023; Maryada et al., 2025). The TPN presented here is placed between reduced spiking models and transfer functions. Like the two-compartment rate units of Murayama et al. (2009) and Hertäg and Sprekeler (2020), which are likewise embedded in circuits of dendrite-targeting interneurons, it describes activity by firing rates. Beyond these, it describes both compartments by conductance-based dynamics, which resolves the time course of apical amplification and of its control by the interneurons (Fig. 5B). Moreover, its apical-to-basal current both amplifies the response and serves as the coincidence signal for learning, so that processing and learning rest on the same mechanism (Sect. 2.2).

#### 6.1.1. Are firing rates the correct level of description?

The TPN describes the average activity of a group of L5Pyr cells, and its apical gate (Eq. 6) reduces the dendritic calcium spike of BAC firing to a threshold nonlinearity (Sect. 2.1.2). Similar threshold terms have been used in rate models of dendritic plateaus and calcium spikes (Murayama et al., 2009; Yang et al., 2016; Hertäg and Sprekeler, 2020), whereas Pastorelli et al. (2025) modeled apical calcium dynamics explicitly. Together with the exchange current (Eq. 8), the gate reproduces two rate-level properties of BAC-mediated coupling. Apical input is ineffective without basal activity (Fig. 5C), and coincident input raises the gain of the basal input-output function (Larkum et al., 2004; Shai et al., 2015). The model phases in Fig. 5B can therefore be mapped onto, but not equated with, the steps of BAC firing (Sect. 4.1)^7^.

Bursts allow somatic and dendritic input streams to be multiplexed and decoded separately (Naud and Sprekeler, 2018; Friedenberger et al., 2023), and they can steer learning (Payeur et al., 2021). The TPN with its scalar output rate does not represent bursts, but bursts also raise the average firing rate. The fitted data measure the rate as the inverse of the shortest inter-spike interval (Sect. 2.3.1), so that bursts set the measured rate in the coincidence regime (Shai et al., 2015). At the population level, transient periods of high firing, termed population bursts, propagate between visual areas (Chen et al., 2022). Their trial-to-trial covariation can be captured by population-level time shifts and gains alone (Chen et al., 2022), and a population rate model expresses both of these quantities. In the model, PV inhibition of the basal compartment bounds the rates that basal input alone can reach. Within the fitted input range, high firing rates therefore indicate coincidence (Fig. 5A,C; cf. Roelfsema, 2023). A threshold on the output rate then suffices to read out coincidence, as in the simplified TPN (Fig. 2B; Schmid and Neumann, 2025). In biological cells, strong somatic drive alone can also produce high rates, and burst-sensitive readouts would disambiguate the two cases for moderate inputs (Larkum et al., 2004; Naud and Sprekeler, 2018). Firing rates are therefore an appropriate level of description for how apical-basal coincidence shapes the gain, the time course, and the learning of population responses, but not for burst multiplexing or spike timing.

#### 6.1.2. When do dynamics and interneurons matter? TPN versus Mod-E-I

Our comparison spans a sequence of reductions, from the multi-compartment model via the TPN to the Mod-E-I point neuron, in which apical input modulates the basal drive without a time course of its own (Fig. 5A; Eq. E.3). That each reduction fits the data less well is expected, but where the reduced model fails is informative. The data show two regions of activity, i.e., intermediate rates for strong basal input with weak or no tuft input and high rates once tuft input coincides even with weaker basal input. The fitted Mod-E-I transfer function (Eq. 22) lowers its basal threshold with tuft input but misses the region of intermediate rates. The composite model of Shai et al. (2015) lowers the basal threshold and raises the maximum rate with apical input and fits best (90.1 % after refitting, 87.8 % with the published parameters). The TPN model presented here ranks second (88.6 %) and reproduces both regions mechanistically, because the apical-to-basal current adds to the basal drive once the apical gate is open. The reduction assumes that the apical compartment and the VIP-SOM circuit have settled, so that their effect can be absorbed into a modulation function *f* without a time course of its own (Appendix E.1). It thereby gives up the timing of amplification, the interneuron control of the regime, and the coincidence current used for learning. Without contextual input (FB = 0), its modulation of the form FF · (1 + FB) leaves the basal drive FF unchanged, and the fitted transfer function responds to this drive with rates well below those of the multi-compartment model in the intermediate region of basal-only firing. A modulation function that lowers the basal threshold while preserving the response to basal input alone could close part of the gap in fit while keeping the efficiency of the reduced model. Candidates are the composite model (Shai et al., 2015), the regime-specific transfer functions of Graham et al. (2025), and the piecewise-planar function of Pastorelli et al. (2025).

Beyond the fit, dynamics and interneurons matter in several respects (see Tab. E.1 in the Appendix for a comparison of the TPN and the Mod-E-I model). First, the TPN captures the timing of amplification. The sequence of dis-inhibition, gate opening, and apical-to-basal current (Fig. 5B) sets response latencies and makes the outcome depend on the timing of FF and FB input. In the Mod-E-I unit, apical input modulates the basal drive directly. The sequence of basal-to-apical and apical-to-basal interaction is therefore lost (Appendix E.3). In the TPN network, a ring disambiguates its estimate once VIP and SOM cells have dis-inhibited the apical compartment of the TPN tuned to the true direction. The subsequent opening of its apical gate amplifies this TPN enough to engage the VIP cells of the next ring (Fig. 8H; see Fig. D.3 in the Appendix), so the propagation speed depends on cell-level dynamics. Second, the TPN admits context only where basal activity suffices to open the apical gate. Context that targets neurons with too little basal drive therefore leaves TPN populations unaffected (see Fig. D.1 in the Appendix), whereas the multiplicative modulation of Mod-E-I units scales even weak residual drive (cf. transfer function M2 in Kay and Phillips, 2020; see Appendix E.3). Third, the interneuron circuit controls the operating regime, e.g., via tonic input to VIP cells (Fig. 6E and Sect. 6.1.3). Fourth, the learning rule uses the apical-to-basal current as coincidence signal (Sect. 2.2.2), for which the Mod-E-I model offers only an algebraic proxy, and VIP gating of learning requires explicit VIP cells (Sect. 6.3). Fifth, only distinct interneuron types allow predictions for cell-type-specific manipulations, such as optogenetic silencing of VIP or SOM cells. Conversely, for the steady state of large networks, the Mod-E-I reduction preserves the relevant motifs at a fraction of the computational cost, as our earlier models of motion integration and incremental grouping illustrate (Bayerl and Neumann, 2004; Bouecke et al., 2011; Schmid and Neumann, 2025; Appendix E.4).

#### 6.1.3. How is the TPN operating regime controlled?

L5Pyr cells operate in several regimes depending on the strength of their input streams and on brain state (Phillips, 2023). Graham et al. (2025) distinguished apical cooperation, amplification, drive, and integration, with apical isolation as a reference mode. Graham and Kay (2026) shifted their model from apical integration to the other regimes by tonic inhibition of the tuft, the basal dendrites, or both, and by neuromodulatory down-regulation of apical calcium channels. Pastorelli et al. (2025) obtained amplification, isolation, and drive by changing parameters that stand for neuromodulation, and related them to wakefulness, deep sleep, and REM sleep. Apical drive has been proposed to underlie dreaming (Aru et al., 2020), and general anesthesia decouples the apical from the basal compartment (Suzuki and Larkum, 2020). In the TPN, apical amplification is the default regime. Apical isolation arises when VIP cells are hyperpolarized, so that SOM inhibition of the apical compartment is not lifted, which shuts off the integration of apical input and learning (Fig. 6E). It also arises when the coupling *g*_*ab*_ is too weak for basal-to-apical influx to open the gate, which prevents evidence propagation at the network level (shown for *g*_*ab*_ = 0 in Fig. 8F). Apical drive is excluded by the current parametrization, as the gate opens only after basal-to-apical influx (Fig. 5C). This design choice enforces contextual modulation in the strict sense that context alone does not drive the output (Kay and Phillips, 2020), which is a property of the modeled amplification regime rather than of cortical feedback in general. Since strong distal input alone can drive biological cells (Larkum et al., 1999), the TPN would enter the regime of apical drive if its apical compartment were tonically depolarized or its gate threshold *ρ*_*a*_ lowered, e.g., by neuromodulatory or higher-order thalamic input. Serotonergic psychedelics have been proposed to induce such a shift, by enhancing apical NMDA and calcium signaling in L5Pyr cells and glutamate release from higher-order thalamocortical terminals (Kaup et al., 2026). In the TPN, such a shift would likely let mismatching context steer population responses (cf. Appendix E.3), i.e., it would remove the robustness that the apical gate provides (Sect. 6.1.2). Apical integration theory attributes disorganization in schizophrenia to a failure of apical amplification and acute psychosis to transient apical drive (Silverstein et al., 2026). In the TPN, these states would correspond to apical isolation, e.g., by inhibitory tonic input 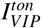 to VIP cells (Eq. 3) or a weak coupling *g*_*ab*_ (Eq. 8), and to a lowered gate threshold *ρ*_*a*_ (Eq. 6), respectively.

Compared to regime switching by cellular parameters (Pastorelli et al., 2025) or by tonic inhibition and neuromodulation (Graham and Kay, 2026), the TPN adds a circuit route via the contextual input to the VIP-SOM circuit, in line with models in which the interplay of PV, SOM, and VIP cells redistributes inhibition between soma and dendrites (Hertäg and Sprekeler, 2019). Through this circuit route, the regime can be set by identified inputs to VIP cells, such as reinforcement-related, higher-order thalamic, and top-down frontal signals (Pi et al., 2013; Williams and Holtmaat, 2019; Anastasiades et al., 2021; Zhang et al., 2014). This form of regime control relates to the gating of dendritic integration by task-related control signals (Yang et al., 2016; Halassa and Kastner, 2017; Schmid and Neumann, 2025). Neuromodulatory signals could likewise set the regime through the VIP-SOM circuit by supplying the tonic excitation of VIP cells that enables learning in Fig. 6E (Khan and Hofer, 2018). Since a constant input current acts like a shift of the leak reversal potential (Eq. 3), tonic input is excitatory or inhibitory only relative to the resting potential of the VIP cells at 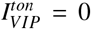. If this resting potential already reflects tonic excitation, inhibitory tonic input stands for its withdrawal, which can lead to apical isolation. Conversely, if it reflects tonic inhibition, excitatory tonic input stands for its lifting, which can enable apical amplification. In sum, the regime of the TPN is controlled by its contextual input, by tonic drive to the VIP-SOM circuit, and by cellular parameters such as the coupling *g*_*ab*_ (Eq. 8) and the gate threshold *ρ*_*a*_ (Eq. 6).

#### 6.1.4. How does the TPN relate to other models of dendrite-targeting interneuron circuits?

Rate models that combine dendritic compartments with PV, SOM, and VIP interneurons have explained pathway-specific gating (Yang et al., 2016), negative prediction-error responses (Hertäg and Sprekeler, 2020), and the layer-specific control of dendritic inhibition by NDNF cells (Naumann et al., 2025). The TPN shares their canonical motif, i.e., dendrite-targeting SOM cells that are dis-inhibited by VIP cells receiving top-down input (Wang et al., 2004; Pi et al., 2013; Zhang et al., 2014), but uses it for a different computation. In the model of Hertäg and Sprekeler (2020), SOM cells are driven by bottom-up input and cancel the top-down prediction at the dendrite when prediction and sensory input coincide. In the TPN, SOM cells are driven by the pyramidal output and silenced by VIP cells driven by contextual input, so that coincidence amplifies the response. The same circuit can thus implement subtractive cancellation or asymmetric amplification, depending on what drives the SOM cells and on the relative strength of top-down input to VIP and SOM cells. This is consistent with the idea of a division of labor between prediction-error neurons in layer 2/3 and layer 5/6 neurons that integrate top-down and bottom-up input positively (Jordan and Keller, 2020). Yet, unexpected stimuli also amplify the most selective layer 2/3 neurons, when VIP cells release the excitatory pulvinar input to V1 from feedforward inhibition by SOM cells (Furutachi et al., 2024). Whether predictive inference in cortex relies on such cancellation is debated, as it has also been argued to rest on the apical amplification of predicted signals without subtractive coding (Marvan and Phillips, 2024). This latter view is consistent with biased competition in the TPN. In contrast to pathway gating by a separate control input (Yang et al., 2016), contextual input in the TPN both targets the apical compartment and recruits its own dis-inhibition.

#### 6.1.5. What does partial information decomposition reveal about the asymmetry?

Partial information decomposition (PID) separates the information that an output carries *uniquely* about each input, *redundantly* (shared) about both, and only about their combination, i.e., *synergistically* (Williams and Beer, 2010; Wibral et al., 2017). We fitted the input-output map of the TPN to the firing rates of the multi-compartment model of Shai et al. (2015). Kay and Phillips (2020) applied PID to published simulation data of the same model and binarized its output into bursts and no bursts. Of the 0.59 bit of joint mutual information, 20 % was *unique* to the basal input, almost none was *unique* to the apical input, 27 % was *shared*, and 53 % was *synergistic*. The basal input thus drives the cell and the apical input amplifies it. With non-negative inputs, however, a transfer function of the Mod-E-I form FF · (1 + FB) also satisfies all criteria of contextual modulation (transfer function M2 in Kay and Phillips, 2020) and is purely modulatory (Kay and Graham, 2026), so that these criteria alone do not separate the TPN from the Mod-E-I model. Graham et al. (2025) showed that the decomposition depends on the operating regime, with unique basal information under apical amplification and unique apical information under apical drive, and targeted tonic inhibition shifted it toward these profiles (Graham and Kay, 2026). In the TPN, the interneurons are recruited by the inputs and the output of the cell, so that the shares of unique, shared, and synergistic information are expected to change with the input and over time. Applied to the TPN in its different regimes, PID could therefore quantify how the interneuron circuit moves the TPN between regimes and how these regimes differ from the Mod-E-I model. The continuous temporal interplay of the two input streams, in turn, would require extensions to information dynamics, such as integrated information decomposition (Mediano et al., 2025). Furthermore, PID could also relate the apical learning rule to learning goals defined over information components, such as coherent infomax (Kay and Phillips, 2011; Wibral et al., 2017).

### 6.2. Learning with apical-basal coincidence

The apical-to-basal current that amplifies a TPN during coincidence also forms its coincidence tag. This tag sets the neuron’s uptake of a dopamine-like reward signal (Sect. 2.2.2). The VIP-SOM circuit gates basal and apical plasticity through apical dis-inhibition, and the trained pool generalizes its responses to ambiguous stimuli that were not part of training (Fig. 6C).

#### 6.2.1. How does coincidence-tagged learning relate to other learning rules for TPNs?

Several learning rules exist that exploit the segregation of basal and apical input. Kay and Phillips (1997) derived learning rules by which local processors maximize the information that their output shares with both their receptive field and their context. In a simpler version of the present task, two such processors served as each other’s context and learned to signal the bar whose sign was correlated across their inputs, without a reward signal (their Experiment 2). In error-based schemes, apical input instructs basal plasticity via dendritic prediction errors (Sacramento et al., 2018) or via bursts that signal the sign of plasticity (Payeur et al., 2021). Capone et al. (2023) proposed a target-based rule, in which apical input that coincides with somatic spiking triggers bursts that instruct local plasticity. They also showed that context delivered apically is more robust than the same context delivered basally, which supports the segregation of input streams assumed here. Baronig and Legenstein (2024) derived a local apical rule that associates context with basal activity, gated by back-propagating action potentials and dendritic NMDA and calcium spikes. The apical rule of the rate-based TPN proposed here pursues a similar goal, namely to find context that co-occurs with the neuron’s own activity. Its gating is implicit, as the rectified apical potential that enters the rule becomes positive only with basal-to-apical influx. In the TPN, pooled PV inhibition takes the role that the k-winner-take-all selection of basal activity plays in the network of Baronig & Legenstein. The basal rule of the TPN differs from these proposals in its use of a global reward signal. It extends three-factor rules with eligibility traces (Izhikevich, 2007; Frémaux and Gerstner, 2016; Gerstner et al., 2018) by a neuron-specific tag for structural credit assignment, related to attentional feedback in attention-gated reinforcement learning (Roelfsema and van Ooyen, 2005) and in the framework of Roelfsema and Holtmaat (2018). Apical input therefore acts as context with amplifying characteristics rather than as a teacher. It selects which neurons learn by enabling the coincidence that forms the tag, whereas the pattern to be learned follows from the Hebbian terms and the direction of change from the reward signal^8^. Without the reward signal, the tag alone still builds a consistent code of stimulus and context, but which neuron reports which class is then left to chance (Fig. 6A). The TPN thus assigns neuron-specific credit without an explicit target or error signal in the apical compartment. Apical dendrites can nevertheless convey instructive signals, such as the target signals that drive behavioral timescale plasticity in the hippocampus (Wilmerding et al., 2026). In L5Pyr cells of the retrosplenial cortex, apical activity relative to somatic activity carries neuron-specific signals during learning, whose sign depends on the role of the neuron in the task and whose disruption impairs learning (Francioni et al., 2026). Its counterpart in the TPN is the excess of the apical over the basal potential. This excess drives the apical-to-basal current that forms the tag (Eq. 15). The tag in turn sets how much the neuron learns, whereas the sign of plasticity follows from the global reward signal. In sum, coincidence-tagged learning shares the local association of context with other apical rules, but apical input selects which neurons learn instead of instructing them.

#### 6.2.2. Why should interneurons gate learning?

Hyperpolarizing VIP cells prevented the TPN pool from learning the task, because SOM inhibition then kept the apical compartment below the gate threshold and no coincidence tag could form (Fig. 6E). This agrees with evidence that dis-inhibition gates plasticity. Higher-order thalamic input gates long-term potentiation in layer 2/3 pyramidal cells via VIP-mediated dis-inhibition (Williams and Holtmaat, 2019). Dendritic inhibition shifts plasticity from potentiation to depression in the model of Yang et al. (2016), and reward-driven VIP dis-inhibition instructs lasting changes of sensory representations in the model of Wilmes and Clopath (2019). On a developmental time scale, the maturation of inhibitory circuits sets the critical period for plasticity in visual cortex (Hensch, 2005). Tags formed by feedback-evoked dendritic calcium events, directly or via VIP-mediated dis-inhibition, that make synapses eligible for reward-steered plasticity have been proposed by Roelfsema and Holtmaat (2018) and Khan and Hofer (2018). The TPN implements these proposals in a circuit that is constrained by the input-output map of a multi-compartment L5Pyr model. In the TPN, contextual input itself drives the VIP cells, and VIP gates learning through the coincidence tag, i.e., through the mechanism that also controls processing. The model predicts that suppressing VIP activity during training prevents the learning of tasks that rely on contextual amplification. In sum, interneuron gating restricts learning to the states in which context is integrated, and inputs to VIP cells can thereby switch learning on and off.

### 6.3. Learning in large-scale models

In its current form, TPN learning acquires basal feature selectivity and apical context associations jointly from local rules and a global reward in a pool of two TPNs (Sect. 4.2). The same apical rule could learn the lateral and feedback connections between populations, as it finds context that co-occurs with the neuron’s own activity. Such learning becomes necessary for distributed codes, in which similarly tuned partners are not known in advance. The two learning rules require neither feedback weights that mirror the forward weights nor neuron-specific error signals in the apical compartment (Sect. 6.2.1), and task goals enter only through the reward. A scalar reward, however, reaches all neurons alike, whether or not they took part in the selected response (Roelfsema and van Ooyen, 2005). Rules that rely on it alone have to infer the contribution of each neuron from the covariation of its activity with the reward over many trials (Williams, 1992), which is expected to become less efficient as networks grow (Roelfsema and van Ooyen, 2005). This issue can be mitigated by structural credit tags that restrict plasticity to the neurons that contributed most to the selected action (Roelfsema and Holtmaat, 2018). The TPN forms such tags from apical-basal coincidence (Eqs. 12, 14, and 15). RELEARNN, a four-factor rule in which a reward-prediction error and an action-related feedback signal gate Hebbian plasticity, trains feedforward, lateral, and feedback connections of recurrent networks to link and trace contours (Brosch et al., 2015) and to perform multi-step visual routines (Mollard et al., 2024). This suggests that rules of this kind scale beyond small pools, although RELEARNN derives its tag from a dedicated credit-assignment shadow network with reciprocal connections, rather than from the apical-basal coincidence of the neuron itself. For large-scale simulations, the reduced models (Fig. 2B,C) are the natural substrate, with the modulatory term 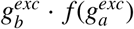 (Eq. E.3) as an algebraic proxy of the coincidence signal. To reproduce the learning dynamics of the TPN, this proxy needs (i) a slow trace, (ii) a gate on *f* that stands in for the VIP-SOM control, as provided by the thalamic gating of the simplified TPN (Schmid and Neumann, 2025), and (iii) a surrogate of the apical potential for the apical rule^9^.

### 6.4. Network level – from local amplification to global coherence

The network model arranges TPN populations as ring attractor networks (RANs) along a feature dimension and couples equally tuned TPNs of neighboring populations through their apical compartments and VIP cells (Fig. 4). Pooled PV inhibition within each ring together with asymmetric FF-FB integration implements biased competition (Desimone and Duncan, 1995), and lateral apical coupling lets populations cooperate (Figs. 7, 8). RANs (Ben-Yishai et al., 1995) impose a localistic code beyond the more general assumption that population activity lies on low-dimensional manifolds (Khona and Fiete, 2022), and this localistic code makes it easy to prescribe the lateral apical connections (Tab. 2). The guiding principles, i.e., normalized pools within populations and lateral coupling between similarly tuned neurons, do not depend on this choice, but for distributed codes the coupling has to be learned (Sect. 6.3).

#### 6.4.1. What do the network simulations add beyond single-cell contextual modulation?

While apical modulation of single cells is well established (Larkum, 2013; Kay and Phillips, 2020; Phillips, 2023), the network simulations show how such modulation, combined with competition, determines the state of populations and of the network. In a small network of compartmental model cells, Graham et al. (2025) already showed that lateral excitation of the apical tufts often converts single spikes into bursts. Three properties emerge that go beyond a single-cell transfer function. First, the effect of context depends on the certainty of the driving evidence. Biasing context steered populations with uncertain input but hardly changed populations with certain input (Fig. 7B, configurations 4 and 5), and the trained TPN pool showed the largest context effect near its point of subjective equivalence (Fig. 6C). This is the population-level counterpart of the criterion that context strengthens the transmission of information about the drive when the drive is weak, whereas the drive alone suffices for this transmission (Kay and Phillips, 2020). In line with this, collinear flankers facilitate the responses of single V1 neurons mainly to near-threshold stimuli (Polat et al., 1998), and feedback from area MT has its strongest effects for stimuli of low salience (Hupé et al., 1998). Second, single-cell amplification combined with pool competition can shift population selectivity (Fig. 7C,D). For single cells, Graham et al. (2025) found that apical amplification preserves basal orientation tuning under mismatched context, whereas apical integration averages the selectivity of both inputs. In the network, context still moves the population estimate in the amplification regime, because it biases the competition within the population (Desimone and Duncan, 1995). Third, local coupling yields global coherence by sequential propagation (Fig. 8C). In the TPN network, the time to full alignment grows linearly with bar length (Fig. 8F,G). In the simplified TPN, the onset latencies of the response enhancement along a traced contour likewise increase steadily with the distance from the attentional seed (Schmid and Neumann, 2025). There, FF-FB coupling across a scale space speeds up the propagation where distractors are farther away. The TPN network presented here couples its populations laterally at a single spatial scale and therefore propagates at constant speed.

Beyond the certainty of the driving input, propagation can be steered by task demands. In the simplified TPN (Fig. 2B; Schmid and Neumann, 2025), higher-order thalamic input gates apical-basal integration, so that an attentional seed determines which contour should become coherently integrated (Appendix E.4). There, we suggested that this gate might be mediated by apical (dis-)inhibitory circuitry. The VIP-SOM circuit of the full TPN model provides a candidate mechanism at such a finer level, as it controls apical integration and learning (Figs. 5C, 6E) and receives higher-order thalamic input in somatosensory cortex (Williams and Holtmaat, 2019). As a testable prediction, silencing VIP cells or their thalamic input should then impair incremental grouping. In line with this, inhibiting VIP cells reduces figure-ground modulation in mouse V1 (Kirchberger et al., 2021). In sum, beyond single-cell modulation, the network simulations show certainty-dependent context effects, shifts of population selectivity, and global coherence, and they suggest that task demands could steer the propagation of coherence via the VIP-SOM circuit.

#### 6.4.2. How do the network dynamics relate to physiological and behavioral data?

In macaque MT, responses to moving bars first signal the motion component perpendicular to the bar and encode the true direction within 150 ms of motion onset (Pack and Born, 2001). Within the receptive fields of MT neurons, end-stopped V1 input can account for this transition, and the deviation of MT tuning curves does not change significantly with bar length (Tsui et al., 2010). The initial deviation of pursuit eye movements toward the perpendicular component, however, increases with bar length (Pack and Born, 2001; Born et al., 2006), consistent with a time-consuming integration of the motion signals from the bar ends. For contours that extend beyond the reach of end-stopped inputs, the TPN network reproduces the temporal sequence of component and true direction. Its absolute time scale (Fig. 8G) was not fitted to these data (Sect. 6.5.1). Our earlier Mod-E-I networks show similar propagation (Bayerl and Neumann, 2004; Bouecke et al., 2011), and the TPN network shows that it follows from the cell-level coincidence mechanism (Fig. 8F) and its interneuron control.

Anesthesia, which decouples the apical from the basal compartment of L5Pyr cells in mouse cortex (Suzuki and Larkum, 2020), offers a test of this integration mechanism. In monkeys anesthetized with isoflurane, MT neurons respond vigorously and keep their direction tuning, but their integration of local motion signals into a global motion estimate is markedly impaired (Pack et al., 2001), and figure-ground modulation in V1 is suppressed (Lamme et al., 1998). Under opioid anesthesia, MT neurons have nevertheless been found to signal pattern motion (Smith et al., 2005), so that the effect may depend on the anesthetic. In the decoupled network variant (*g*_*ab*_ = 0), the local motion signals remain represented but are no longer integrated into a coherent representation of object motion (Fig. 8F). The model therefore predicts that anesthetics that decouple the compartments of L5Pyr cells impair the propagation of contextual signals across receptive fields, while local feature tuning is preserved. In sum, the network dynamics qualitatively match the transition from component to true direction in MT and in pursuit eye movements, and the decoupled variant matches the impaired motion integration under isoflurane anesthesia.

### 6.5. Limitations and outlook

#### 6.5.1. Scope and model limitations

The TPN is designed to capture the properties that we consider critical for contextual amplification, learning, and coherence at the level of population rates, and to explain them mechanistically and dynamically. Several simplifications follow from this aim, and they delimit the claims that the model supports. As a population rate model, the TPN does not represent burst multiplexing or spike timing (Sect. 6.1.1). Its apical compartment has a single threshold, whereas tuft dendrites contain many branches with local NMDA spikes (Larkum et al., 2009) that may associate context branch by branch (Baronig and Legenstein, 2024). NMDA-driven dendritic modulation can implement task-specific modulation of feedforward processing (Wybo et al., 2023), and context can also be mapped onto the apical compartment by a learned nonlinear function (Islah et al., 2025). The gating of individual afferents in the network model (Eq. 25) is a first step in this direction. It suffices for the generic simulations presented here, whereas branch-specific integration would add to it when many heterogeneous sources of context have to be arbitrated. The proposed interneuron circuit, which controls the basal and apical activation of the pyramidal cell compartments, is reduced to the connections needed for the computations studied (Sect. 6.1.4) and omits further types such as NDNF cells, which compete with SOM cells for dendritic inhibition (Schuman et al., 2019; Naumann et al., 2025). It also omits further connections, such as the inhibition of PV cells by SOM and PV cells (Pfeffer et al., 2013).

Other limitations concern the validation. The TPN was compared with the multi-compartment model of Shai et al., which covers one cell type and was itself constrained by in vitro recordings (Hay et al., 2011; Shai et al., 2015). The internal parameters of the TPN were tuned in a non-exhaustive search against the input-output map, but not against response time courses (Appendix B.1). The time courses in Figs. 5B, 7, and 8 are therefore qualitative, and the absolute propagation speed is not a prediction of the model. To isolate the core mechanisms of contextual evidence integration, the network simulations use synthetic inputs along one spatial and one feature dimension and populations that are uniformly spaced at a single scale (Sect. 3.3.1). Earlier models in the lineage of the TPN have processed real image sequences on two-dimensional maps and reproduced findings on motion integration, with rate-based Mod-E-I units (Bayerl and Neumann, 2004; Bouecke et al., 2011) and with spiking ones (Löhr et al., 2019). The TPN network supplies the cell and circuit mechanisms for such models and can thereby relate these findings to the dynamics of identified cell types.

#### 6.5.2. Implications beyond biological systems

The model realizes cooperation for evidence integration with generic motifs, i.e., asymmetric FF-FB integration, pooled inhibition, and lateral coupling of similarly tuned units. Coherent states of this kind have been proposed to underlie feature binding (Roelfsema, 2023) and consciousness (Storm et al., 2024), whereas binding remains a challenge for artificial neural networks (Greff et al., 2020). Grouping over long distances, as in tracing a path between two points, is hard for feedforward networks, and such tasks have become benchmarks for long-range dependencies (Linsley et al., 2018; Tay et al., 2021). They have been addressed by learned horizontal recurrence (Linsley et al., 2018), attention spreading (Adeli et al., 2023), learned visual routines (Mollard et al., 2024), recurrent dynamics with neural synchronization (Darlow et al., 2025), and learned multiplicative top-down and lateral gating (Salehi et al., 2026). These mechanisms resemble the evidence propagation studied here. With such propagation, the simplified TPN traced target contours in example stimuli of the Pathfinder task without training (Schmid and Neumann, 2025). TPN- and dendrite-inspired units have been used in artificial neural networks for multi-modal fusion (Adeel et al., 2026), multi-task learning with sparse representations (Iyer et al., 2022), efficient reinforcement learning (Adeel et al., 2025), and combining context with ambiguous input (Islah et al., 2025). Reduced transfer functions, such as the Mod-E-I modulation function or the piecewise-planar function of Pastorelli et al. (2025), offer units for such networks, and the TPN specifies which of their properties derive from cell and circuit mechanisms. Three leads follow for such architectures. First, the same coincidence signal amplifies processing and gates plasticity in the TPN, which restricts learning to the units that take part in a coherent representation. Units built on this principle could learn online during inference rather than in a separate training phase. Second, asymmetric gating of apical input prevents context from creating responses without driving evidence and from overriding certain evidence (Fig. 7B, configuration 5; Appendix D.2), which may make networks more robust to misleading context (cf. Islah et al., 2025). Third, local lateral propagation remains coherent over long distances with a time to full alignment that grows only linearly with distance (Fig. 8G; see Appendix D.3 for bars of 500 and 1000 rings). Such recurrent propagation outperforms deep feedforward networks on connectedness tasks when enough processing time is available, whereas feedforward solutions win under tight time limits (Larsen and Druckmann, 2022), a trade-off that propagation across coarser spatial scales may loosen (Schmid and Neumann, 2025). Thus, such lateral propagation offers a local alternative to global interactions for long-range dependencies.

### 6.6. Conclusion

We proposed a TPN model that links the cell, circuit, and network levels of contextual integration. It reproduces the asymmetric input-output map of a multi-compartment L5Pyr model with a mechanism whose time course and operating regime are controlled by PV, SOM, and VIP interneurons. The same coincidence mechanism tags neurons for neuromodulator-driven four-factor learning, and lateral apical coupling between TPN populations turns local amplification into global coherence. Reduced model variants carry these principles to large-scale simulations, and the TPN specifies when dynamics and interneurons have to be modeled explicitly. The model predicts that (i) strong VIP silencing abolishes the contextual amplification of L5Pyr cells and the learning that depends on it, (ii) context biases population responses most when driving evidence is uncertain, and (iii) the source of SOM drive distinguishes amplifying from canceling circuits, a distinction that follows from the comparison with prediction-error circuits (Sect. 6.1.4). Taken together, the TPN shows how pyramidal cells and their interneurons let context amplify but not create sensory evidence, select the neurons that learn, and turn local amplification into coherent network states. The TPN thus provides a biologically grounded reference from which, as we show, established reduced units for large-scale and artificial neural networks can be derived, together with an account of what such reductions retain and what they lose.

## CRediT authorship contribution statement

**Conceptualization:** Daniel Schmid, Heiko Neumann.

**Data curation:** Daniel Schmid.

**Formal analysis:** Daniel Schmid.

**Funding acquisition:** Heiko Neumann.

**Investigation:** Daniel Schmid.

**Methodology:** Daniel Schmid, Heiko Neumann.

**Project administration:** Daniel Schmid, Heiko Neumann.

**Resources:** Heiko Neumann.

**Software:** Daniel Schmid.

**Supervision:** Heiko Neumann.

**Validation:** Daniel Schmid.

**Visualization:** Daniel Schmid, Heiko Neumann.

**Writing - original draft:** Daniel Schmid, Heiko Neumann.

**Writing - review & editing:** Daniel Schmid, Heiko Neumann.

## Acknowledgments

The authors thank Christian Jarvers for helpful discussions and input during early stages of developing the model. The authors thank the anonymous reviewers for their helpful comments, which improved the quality of the manuscript. The authors acknowledge support by the state of Baden-Württemberg through bwHPC.

## Declaration of competing interest

The authors declare that no competing interests exist.

## Funding

This research did not receive any specific grant from funding agencies in the public, commercial, or not-for-profit sectors.

## Data availability

Code for the two-point model, the ring attractor model and for the simulation and the analyses will be provided via an open repository upon acceptance.

## Declaration of generative AI and AI-assisted technologies in the manuscript preparation process

During the preparation of this work, the authors used the services of Anthropic Claude and additionally Mam-mouth.ai to access AI models from OpenAI ChatGPT, Google Gemini, Anthropic Claude, and Mistral AI in order to assist with performing the investigation and to improve the writing of this manuscript. They also used the tools Claude Code and GitHub Copilot Pro to help implement the code used for this investigation, and the models Claude Opus 5.5 and Claude Fable 5 (Anthropic) to improve existing visualizations in the manuscript. After using these tools and services, the authors reviewed and edited the content as needed and take full responsibility for the content of the published article.

## Appendix

The Appendix gives the details of the methods and additional results in five parts, which follow the main text from the single TPN to its learning, its networks and its reduced versions.

### Appendix A. Two-point neuron model

#### A.1 Model parameters and time constants

Table A.1 lists the parameters of the TPN equations (Eqs. 1–10) in the three settings of this study: the single TPN of Fig. 5, the pool of two TPNs of the flanker task (Fig. 6) and the ring networks of Figs. 7 and 8. The parameters of the single TPN were tuned against the data of Shai et al. during model development, first manually and then in a refinement with the help of AI coding agents (Appendix B.1). In the comparison with the other models, only the map between synapse numbers and model inputs and outputs of the TPN was fitted, which relates its input-output space to these data (Sect. 2.3.1). The three settings share these cell parameters. They differ only in the connectivity of the TPNs and of their interneurons, in the tonic input to the VIP cells, and in the dendritic threshold on the contextual afferents, which only the ring networks have. In the flanker pool and the ring networks, the PV cells receive the pooled rates of all TPNs of a population instead of the TPN’s own rate. The weights of these pooled projections were adjusted for TPNs that compete within a population. Regime boundaries measured for the single TPN therefore do not transfer one to one to a TPN within a population. The pool of the flanker task is a population of *N* = 2 TPNs, in which the Gaussian kernels reduce to the identity. Its VIP cells receive an excitatory tonic input 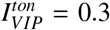, which depolarizes them toward their output threshold and thus facilitates the dis-inhibition of the apical compartment by contextual input (Appendix C.2). For the ring networks, a few connection parameters were set in addition. These are the width of the recurrent excitation, the weights of the pooled projections onto the PV and VIP cells, and the weight and the dendritic threshold of the contextual projections.

All time constants are those of rate units that represent the mean activity of a local group of neurons. The basal compartment relaxes with *τ*_*b*_ = 10 ms, on the order of the response times of cortical neurons (Koch et al., 1996). The apical compartment integrates its input 3.5 times more slowly (*τ*_*a*_ = 35 ms), in line with the duration of NMDA spikes in tuft dendrites (Larkum et al., 2009; Antic et al., 2010), so that the apical potential needs tens of milliseconds to rise above the gate threshold *ρ*_*a*_. Conversely, an apical depolarization also persists for longer once its input ceases. The interneurons follow their inputs faster than both compartments of the TPN (*τ*_*q*_ = 1 ms). Together with their leak conductances, their relaxation times 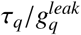 are 1.7 ms (PV), 5 ms (SOM) and 1 ms (VIP). Fast interneurons are also assumed in rate models of the PV-SOM-VIP circuit, in which the interneuron time constants resemble fast GABA_*A*_ kinetics (2 ms against 60 ms for the pyramidal cells; Hertäg and Sprekeler, 2020). The three settings share these time constants, and all times of this study are given in milliseconds.

#### A.2. Numerical methods and software

The simulations of the three settings were run with different integrators (Tab. A.2).

All simulations and analyses were implemented in Python (3.12). The input-output maps of the single TPN were integrated with NumPy (Harris et al., 2020) (2.5). Its time courses and the ring networks were integrated with JAX (Bradbury et al., 2018) (0.11.0) and diffrax (Kidger, 2021) (0.7.2, with Equinox 0.13.8). The ring networks were, in addition, set up with Hydra (1.3.6) for their configurations. The flanker pool was integrated with NumPy and a right-hand side compiled with Numba (Lam et al., 2015) (0.67.0). The fits used NumPy and SciPy (Virtanen et al., 2020) (1.18.0). All plots were drawn with Matplotlib (Hunter, 2007) (3.11.1). Parts of the code were written with the help of AI coding assistants (Claude Code, GitHub Copilot) and checked by the authors. All random numbers (initial states, input noise, initial efficacies, starting points of the fits) were drawn from seeded generators, so that every result can be reproduced exactly.

#### A.3. Relation to the biophysically detailed BAC mechanism

Coincidence detection in L5Pyr cells can be mediated by backpropagation-activated calcium spike firing (BAC; Fig. 1B; Larkum et al., 1999; Larkum, 2013). There, BAC ensures a causal ordering of basal-to-apical and apical-to-basal depolarization. In biological cells BAC requires intricate ion-specific depolarization dynamics and the propagation of spikes between dendritic sites. A sodium spike back-propagates from the soma into the dendrites (Na^+^ from basal to apical). If it coincides with tuft input within a few milliseconds, it lowers the threshold of a dendritic Ca^2+^ spike, which is generated by voltage-gated calcium channels (Larkum et al., 1999). Its plateau, together with NMDA-receptor-mediated tuft spikes (Larkum et al., 2009), serves as a temporal aggregation window for detecting coincidence. Then, coincidence is signaled back to the soma (Ca^2+^ spike from apical to basal) and results in a spike burst at the axon.

**Table A.1:**
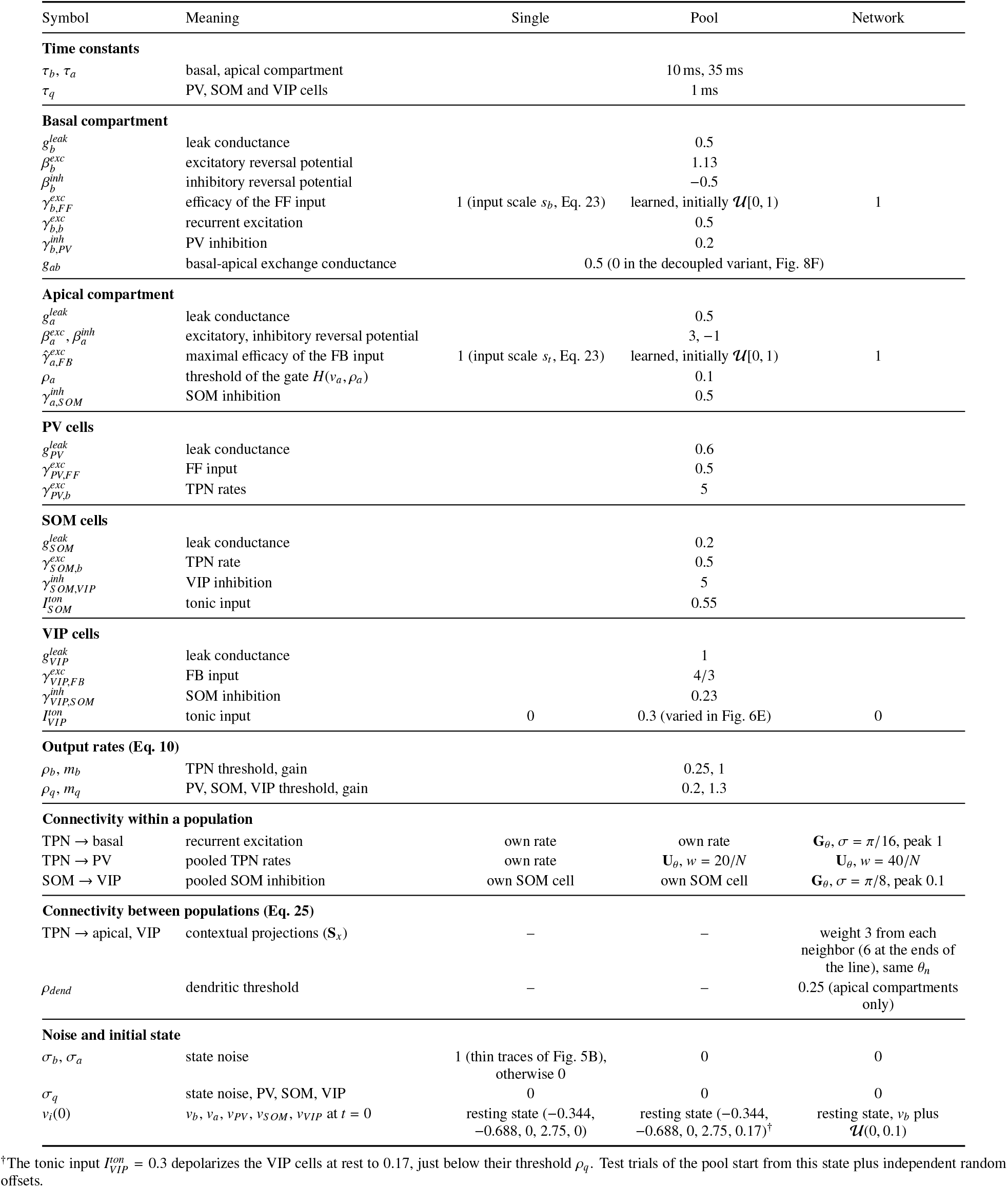
Parameters of the TPN (Eqs. 1–10). The columns give the values for the single TPN of Fig. 5 (Single), for the two TPNs of the flanker task (Pool, Fig. 6) and for the TPNs of the ring populations of Figs. 7 and 8, with *N* = 200 TPNs per ring (Network). A value that spans several columns holds for each of them. Potentials, conductances, rates and inputs are dimensionless, and the leak currents have the reversal potential 0. Kernels are named as in Tab. 2, and all projections not listed under connectivity are local (Id). Since the weights *w* of the uniform kernels scale with 1*/N*, the PV cells receive a multiple of the mean rate of their population.

**Figure A.1:**
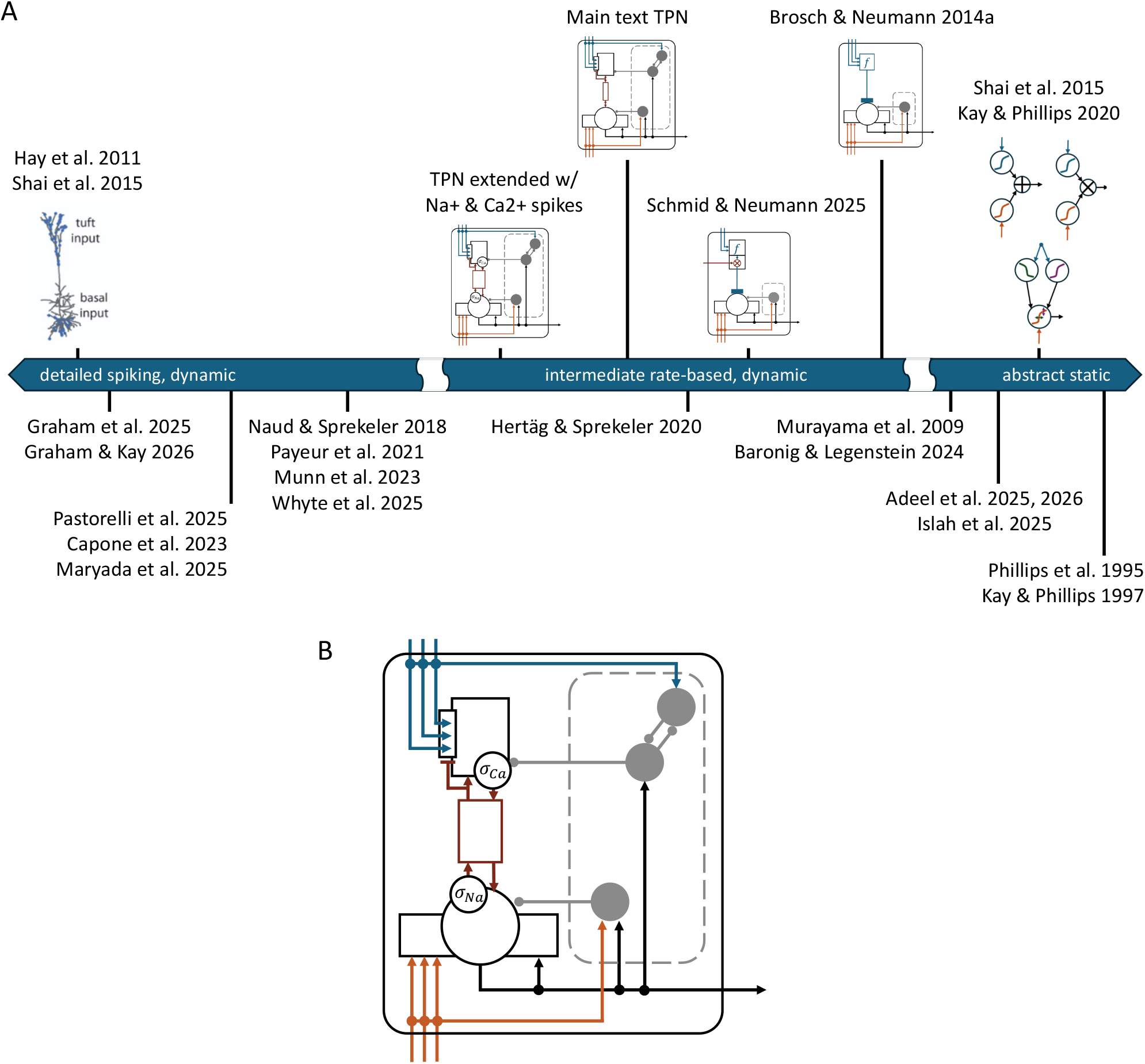
Classification of models of pyramidal cell computation and schematic depiction of an extended TPN variant with sodium- and calcium-spike-based BAC. (A) Relation of the TPN model to other models of pyramidal cell computation arranged by biophysical detail. Different models exist capturing the underlying processes of pyramidal cells either on a detailed, spike-based level (left; Hay et al., 2011; Shai et al., 2015; Graham et al., 2025; Graham and Kay, 2026; Pastorelli et al., 2025; Capone et al., 2023; Maryada et al., 2025; Naud and Sprekeler, 2018; Payeur et al., 2021; Munn et al., 2023; Whyte et al., 2025) or on an abstract, static level (right; Baronig and Legenstein, 2024; Adeel et al., 2025, 2026; Islah et al., 2025; Shai et al., 2015; Kay and Phillips, 2020; Phillips et al., 1995; Kay and Phillips, 1997). The presented model family falls onto an intermediate position along this axis capturing dynamical properties of the underlying mechanisms while abstracting away many biological details, and its simplified versions extend toward the abstract end (Schmid and Neumann, 2025; Brosch and Neumann, 2014a). Two-compartment rate units in circuits of dendrite-targeting interneurons share this intermediate position (Hertäg and Sprekeler, 2020) or, analyzed in their steady state, lie toward the static end (Murayama et al., 2009). Models with a schematic stand above the axis, and models named only by their references below it. (B) Extended variant of the presented TPN model with nonlinear exchange coupling between basal and apical compartments. Nonlinearities *σ*_*Na*_ and *σ*_*Ca*_ capture proportions of neurons within the neural mass model currently signaling bottom-up evidence to apical compartments via sodium (Na^+^) spikes or coincidence with top-down evidence to basal compartments via calcium (Ca^2+^) spikes. Multi-compartment pyramidal cell schematic in A reproduced from Shai et al. (2015) under the CC BY 4.0 license (https://creativecommons.org/licenses/by/4.0/).

**Table A.2:** Numerical integration. The fits to the multi-compartment model (Sect. 2.3.1) need no integration, as they minimize the squared error over the stored model outputs (double precision).

| Simulation | Integrator | Time step | Precision |
| --- | --- | --- | --- |
| Single TPN, input-output map (Fig. 5A,C) | explicit Euler until the state settles | 0.05 ms | double |
| Single TPN, time courses (Fig. 5B) | explicit Euler, for the noisy traces<br>Euler–Maruyama | 0.1 ms | single |
| Flanker pool (Fig. 6) | exponential Runge–Kutta<sup>a</sup> | 0.2 ms | double |
| Ring networks (Figs. 7, 8) | Dormand–Prince 5(4) (Dormand and Prince, 1980)<sup>b</sup> | adaptive | single |

<sup>a</sup>Strang splitting (Strang, 1968): the potentials of PV, SOM and VIP cells and the coincidence tag, whose equations are linear in their own state, advance by exact exponential half steps around a classical fourth-order Runge–Kutta step of all other variables. <sup>b</sup>Adaptive step size with PID control, relative and absolute tolerance $10^{-5}$ .

The presented TPN model abstracts away from many of these biological details, such as action potentials, bursts and voltage-gated calcium channels, while still capturing the implied computational logic and its dynamics. The TPN model represents a dynamical neural mass of spiking pyramidal cells and their rate-based activity level (Fig. A.1A, cf. Sect. 2 for details). The model rests on three main functional components that constitute apical-basal coincidence detection, namely

i. compartments integrate different streams of information separately,
ii. compartments exchange information about the state of their input, and
iii. in a modulatory regime, basal activation is a prerequisite to establish a subsequent apical plateau.

The model implements this logic while not spending additional detail on spike-based dynamics or receptor-specific dynamics. The two-point characteristic of the TPN with its separate compartments forms the basis for anatomically distinct spatio-temporal integration processes (functional component (i); cf. main text Eqs. 1, 2 for details, recapitulated below),

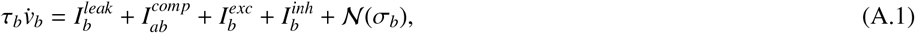

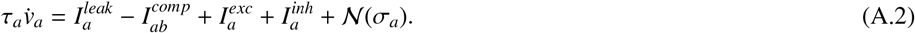

The exchange of information (functional component (ii)) between compartments is established by a conductance-based compartmental exchange equation (Eq. 8)

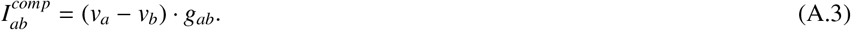

The model does not separate incoming projections by synapse receptor type. Instead, it captures whether they depolarize or hyperpolarize the apical and basal compartments, i.e., their excitatory or inhibitory impact (Eq. 5),

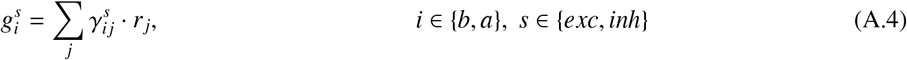

with constant synaptic efficacies 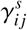 barring one exception, namely the voltage-dependent gating of excitatory apical

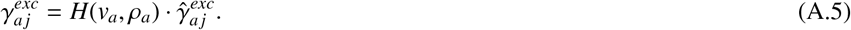

This apical gate is a single threshold nonlinearity, which is motivated by NMDA-receptor-mediated tuft spikes (Larkum et al., 2009) and represents an equivalent threshold of the dendritic calcium spike in BAC firing (Larkum et al., 1999). This activity-dependent gating of apical input establishes the basal-apical causality of the BAC mechanism (functional component (iii)). The apical compartment can only be depolarized above the gate threshold *ρ*_*a*_ *>* 0 by the terms 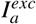 and 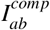. Here, 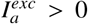 holds only if the compartment is already sufficiently depolarized, i.e., *H*(*v*_*a*_, *ρ*_*a*_) *>* 0 only if *v*_*a*_ *> ρ*_*a*_. Hence, apical integration depends on initial basal-to-apical influx via the other term 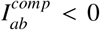. This necessity of basal-to-apical input corresponds to the role of the backpropagating sodium spikes of BAC. We assume this causal order for the regime of apical amplification, in which context modulates the response to basal input. In biological cells, strong tuft input alone can also evoke a dendritic calcium spike at a higher threshold and drive the cell without basal coincidence (Larkum et al., 1999; Larkum, 2013), which corresponds to the regime of apical drive (Graham et al., 2025).

The model assumes linearity in the compartmental exchange and rests directly on voltage levels of the compartments (Eq. A.3). This exchange mechanism is simpler than the pyramidal cell’s sodium and calcium spike generation and propagation, but suffices to capture the main computational aspects of BAC dynamics at the level of firing rates. Yet, the neural mass model could capture the spike-based nature of apical-basal voltage exchange more faithfully by a simple extension (Fig. A.1B). The variant replaces the term 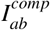, which enters both compartments with opposite signs (Eq. A.2), by two individual terms. They represent the proportions of cells in the mass model that currently propagate sodium (Na^+^) spikes to the apical compartment, 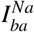, or calcium (Ca^2+^) spikes to the basal compartment,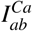,

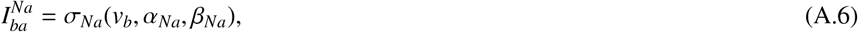

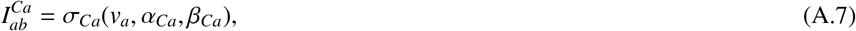

where *σ*_*i*_(·) represents the proportion of cells firing by a sigmoid function with parameters *α*_*i*_ and *β*_*i*_. Such a nonlinear, activation-function-based voltage-to-spike description is also employed in biologically more detailed spike-based neuron models of pyramidal cell firing (Fig. A.1A; Naud and Sprekeler, 2018; Payeur et al., 2021; Whyte et al., 2025). As a calcium spike that reaches the soma turns single action potentials into a burst, *σ*_*Ca*_ also gives the proportion of bursting cells. The variant could thus express the burst signals that burst-dependent codes read (Naud and Sprekeler, 2018; Payeur et al., 2021). It could also be related to the transient population bursts, i.e., brief peaks of population firing, that propagate between visual areas (Chen et al., 2022).

### Appendix B. Comparison with a multi-compartment model

#### B.1. Comparison with the multi-compartment model

This section complements Sect. 2.3.1 with the coding of the reference data, the simulation of the TPN, the fitting procedure, the fitted parameters (Tabs. B.1 and B.2) and the residuals of the fits (Fig. B.1).

##### Reference data

In the simulation code of Shai et al. (2015), the output rate of the multi-compartment model is the inverse of the shortest inter-spike interval. As in their analysis code, conditions without spikes (163 of the 651 conditions) are coded as −1 Hz and conditions with a single spike (30) as 0 Hz. The highest rate is 177 Hz.

##### Simulation of the TPN

The input-output map of the TPN (Fig. 5A,C) was computed with the parameters of the single TPN (Tab. A.1) on a grid of 77 × 77 inputs *r*_*FF*_, *r*_*FB*_ *∈* [0, 4]. Each simulation started from the resting state of Tab. A.1, i.e., the equilibrium without input, and ran without noise under constant inputs until no state variable changed by more than 10^−7^ within 100 ms. All inputs reached this equilibrium within 900 ms, and the equilibrium state was read out. For FB inputs between about 0.24 and 0.27, i.e., about 51 to 56 tuft synapses under the fitted map, the mutual inhibition of the VIP and SOM cells is bistable. There, the read-out is the state that is reached from rest, in which the SOM cells remain active. Between the grid points, the rate 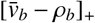 of Eq. 23 is interpolated linearly in both inputs. The time courses of Fig. 5B were simulated from the same resting state with an FF step at 25 ms and an FB step at 100 ms, with inputs that correspond to 250 basal and 150 tuft synapses under the fitted map (*r*_*FF*_ = 1.63, *r*_*FB*_ = 0.72). In addition to the deterministic solution, one simulation with additive Gaussian white noise on *v*_*b*_ and *v*_*a*_ shows the variability of the response, and its rate is smoothed with a 10-ms running mean.

##### Fitting procedure

All models were fitted to the 651 conditions by minimizing the sum of squared errors (Sect. 2.3.1). The composite, additive, multiplicative and Mod-E-I models were fitted by nonlinear least squares with a trust-region reflective algorithm (Branch et al., 1999), which respects the bounds of the Mod-E-I parameters. Because these fits can end in local minima, each fit was started from 500 points, and the solution with the lowest error was kept. For the composite model, the first starting point was the parameter set published by Shai et al. All other starting points were drawn at random over several orders of magnitude, each parameter as ±10^*u*^ with *u* uniform in [−1, 2.5]. The one-sigmoid forms of the additive and the multiplicative model that the released code of Shai et al. implements were fitted in the same way (Tab. B.1, bottom).

The TPN keeps its internal parameters, and only the four parameters of its map are fitted (Eq. 23). For given input scales *s*_*b*_ and *s*_*t*_, the prediction is linear in the offset *c*_0_ and the gain *c*_1_, which therefore follow in closed form from a linear regression of the data on the rate 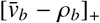. If the regression violates *c*_0_, *c*_1_ ≥ 0, the optimum lies on the boundary *c*_0_ = 0 or *c*_1_ = 0, where it again follows in closed form. The two input scales were found by evaluating the error on a regular grid of scale pairs and refining the best pair with the Nelder–Mead method (Nelder and Mead, 1965). The grid maps the largest synapse numbers of the data, 300 basal and 200 tuft synapses, onto model inputs between 10 % and 100 % of the simulated input range (181 values per input). The TPN fit thus needs no random starting points and is deterministic.

##### Fitted parameters

A sigmoid term can be written with either sign of its slope, since *c σ*(*x/s*) = *c* − *c σ*(−*x/s*). Table B.2 gives all parameters with positive slopes *a*_4_ and *b*_4_, into which the published parameters of the composite model were converted as well. Only the sum *a*_1_ + *b*_1_ of the additive model and the product of the two scale factors of the multiplicative model are determined by the data, so that *b*_1_ = 0 and *a*_2_ = 1 were set, respectively. In the fits of both models, the threshold *b*_3_ of the basal sigmoid lies far below the range of the data. The basal sigmoid then acts only through its saturating tail, in which *b*_1_, *b*_2_ and *b*_3_ trade off against each other and are not individually meaningful. **Mod-E-I model**. Equation 22 is the steady state of the Mod-E-I unit (Eq. E.1) for constant input. Its excitation combines the basal and the apical conductance with the modulation function 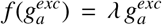 (Eq. E.3), with conductances 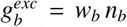 and 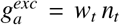 that are proportional to the numbers of active synapses. Its leak current is 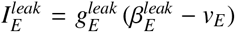, and its inhibitory current 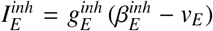 is driven by the basal input, 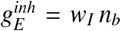, as for an inhibitory unit (Eq. E.4) whose steady state follows its FF drive linearly. Setting 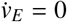 and applying the linear rate function *m*_*E*_ (*v*_*E*_ − *ρ*_*E*_) gives Eq. 22 with

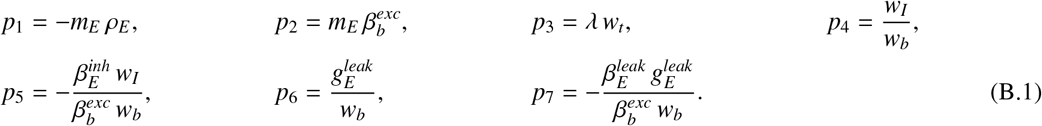

The bounds *p*_2_, … , *p*_7_ ≥ 0 hold for 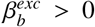 and reversal potentials 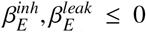. The seven fitted parameters determine seven combinations of the ten parameters of the unit. For 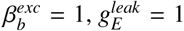 and the same weight *w*_*t*_ = *w*_*b*_ of tuft and basal input, the fit corresponds to the unit parameters of Tab. B.2. Its inhibitory conductance hardly enters the denominator of Eq. 22 (*w*_*I*_ ≈ 10^−3^ *w*_*b*_), whereas its very negative reversal potential lets it subtract a current proportional to the basal input in the numerator. The fitted inhibition is therefore subtractive rather than divisive (shunting). The fitted rate is not rectified, so the model can predict negative rates. The fit thus describes a Mod-E-I unit with a linear rather than a threshold-linear rate function (cf. Eq. 10).

##### TPN map

The best TPN fit maps 300 basal synapses onto *r*_*FF*_ = 1.96 and 200 tuft synapses onto *r*_*FB*_ = 0.95, both inside the search range (Tab. B.2). Two checks show that neither the constraint on the offset nor the exact input scales determine the result (Tab. B.1, bottom). First, without the constraint *c*_0_ ≥ 0, the offset becomes slightly negative with the same *R*^2^. Second, *R*^2^ exceeds 0.80 over a range of FF input scales around the optimum, whereas it falls off faster with the FB input scale (Fig. B.1B).

#### Parameter search

The internal parameters of the TPN were set during model development in an iterative exploration, carried out manually and with the help of AI coding agents. In this exploration, candidate parameter sets and circuit variants were scored against the data of Shai et al. with the procedure above. This exploration did not sample the parameter space systematically and is therefore not exhaustive. Its last step screened how single parameters affect the fit and then refined four of them. These were the tonic input to the SOM cells, the efficacy of the FB input onto the apical compartment, the SOM inhibition of the VIP cells and the excitatory reversal potential of the basal compartment. A coordinate descent followed by Nelder–Mead searches over these four parameters maximized the *R*^2^ of the map fit at equilibrium and thus found a local optimum. The best values were rounded to two decimal places, which changed 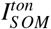 from 0.3 to 0.55, 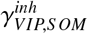 from 1 to 0.23 and 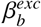 from 1 to 1.13 (Tab. A.1). The search lowered the efficacy of the FB input onto the apical compartment to 0 75. This is equivalent to 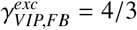 for an FB input that is rescaled by 0.75, a factor that the input scale *s*_*t*_ of the map absorbs. Earlier development versions of the TPN, which differed in parameter values and in details of the interneuron circuit, reached *R*^2^ between 0.71 and 0.85.

**Table B.1:** Fits to the input-output map of the multi-compartment model (651 conditions). *k* is the number of fitted parameters, *R*^2^ the coefficient of determination (Sect. 2.3.1) and RMSE the root-mean-square error. The upper part lists the models of Fig. 5A and the lower part further fits and checks of the composite, the additive, the multiplicative and the TPN model.

| Model | Eq. | $k$ | $R^2$ | RMSE (Hz) |
| --- | --- | --- | --- | --- |
| Composite | 19 | 8 | 0.901 | 21.1 |
| Additive | 20 | 7 | 0.801 | 29.9 |
| Multiplicative | 21 | 7 | 0.788 | 30.9 |
| Mod-E-I | 22 | 7 | 0.779 | 31.6 |
| TPN | 23 | 4* | 0.886 | 22.7 |
| **Checks** |  |  |  |  |
| Composite, published parameters |  | 8 | 0.878 | 23.5 |
| Additive, one-sigmoid form of the released code |  | 5 | 0.788 | 30.9 |
| Multiplicative, one-sigmoid form of the released code |  | 5 | 0.802 | 29.9 |
| TPN, free offset ($c_0 = -0.7$ Hz) | | 4 | 0.886 | 22.7 |

\*Only the four parameters of the map are counted, here and in the checks. The internal parameters of the TPN were tuned during model development, in part by a non-exhaustive search against the same data (Appendix B.1), and are not counted in $k$ .

**Table B.2:**
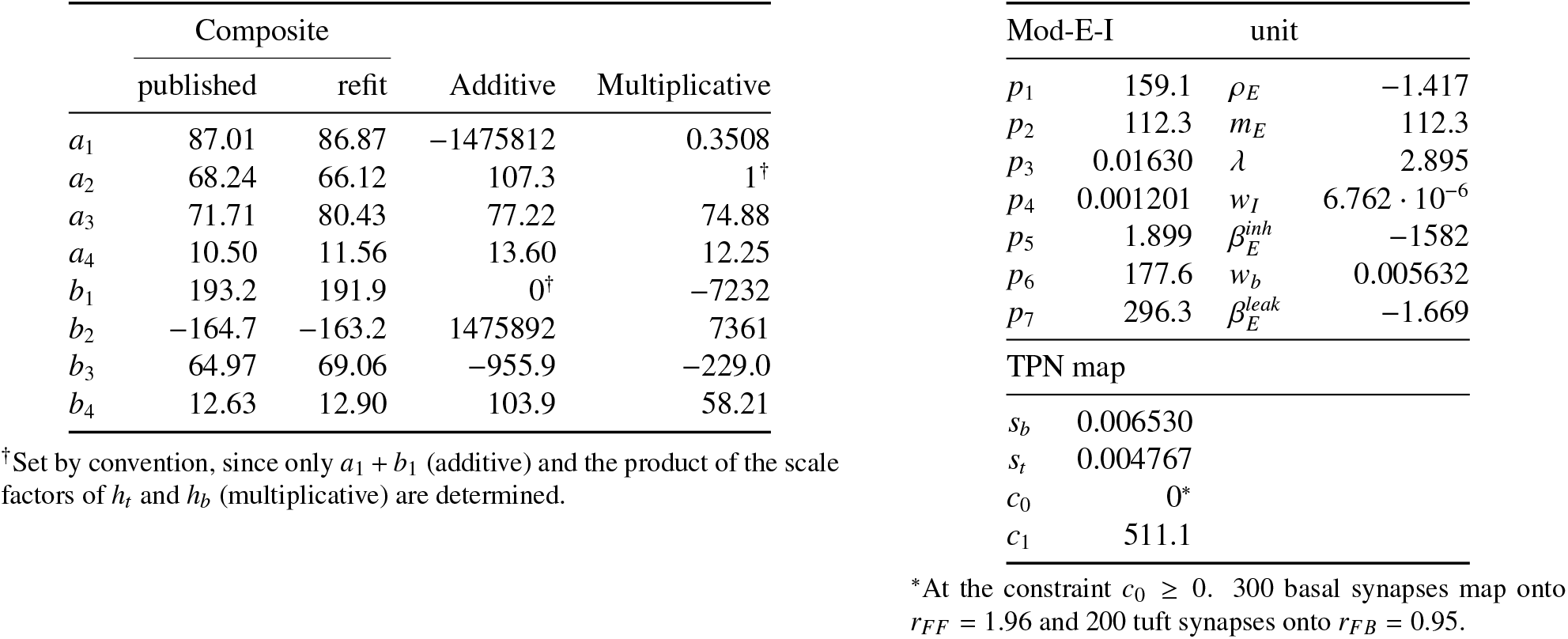
Fitted parameters. The left part gives the models of Shai et al. (Eqs. 19–21), written with positive slopes *a*_4_, *b*_4_. The composite model is given with the parameters published by Shai et al. (2015) and refitted here. The right part gives the Mod-E-I model (Eq. 22) and the map of the TPN (Eq. 23). Next to each *p*_*i*_, it lists the parameter of the Mod-E-I unit that follows from it (Eq. B.1) for 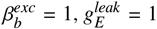 and *w*_*t*_ = *w*_*b*_, which the data do not determine. The thresholds *a*_3_, *b*_3_ and slopes *a*_4_, *b*_4_, as well as *b*_1_, *b*_2_ of the composite model and *p*_6_, *p*_7_, are given in synapses. The parameters *a*_1_, *a*_2_ of the composite and additive models, *b*_1_, *b*_2_ of the additive and multiplicative models, *p*_1_, *p*_2_, *c*_0_ and *c*_1_ are given in Hz, and *p*_3_, *s*_*b*_ and *s*_*t*_ per synapse, while all other parameters are dimensionless. Of the unit parameters, *m*_*E*_ is given in Hz, and the weights *w*_*b*_ and *w*_*I*_ of the basal input relative to 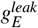 per unit of *n*_*b*_, while all others are dimensionless.

**Figure B.1:**
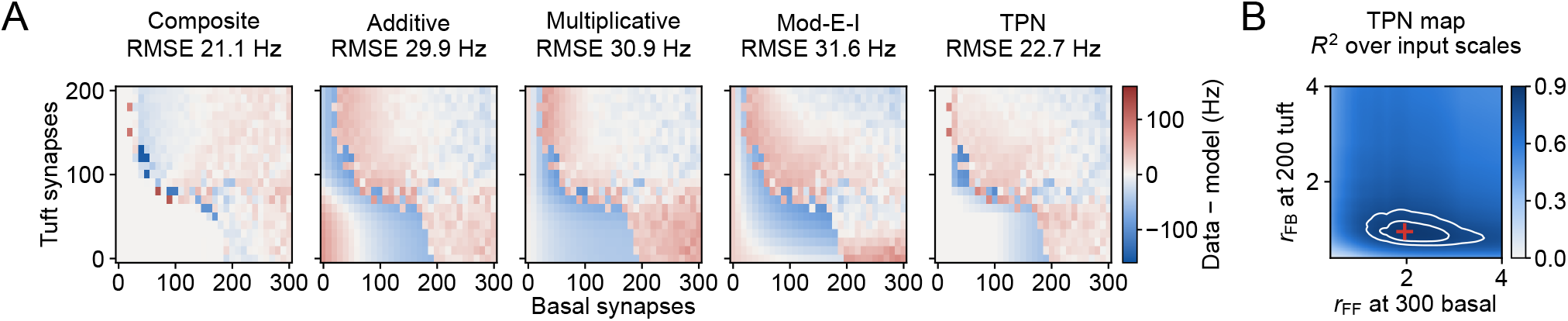
Residuals of the fits and *R*^2^ of the TPN map. (A) Residuals 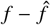 of the five fits of Fig. 5A as a function of the numbers of active basal (x-axis) and tuft (y-axis) synapses. Red marks where the model lies below the data and blue where it lies above them, and the panel titles give the root-mean-square error. (B) *R*^2^ of the TPN fit as a function of its two input scales, expressed as the FF input *r*_*FF*_ reached by 300 basal synapses (x-axis) and the FB input *r*_*FB*_ reached by 200 tuft synapses (y-axis). For each pair of scales, offset and gain are optimal. White contours mark *R*^2^ = 0.80 and 0.85, and the red cross marks the best fit (Tab. B.2).

### Appendix C. Learning in a pool of two-point neurons

#### C.1. Coincidence tag of the single TPN

The coincidence tag (Eqs. 14 and 15) marks the TPNs whose basal synapses may change. Figure C.1 shows where in the input space of the single TPN a tag forms. The tag was computed together with the input-output map of Fig. 5C, with the tag parameters of the flanker task, and it does not act back on the other state variables of the TPN. A tag forms only where FF input coincides with an open apical gate (Fig. C.1A). Within this region, the coincidence term *c*_*ab*_ grows with both inputs, whereas the tag saturates close to its maximum *k*_2_*/k*_3_ = 1. Tonic input to the VIP cells shifts the FB input that is needed for a tag (Fig. C.1B). Inhibitory tonic input raises this FB threshold, and excitatory tonic input lowers it, because the VIP cells dis-inhibit the apical compartment through the SOM cells. For tonic inputs of −1 and below, no tag forms within the input region of the data.

**Figure C.1:**
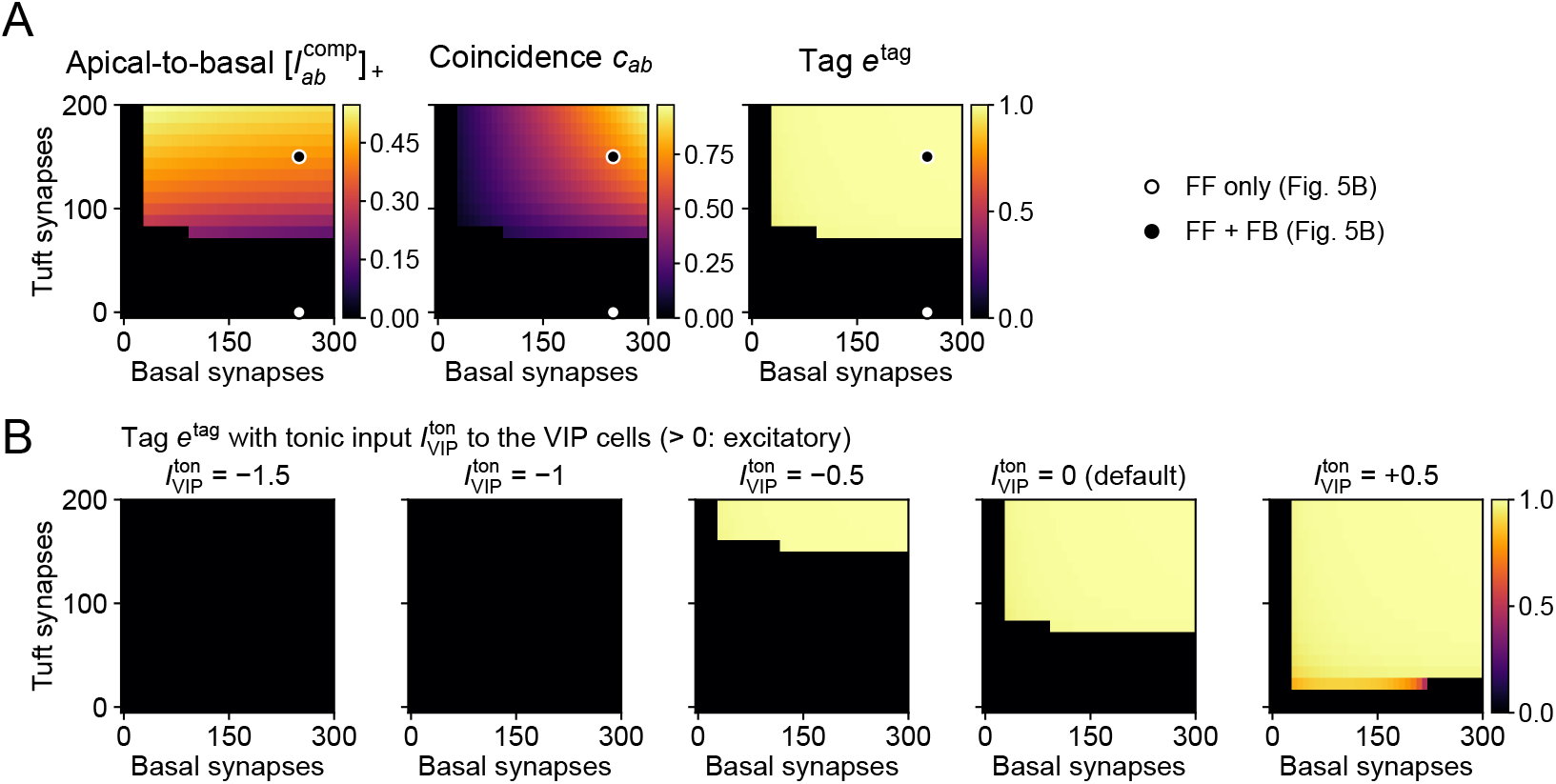
Coincidence tag of the single TPN over FF and FB input. (A) Rectified exchange current 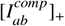 (Eq. 8), coincidence term *c*_*ab*_ (Eq. 15) and tag *e*^*tag*^ (Eq. 14) at equilibrium under constant FF input *r*_*FF*_ and FB input *r*_*FB*_, starting from the resting state. The input region is that of Fig. 5A,C, with the inputs given as numbers of basal and tuft synapses under the fitted map (Tab. B.2). Open and filled dots mark the two input conditions of Fig. 5B. (B) Tag for tonic input 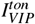 to the VIP cells between −1.5 and +0.5 (positive values excitatory, 0 is the default of the single TPN in Tab. A.1), computed on the same grid of inputs.

#### C.2. Flanker learning task

This section gives the protocol and the evaluation of the learning simulations of Sect. 4.2 (Figs. 3 and 6). The TPNs have the parameters of the column Pool of Tab. A.1, and the parameters of the learning rules and of the protocol are listed in Tab. C.1.

##### Network

The pool consists of two TPNs, each with its own PV, SOM and VIP cells. The two TPNs interact only through their PV cells, which receive the pooled rates of both TPNs, whereas the SOM and VIP cells act within the circuit of their own TPN. The recurrent and interneuron drives, the decision and the teacher use the rates [*v*_*b,i*_ − *ρ*_*b*_]_+_ of Eq. 10 (Sect. 2.3.2). The basal learning rule reads the rectified basal potential [*v*_*b,i*_]_+_ instead, and the apical learning rule reads the rectified apical potential [*v*_*a,i*_]_+_ (Sect. 2.2.2). Every VIP cell of the pool receives the tonic input 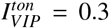 Without it, a context whose initial FB input onto its TPN is weak does not open the apical gate during training, so that its apical efficacies decay with *k*_4_ and the context is not learned. In 10 networks trained without the tonic input, this happened for 3 of the 20 pairings of a context with its TPN. With 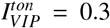, every context is learned, while the firing rates of the circuit at rest stay the same, because the VIP cells rest at *v*_*VIP*_ = 0.17, below their output threshold *ρ*_*q*_ = 0.2.

##### Inputs

The stimulus *y* is an image of 5 × 5 pixels, in which *y*^*A*^ is a horizontal and *y*^*B*^ a vertical bar through the center, with the value 1 on the bar and 0 elsewhere (Fig. 3). The context *c* consists of two flanker patches of 5 × 5 units each. The context *c*^*A*^ carries a horizontal bar in both patches, collinear with *y*^*A*^, and *c*^*B*^ a vertical bar. The uninformative context *c*^?^ is the superposition of the two, and the silenced context is zero. Every context is normalized to unit sum, so that the contexts differ in which units are active, not in their total input. The stimulus drives the basal compartment and the PV cell of TPN *i* with the FF input 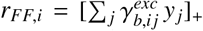. The context drives the apical compartment and the VIP cell with the FB input 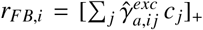. Both sets of efficacies are learned, and their initial values are drawn independently from *U*[0, 1) for each of the 30 networks.

##### Training

Each network is trained on 320 trials of the four unambiguous pairings in random order, 75 % of them congruent. These are 120 trials each of *y*^*A*^ with *c*^*A*^ and of *y*^*B*^ with *c*^*B*^, and 40 trials each of the incongruent pairings of *y*^*A*^ with *c*^*B*^ and of *y*^*B*^ with *c*^*A*^. Ambiguous stimuli are never shown during training. A training trial starts from the resting state and presents stimulus and context for its whole duration *T*_*train*_. Eligibilities and tags start every trial at zero, and only the efficacies are carried over to the next trial. Before the dopamine onset *t*_*DA*_, no dopamine is released (*D*^*\**^ = 0), so that the basal efficacies do not change (Sect. 2.2.2). From *t*_*DA*_ on, the teacher compares the output rates of the two TPNs at every time step. It sets *D*^*\**^ = +1 if the TPN assigned to the class of the stimulus has the higher value and *D*^*\**^ = −1 otherwise, and exact ties are broken by a seeded coin. Because *c*_*ab*_ ≥ 0, the tag stays between 0 and *k*_2_*/k*_3_ = 1 (Eq. 14). The eligibility traces decay with time constants of 100 ms, whereas biological eligibility traces bridge delays of seconds between activity and reward (Gerstner et al., 2018). Traces of this length are not needed in the task, because dopamine is released while stimulus and context are still present, so that coincidence and reward overlap within every trial.

**Table C.1:** Protocol and parameters of the learning rules of the flanker task (Sect. 2.2.2). The tag parameters also hold for the single TPN of Fig. C.1.

| Quantity | Symbol | Value |
| --- | --- | --- |
| **Protocol** |  |  |
| Networks, training trials per network |  | 30, 320 |
| Fraction of congruent training pairings |  | 0.75 |
| Duration of a training trial | $T_{train}$ | 200 ms |
| Dopamine onset | $t_{DA}$ | 20 ms |
| Duration of a test trial | $T_{test}$ | 200 ms |
| **Basal learning** |  |  |
| Efficacy time constant | $\tau_b^\gamma$ | 200 ms |
| Eligibility time constant, learning rate | $\tau_{e_b}^{syn}, \eta_b$ | 100 ms, 1 |
| **Coincidence tag** |  |  |
| Time constant | $\tau^{tag}$ | 1 ms |
| Decay, rise, saturation | $k_1, k_2, k_3$ | 0.01, 4, 4 |
| **Dopamine** |  |  |
| Dopamine time constant | $\tau_D$ | 1 ms |
| **Apical learning** |  |  |
| Efficacy time constant | $\tau_a^\gamma$ | 200 ms |
| Eligibility time constant, learning rate | $\tau_{e_a}^{syn}, \eta_a$ | 100 ms, 1 |
| Efficacy decay | $k_4$ | 0.008 |
| **Testing** |  |  |
| Test trials per condition: accuracy, all other measures | $n$ | 10, 40 |
| Decision threshold | $\vartheta$ | 0.05 |
| Bisection steps for $\alpha_{seed}^*$ | | 40 |

##### Testing

Test trials run without dopamine and with frozen efficacies. Stimulus and context are presented for the duration *T*_*test*_, and the decision is read from the margin *m* (Sect. 2.3.2) at the end of the presentation. Each test trial starts from a state in which each of the five potentials of each TPN circuit is offset from the resting state by an independent draw from U(−0.5, 0.5). All conditions are tested with the same *n* initial states. In *P*(*A*), trials with *m* = 0 count half. All measures were computed from the efficacies at the end of training.

##### Accuracy

A decision counts as correct only if the margin exceeds the decision threshold *ϑ* in the direction of the stimulus class, i.e., *m > ϑ* for *y*^*A*^ and *m <* −*ϑ* for *y*^*B*^. The accuracy was measured before training and after every 10 training trials, and the accuracy after training is the last of these values. A network discriminates the stimuli if it decides correctly in more than half of the test trials with *y*^*A*^ and in more than half of those with *y*^*B*^.

##### Point of subjective equivalence

The point of subjective equivalence 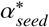 of a network is located in test trials with silenced context that start from the resting state. Bisection halves the interval [0, 1] 40 times around the sign change of the margin, which determines 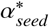 to better than 10^−12^. By construction, *P*(*A*) for *y*^?^ with silenced context is close to 0.5, which serves as a control.

##### Neurometric curves

To average the neurometric curves across networks, each curve was expressed as a function of the distance 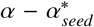 to the point of subjective equivalence of its network. The curves were then interpolated linearly onto a common grid with a step of 0.05 and averaged over the networks with data at each distance. Averaging along *α* itself would blur the effect of context, because the steep part of the curve lies at a different *α* for each network.

##### Context effect and apical selectivity

The context effect Δ (Sect. 2.3.2) is 0 if context has no influence on the decision and 1 if context alone determines it. The accuracy with incongruent context is the accuracy on the unambiguous stimuli paired with the context of the other class, with 40 test trials per condition. The apical selectivity of a TPN (Fig. C.2E,F) is the mean apical efficacy 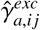 from the context units *j* that are active only in *c*^*A*^ minus the mean from those that are active only in *c*^*B*^. It is positive for a TPN that has associated *c*^*A*^ with its own activity.

##### Ablations

In the ablations of Fig. 6A, the tag 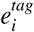, the dopamine level *D* or both were replaced by the constant 1 in the basal rule (Eq. 12). The code that a network has learned is read from its efficacies, independently of the class labels. The code is consistent if each TPN prefers one stimulus by its basal drive and the collinear context by its apical input, and if the two TPNs prefer different classes. A consistent code is correct if TPN 1 prefers class A, and mirrored if TPN 1 prefers class B. For mirrored networks, the accuracy and the context effect are also given with the class labels of the two TPNs swapped (hatched in Figs. 6A and C.7B).

##### Sensitivity analyses

In the tonic-VIP analysis (Fig. 6E), the constant current 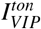 to the VIP cells was set during training and testing to the values −2, −1.2, −0.9, −0.6, −0.45, −0.3, −0.2625, −0.225, −0.1875, −0.15, −0.1125, −0.075, −0.0375, 0, 0.15, 0.3, 0.45, 0.6, 0.9, 1.2 and 2, i.e., more densely where the results change most. From 0.6 on, the SOM cells are already silent at rest. From −0.45 downward, the VIP cells stay below their output threshold for all FB inputs of the task. Further values in either direction therefore change neither training nor test results. Every condition, including the ablations, was run with the same 30 networks, i.e., with identical initial efficacies, and otherwise unchanged parameters.

#### C.3. Additional results on four-factor learning

Figures C.2–C.7 show the learning simulations of Sect. 4.2 in more detail, with the protocol and the measures of Appendix C.2. All 30 networks discriminate the stimuli from training trial 50 on (Fig. C.2). Meanwhile, the apical selectivities of the two TPNs grow in opposite directions. In the training trials shown, both TPNs form a tag once their apical gates open, and the sign of the dopamine signal then determines whether the basal efficacies of both TPNs grow or shrink, more strongly for the TPN whose tag forms first (Figs. C.3 and C.4). Figure C.6 details the sensitivity to tonic VIP input of Fig. 6E. Below a tonic VIP input of about 0.15, the context effect decreases. Below about 0, the accuracy and the accuracy with incongruent context decrease as well. From −0.45 downward, all three measures stay at the level of untrained networks. From 0.6 upward, the SOM cells are silent at rest, and one of the 30 networks does not learn to discriminate the stimuli. Both accuracies and the context effect therefore stay at 0.97. Because of the strong VIP dis-inhibition, the two TPNs of this network form tags alike and receive the same updates. Their basal efficacies become identical within the first 50 training trials, so that almost all test trials remain undecided (|*m*| ≤ 0.05). For tonic inputs of 0 and above, a point of subjective equivalence exists in all 30 of the 30 networks. For lower inputs, it exists in 23 networks at −0.15, in 17 at −0.3, and in 13 from −0.45 downward. Figure C.7 details the ablations of Fig. 6A. Without the tag, the two TPNs approach the same basal pattern, and 28 of the 30 networks end without a consistent code. Without dopamine, half of the networks discriminate the stimuli within 20 training trials, but 14 of the 29 consistent codes are mirrored. Without both factors, 28 networks end without a consistent code.

**Figure C.2:**
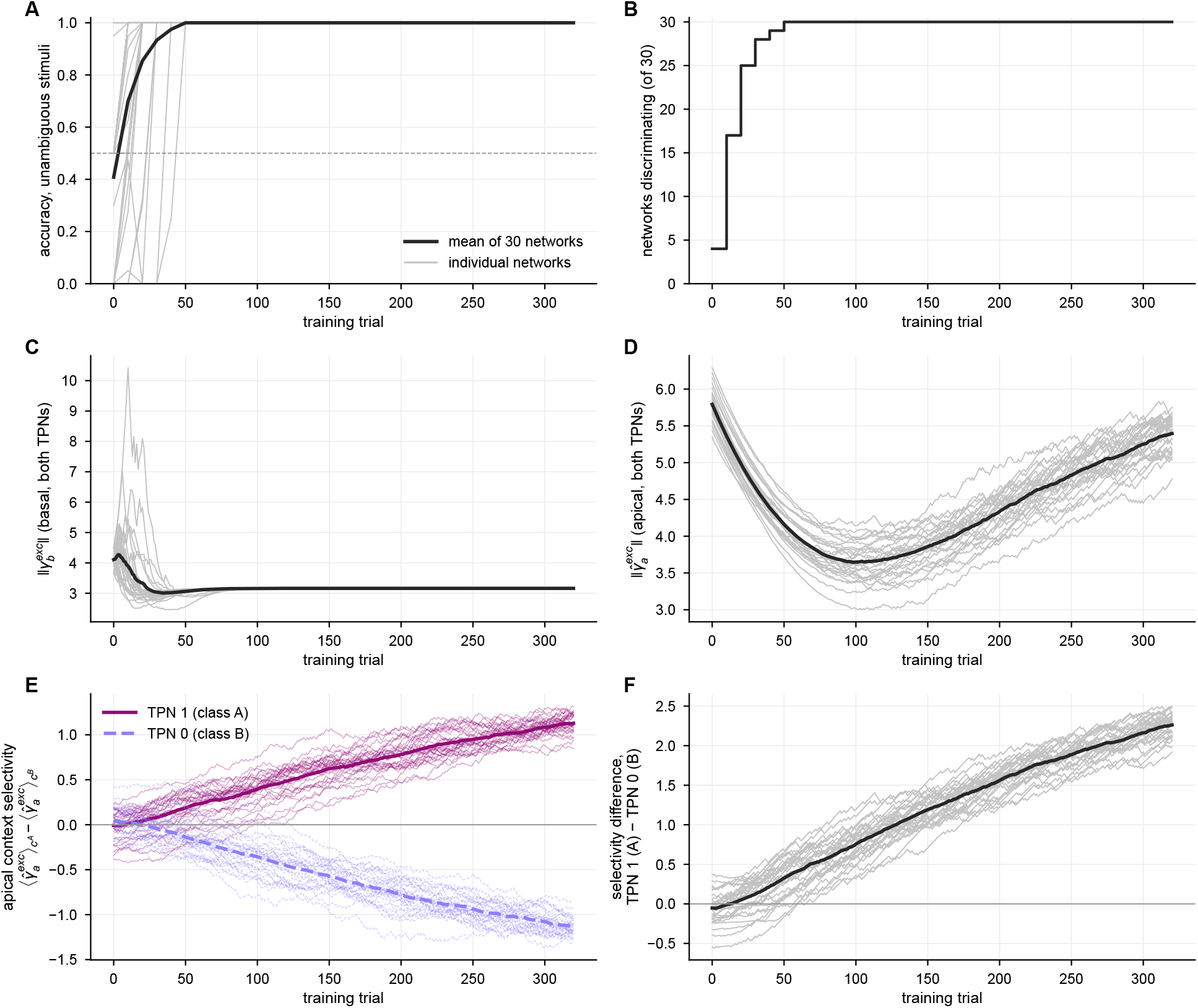
Learning curves of the flanker task. Over the training trials, the panels show the accuracy on the unambiguous stimuli (A), the number of networks that discriminate the stimuli (B) and the norms of the basal and the apical efficacies of both TPNs (C,D). E shows the apical selectivity of TPN 1 (class A, magenta, solid) and TPN 0 (class B, lavender, dashed), and F the difference between the two. In A and C–F, thin lines show the 30 training runs and thick lines their mean, and the dashed line in A marks an accuracy of 0.5.

**Figure C.3:**
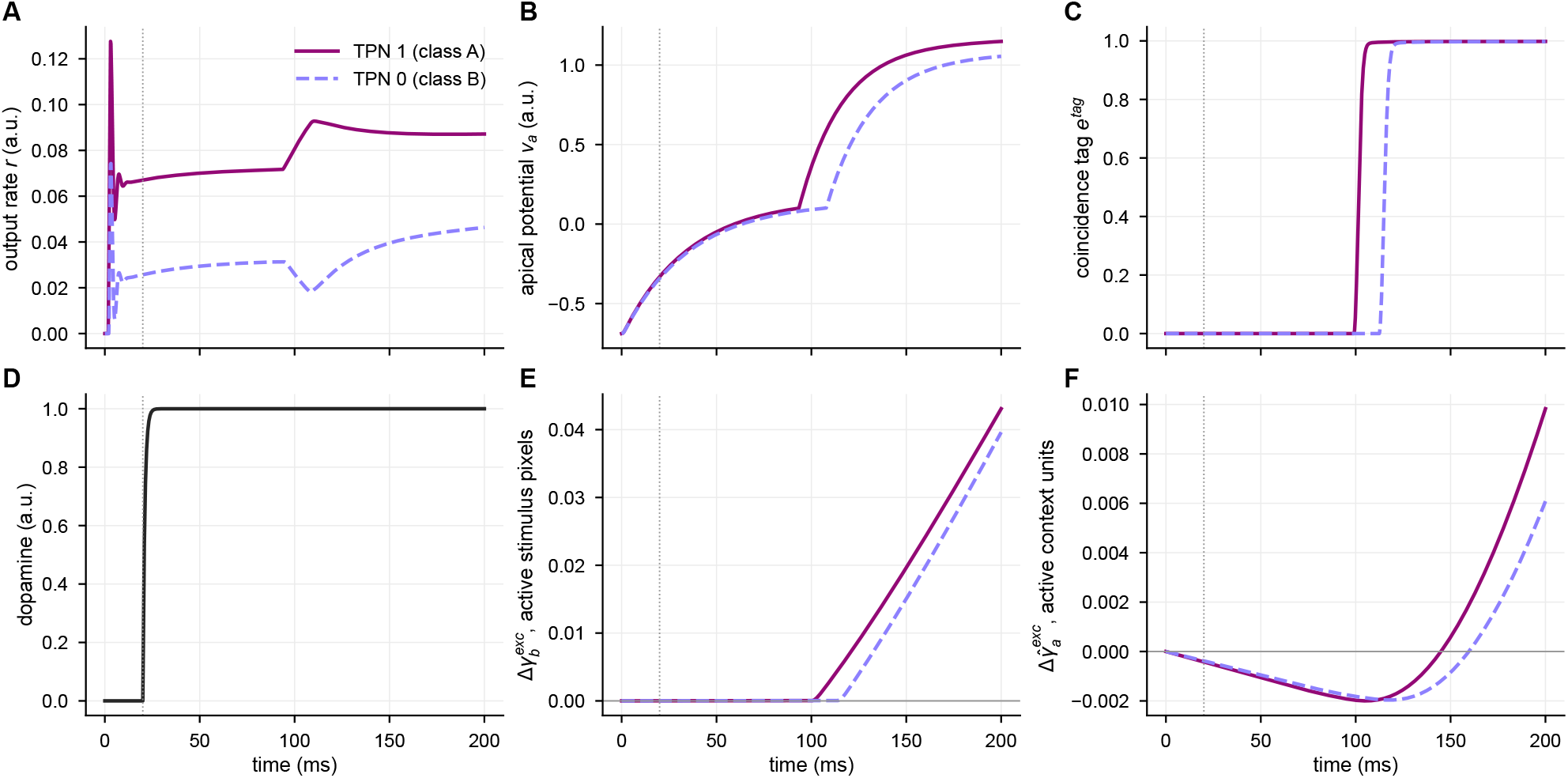
Rewarded training trial. In the first rewarded trial of the example training run of Fig. 6B, the horizontal bar *y*^*A*^ is paired with the incongruent context *c*^*B*^. The panels show the output rates *r* that the decision function reads (A), the apical potentials *v*_*a*_ (B) and the coincidence tags *e*^*tag*^ (C) of TPN 1 (class A, magenta, solid) and TPN 0 (class B, lavender, dashed). Panel D shows the dopamine level *D*, and E and F show the mean change since trial onset of the basal efficacies from the active stimulus pixels and of the apical efficacies from the active context units, respectively. Dotted vertical lines mark the dopamine onset *t*_*DA*_.

**Figure C.4:**
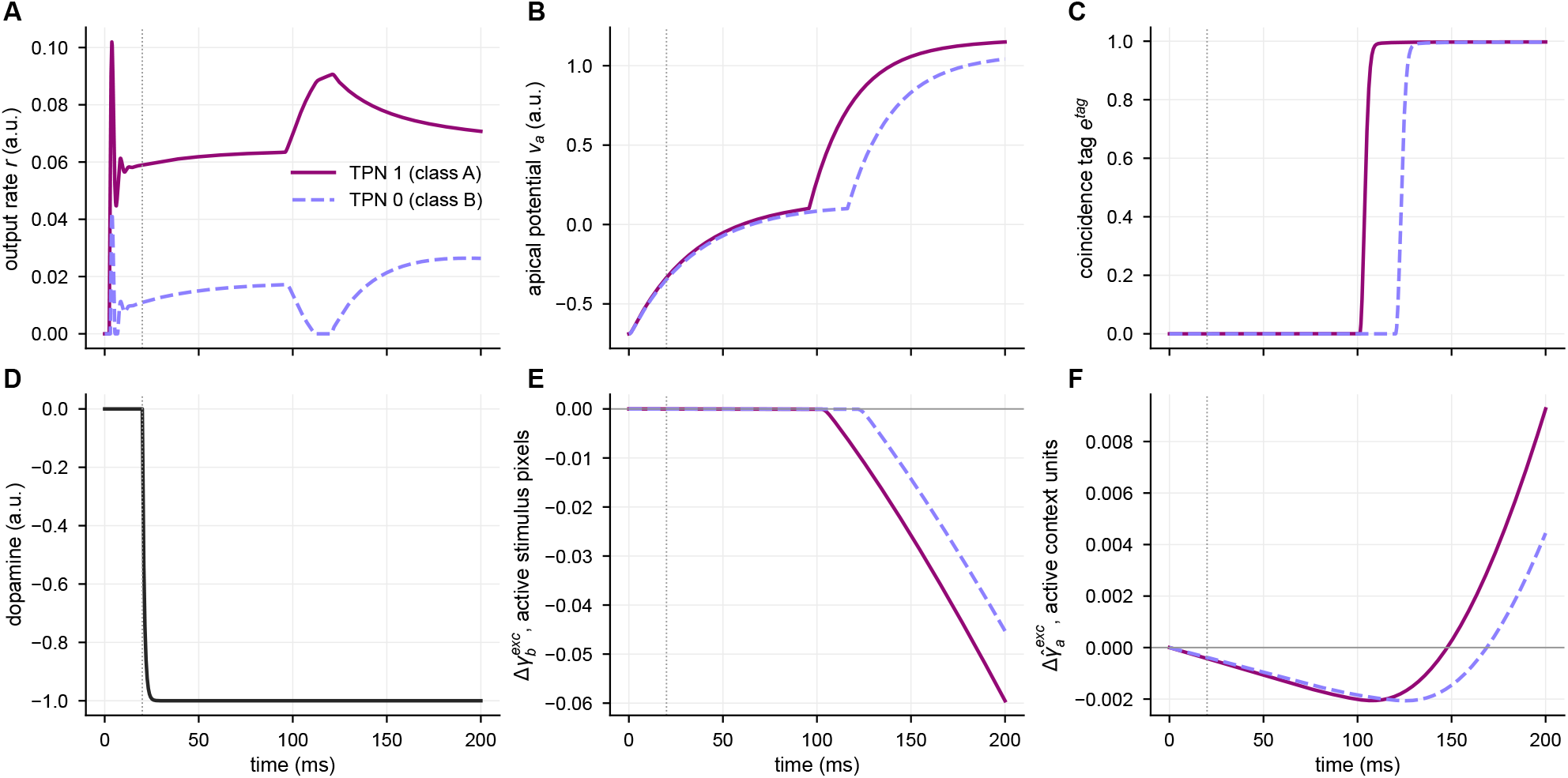
Punished training trial. In the first punished trial of the same training run, the vertical bar *y*^*B*^ is paired with the congruent context *c*^*B*^, with panels as in Fig. C.3. TPN 1 has the higher output rate, the dopamine signal is negative, and the basal efficacies of both TPNs decrease, those of TPN 1 more strongly (E).

**Figure C.5:**
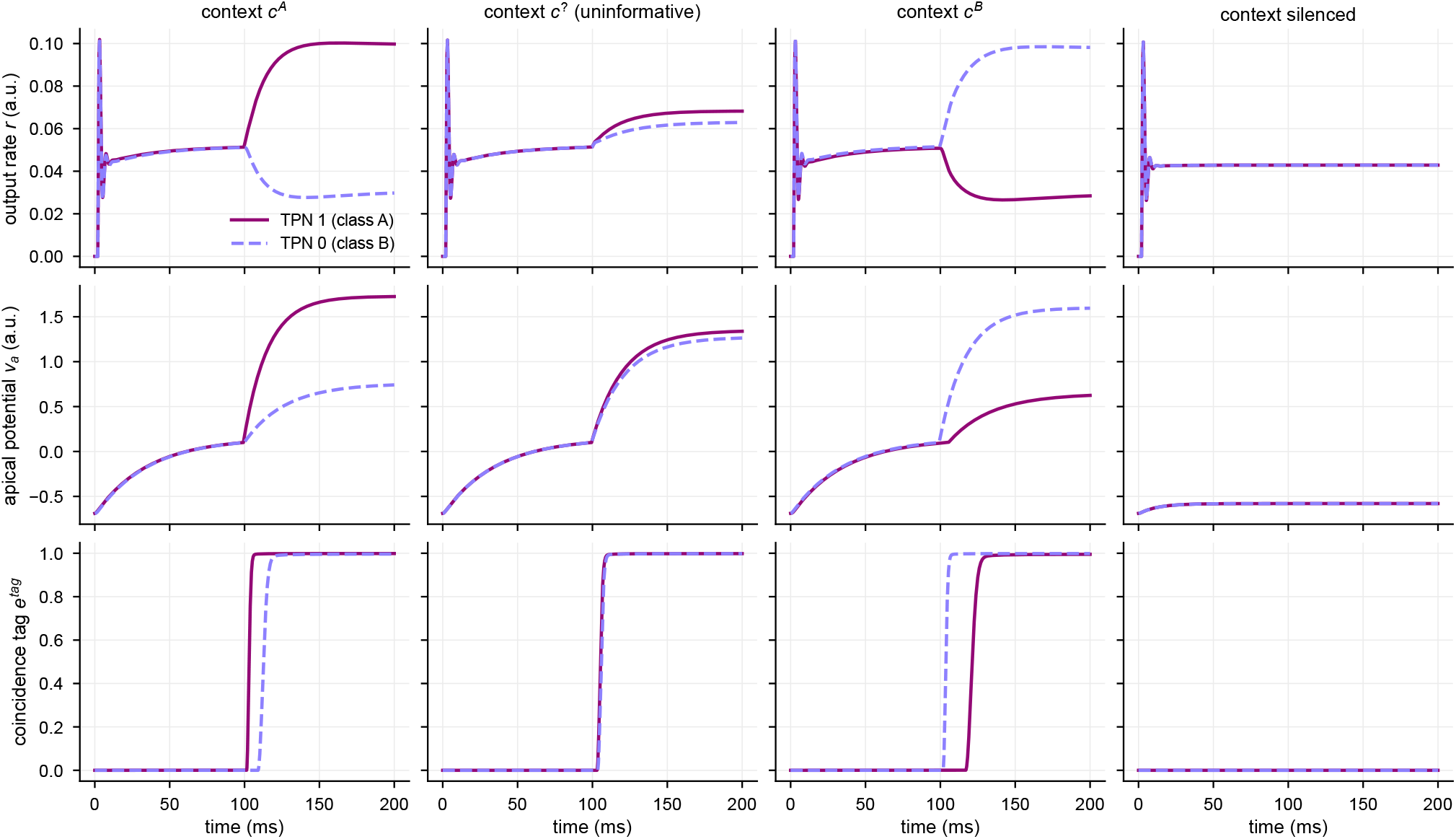
Test trials after training. The trained network of the example training run receives its ambiguous stimulus *y*^?^ with the contexts *c*^*A*^, *c*^?^, *c*^*B*^ or silenced context (columns from left to right), starting from the resting state without offsets. The rows show *r, v*_*a*_ and *e*^*tag*^ of both TPNs as in Fig. C.3A–C.

**Figure C.6:**
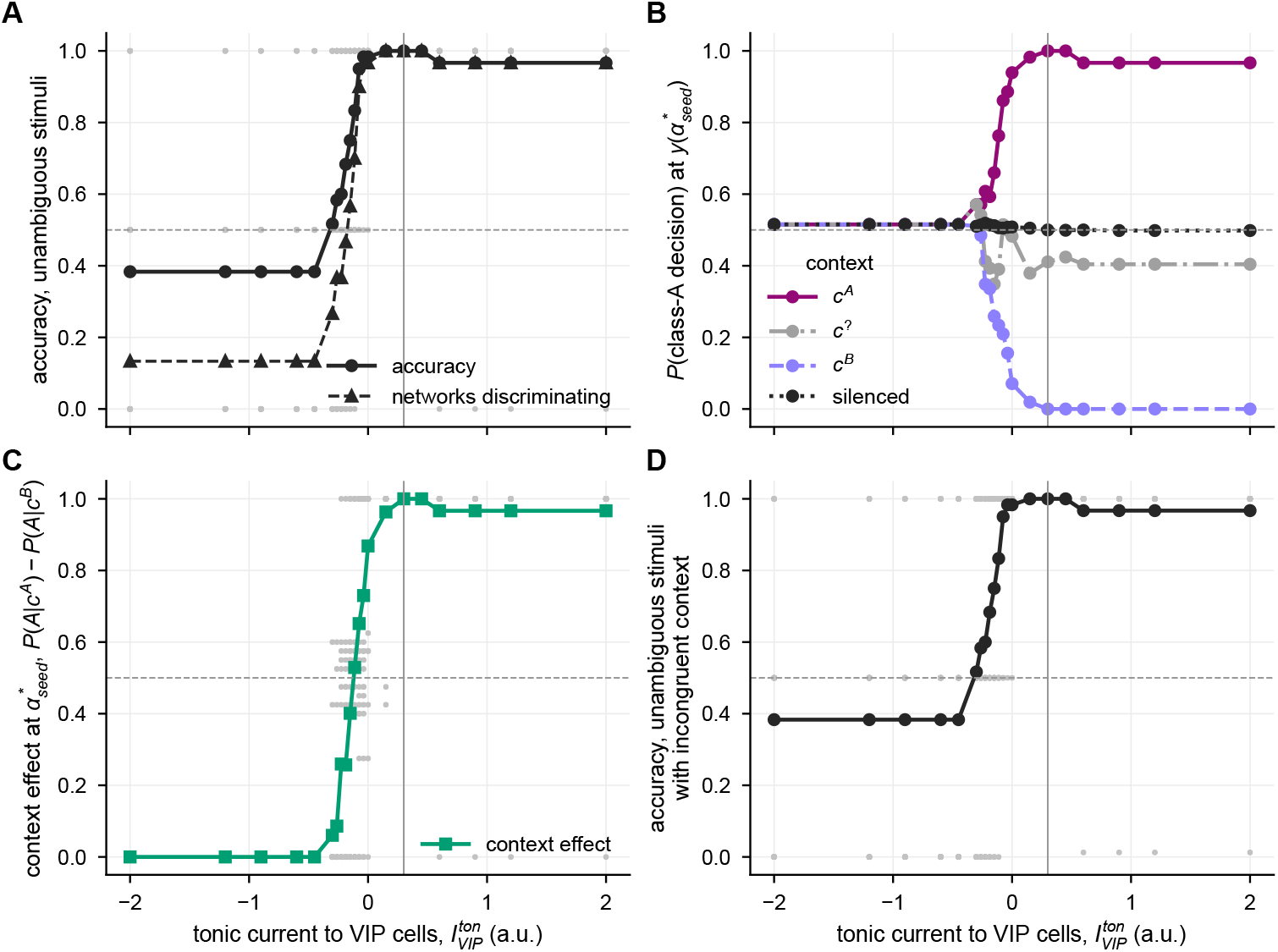
Sensitivity of learning to tonic VIP input. Over the tonic input 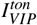 to the VIP cells (default 0.3, vertical line), the panels show the accuracy on the unambiguous stimuli and the fraction of networks that discriminate them (A) as well as the accuracy with incongruent context (D). B shows the fraction *P*(*A*) of decisions for class A on the ambiguous stimulus *y*^?^ with context *c*^*A*^ (magenta), *c*^?^ (gray), *c*^*B*^ (lavender, dashed) or silenced context (black, dotted). C shows the context effect Δ. Lines connect the means over the 30 training runs, in B and C over those with a point of subjective equivalence, and gray dots show individual runs.

**Figure C.7:**
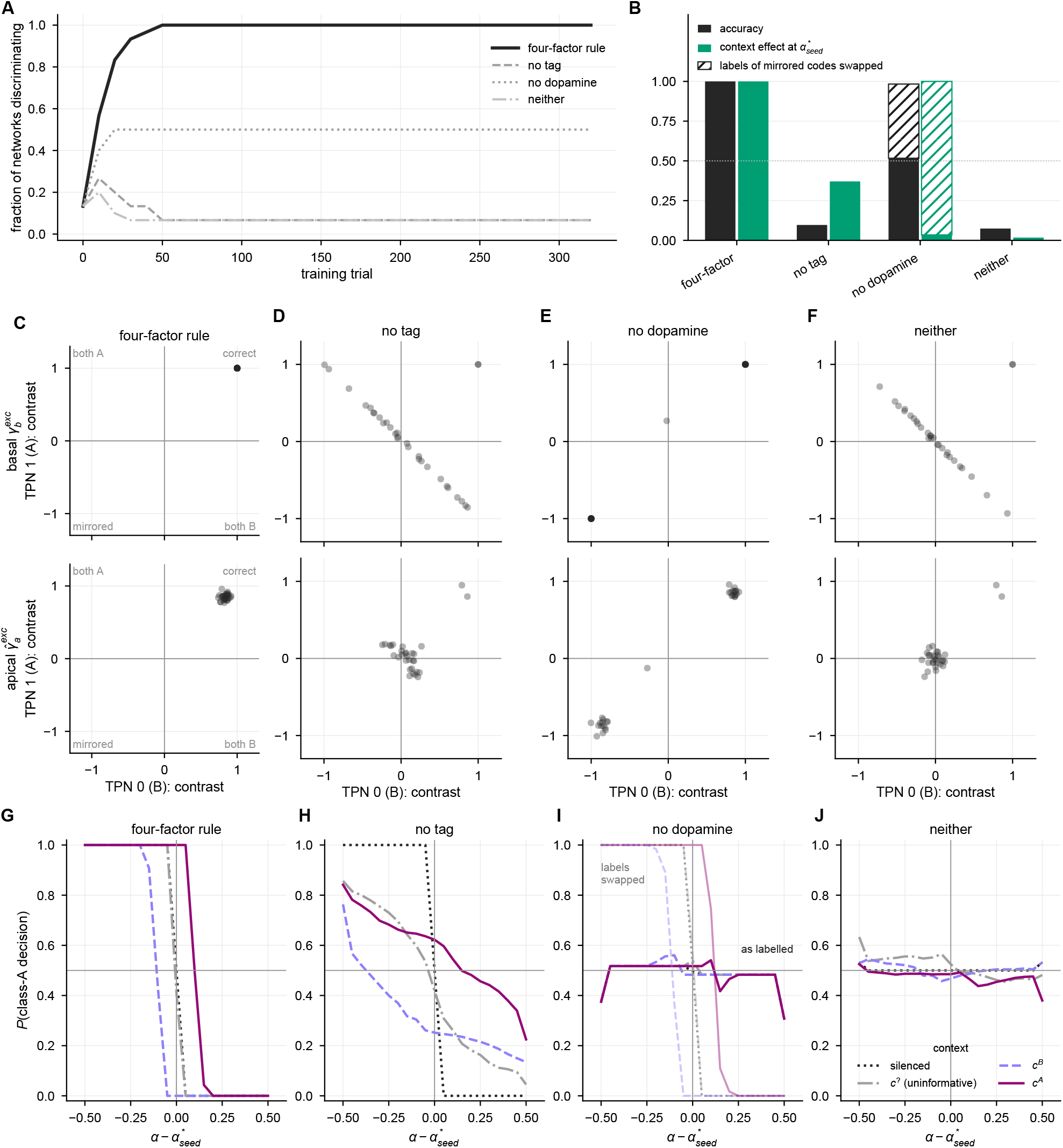
Ablations of the basal learning rule. The four-factor rule (Eq. 12) is compared with the same rule without the coincidence tag (no tag), without the dopamine signal (no dopamine) and without both (neither), each with the same 30 networks. (A) Fraction of networks that discriminate the stimuli over the training trials. (B) Accuracy on the unambiguous stimuli (black) and context effect Δ (green) after training, with the accuracy and the context effect of mirrored codes with swapped labels hatched. (C–F) Code of each network after training for the four rules (Appendix C.2). Each dot is one network, placed by the template contrasts of TPN 0 (horizontal) and TPN 1 (vertical) for the basal (top) and the apical efficacies (bottom). The template contrast of a TPN is the correlation of its efficacies with the stimulus or context pattern of its own class minus that with the pattern of the other class. Correct codes lie near (1, 1), mirrored codes near (−1, −1), and codes in which both TPNs prefer the same class in the other two quadrants. (G–J) Neurometric curves averaged as in Fig. 6C for the four rules. Without dopamine (I), light lines show the curves with the class labels of mirrored codes swapped.

### Appendix D. Networks of two-point neurons

#### D.1. Network simulations

This section complements Sect. 3.3 with the inputs and the initial states of the network simulations of Figs. 7 and 8. The parameters and the connectivity of the TPNs are listed in Tab. A.1 (column Network), and the numerical methods in Tab. A.2.

##### Inputs

The FF inputs and the external FB input are Gaussian profiles over the preferred feature values *θ*_*n*_ with additive noise,

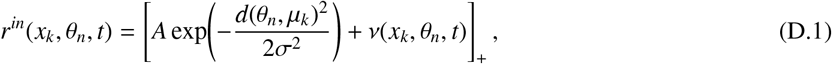

with peak *A*, center *µ*_*k*_ and width *σ*, where *d*(*θ*_*n*_, *µ*_*k*_) is the shortest distance on the ring. The noise *ν* is drawn independently for every TPN from a Gaussian distribution with standard deviation *σ*_*ν*_ and is held constant within time bins (Sect. 3.3.1). Because the noise can make the sum negative where the profile is small, the input is rectified. Table D.1 lists the three types of input. The bars of Fig. 8 receive no external FB input.

**Table D.1:** Inputs of the network simulations (Eq. D.1). In the line of rings (Fig. 7), configurations 1, 2 and 4 receive the uncertain and configurations 3 and 5 the certain FF input, and configurations 4 and 5 receive the bias in addition from 0.5 s on (Sect. 5.1). Configuration 6 equals configuration 4 with the bias at the opposite side of the ring. In the bars (Fig. 8), the two end rings receive the certain and the interior rings the uncertain FF input.

| Input | Target | Peak $A$ | Width $\sigma$ | Noise $\sigma_\nu$ | Center $\mu_k$ |
| --- | --- | --- | --- | --- | --- |
| Uncertain FF input | basal compartment, PV cells | 1 | $0.35\pi$ | 0.05 | line: $1.00\pi, 1.05\pi, 0.97\pi, 1.03\pi, 0.95\pi$ (rings 0 to 4); bars: $\pi$ |
| Certain FF input | basal compartment, PV cells | 4 | $0.05\pi$ | 0.05 | line: as uncertain input; bars: $0.75\pi$ |
| Bias (external FB) | apical compartment, VIP cells of ring 0 | 4 | $0.05\pi$ | 0.03 | $0.85\pi$ (configuration 6: 0) |

##### Initial state and random numbers

All variables start from the resting state of the TPN (Tab. A.1), except the basal potentials, to which independent draws from U(0, 0.1) are added. The initial states and the input noise are drawn from seeded generators and are the same in all configurations, so that the configurations differ only in their inputs and connections.

#### D.2. Context opposite to uncertain FF input

Configuration 6 tests whether biasing context can steer a TPN population toward a feature value for which the population receives almost no FF evidence (Fig. D.1). It equals configuration 4 of Fig. 7, except that the bias excites ring 0 at the direction 0, where the uncertain FF input is less than 2 % of its peak (Tab. D.1). None of the rings changes its response. The bias depolarizes the apical compartment of the TPN at the bias direction from −0.69 to only about −0.05, below the gate threshold *ρ*_*a*_, because this TPN lacks the basal drive that would depolarize its apical compartment through the exchange current. Context that mismatches uncertain FF input thus leaves the TPN population unaffected.

**Figure D.1:**
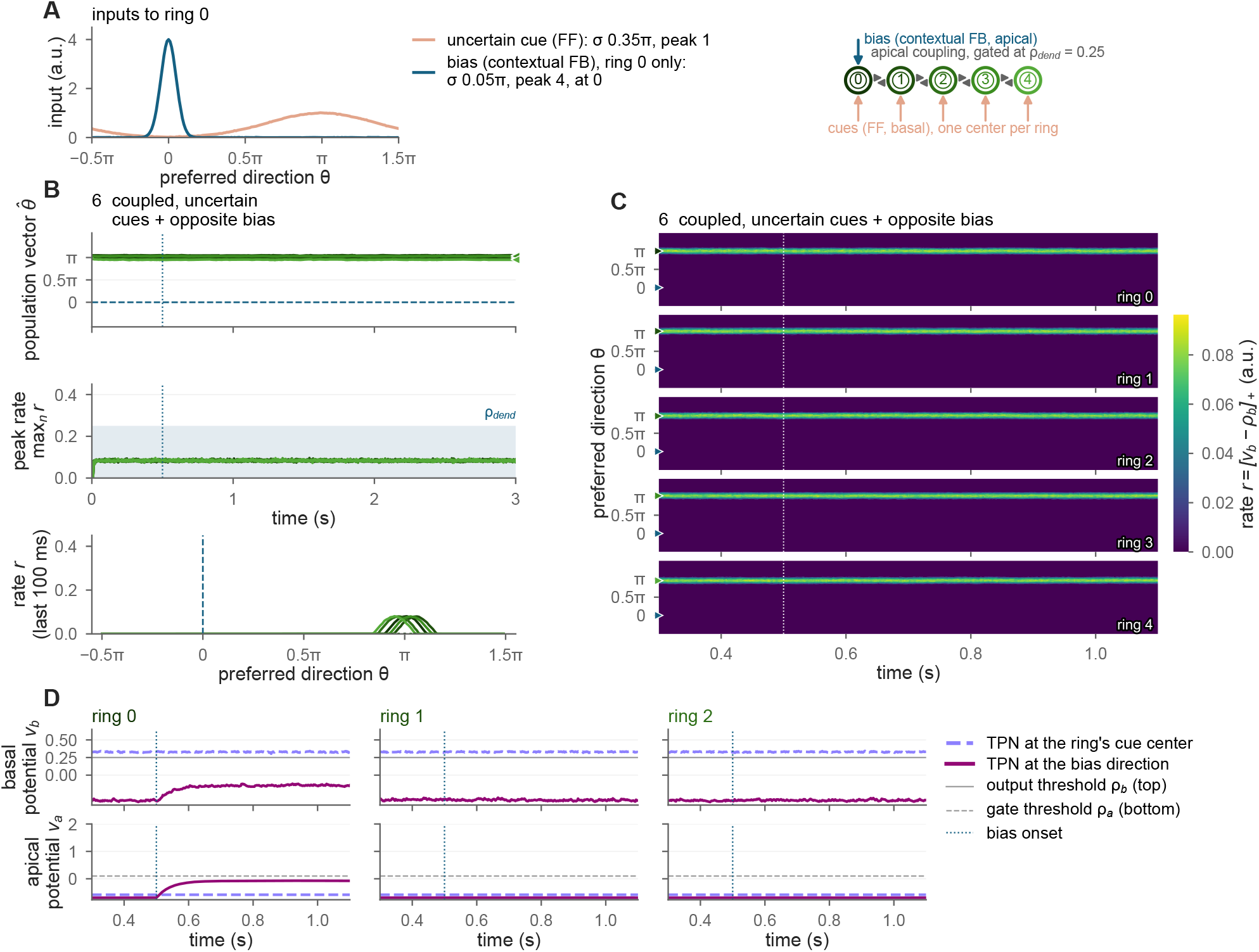
Context opposite to uncertain FF input in a line of TPN rings. Panels as in Fig. 7A–D for configuration 6, in which ring 0 receives the uncertain FF cue and, from *t* = 0.5 s on, the bias at the direction 0 (A). The bottom row of B shows the rate profiles at the end of the simulation, and D shows *v*_*b*_ and *v*_*a*_ of the TPN closest to the ring’s cue center and of the TPN closest to the bias direction in rings 0 to 2.

#### D.3. Propagation over bars of 500 and 1000 rings

Bars of 500 and 1000 rings test whether the evidence propagation of Fig. 8 keeps its speed over longer distances (Fig. D.2). They were simulated with the protocol of the shorter bars (Sect. 3.3.3) for 64 s and 115 s, which suffices for every interior ring to disambiguate its estimate. The input noise was held constant within bins of 5 ms, and its sequence repeated every 7 s, because a table of the noisy inputs for the whole run (Sect. 3.3.1) would have exceeded the available memory. The two fronts that start at the ends of a bar advance by one ring about every 192 ms, and the bars align after 47.9 s and 95.8 s. The time to full alignment thus grows by about 96 ms per ring of bar length over bars of 10 to 1000 rings, close to the 102 ms of the bars of up to 100 rings (Fig. 8G).

**Figure D.2:**
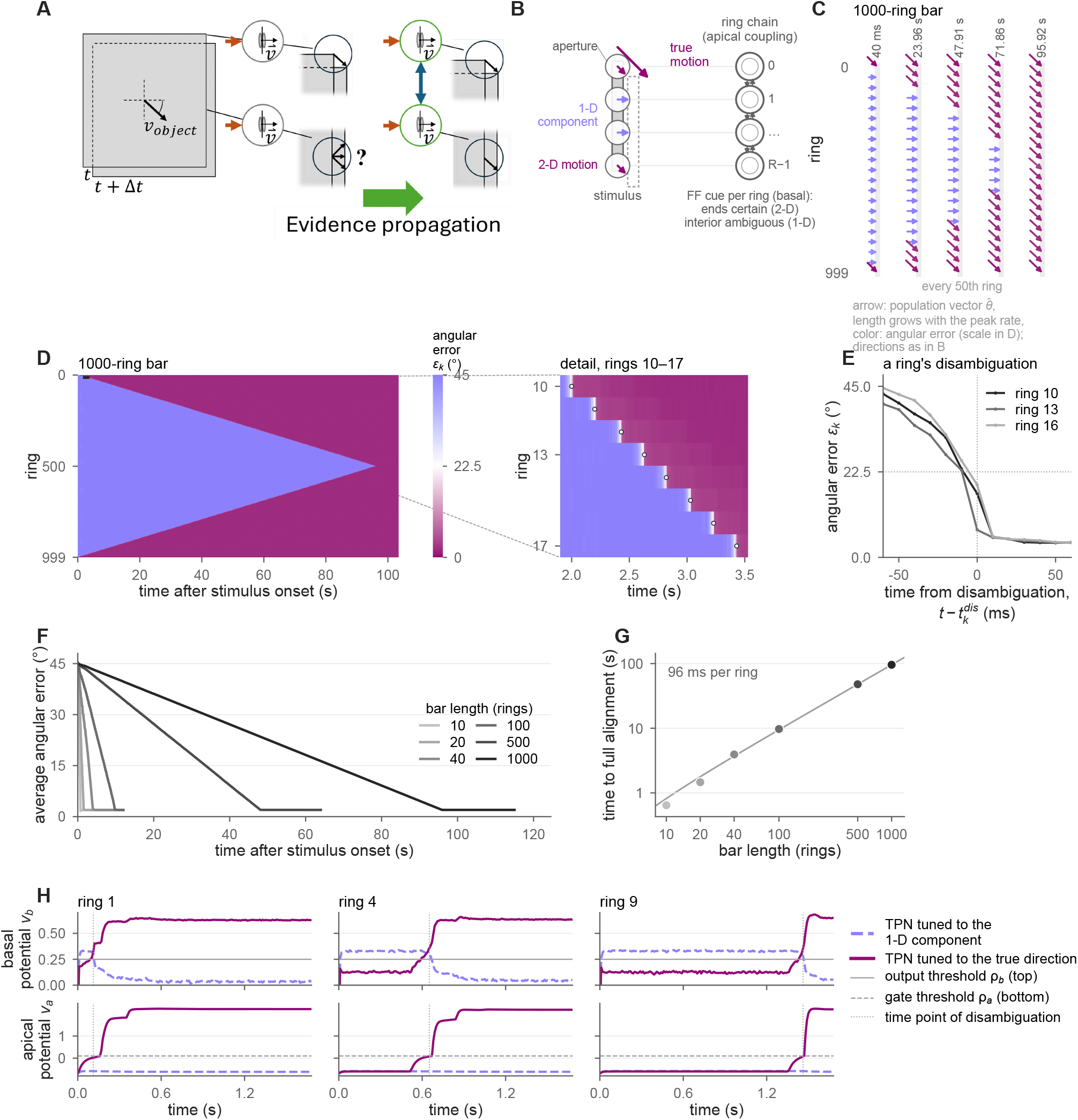
Visual motion integration over bars of 500 and 1000 TPN rings. Panels as in Fig. 8 for the TPN network. C to E show a bar of 1000 rings, of which C shows every 50th ring. F and G include the bars of 500 and 1000 rings together with those of 10 to 100 rings, with the intact network only in F and logarithmic axes in G.

**Figure D.3:**
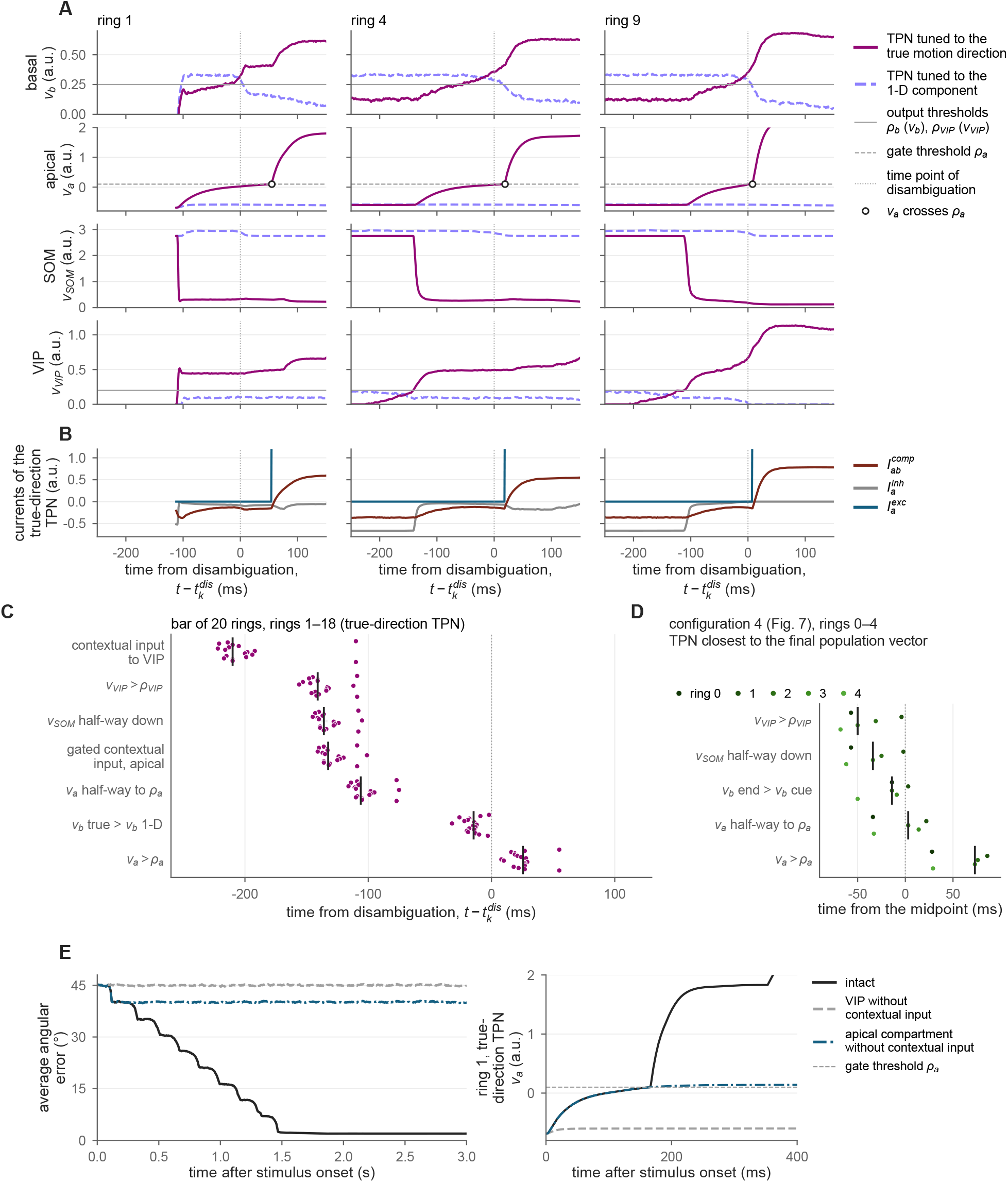
Order of events during disambiguation in a network of TPN rings. (A) Potentials of the TPN tuned to the true motion direction (magenta, solid), of the TPN tuned to the 1-D motion component (lavender, dashed) and of their SOM and VIP cells in rings 1, 4 and 9 of the bar of 20 rings of Fig. 8. They were recorded every 1 ms and are aligned to each ring’s time point of disambiguation 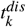 (Eq. 29, dotted lines). From top to bottom, the rows show *v*_*b*_, *v*_*a*_, *v*_*S OM*_ and *v*_*VIP*_. Gray lines denote the output thresholds *ρ*_*b*_ and *ρ*_*VIP*_ (Eq. 10) and the apical gate threshold *ρ*_*a*_ (dashed), and circles mark the time point at which *v*_*a*_ of the TPN tuned to the true motion direction crosses *ρ*_*a*_. (B) Currents of the TPN tuned to the true motion direction in the same rings, namely the exchange current 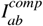 (Eq. 8), the inhibition 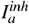 of its apical compartment by its SOM cell and the gated apical excitation 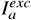. The gated apical excitation is zero while the gate is closed and leaves the axis once it opens. (C) Times of events at the TPN tuned to the true motion direction relative to 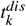 in the 18 interior rings of the bar (dots, one per ring, with black ticks at the medians). Each event is the first time point from which a condition holds until the end of the recording. From top to bottom, the rows show the events in the order of their median times, namely the arrival of contextual input at the VIP cell, the onset of VIP output (*v*_*VIP*_ *> ρ*_*VIP*_), the drop of *v*_*S OM*_ half-way between its levels before the front arrives and 200 ms after 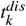, the arrival of gated contextual input at the apical compartment, the rise of *v*_*a*_ half-way to *ρ*_*a*_, the time point at which *v*_*b*_ of the TPN tuned to the true motion direction exceeds *v*_*b*_ of the TPN tuned to the 1-D motion component, and the crossing of *ρ*_*a*_. For SOM, gray bars span the medians obtained with levels at 25 % and 75 % of the way instead, and the order of the events does not change with this choice. (D) Times of the same events as in C, except the arrivals of contextual input, at the TPN closest to the final population vector of each ring in configuration 4 of Fig. 7, with the ring colors of that figure. The arrivals of contextual input are not shown, as ring 0 receives the bias directly and ring 3 receives contextual input from its neighbors from stimulus onset on. The times are given relative to the time point at which the population vector passes the midpoint between the ring’s cue center and the bias direction. Here the rows are again ordered by their median times, and the basal row marks the time point at which *v*_*b*_ of this TPN exceeds *v*_*b*_ of the TPN closest to the ring’s cue center. (E) Average angular error over the interior rings of the bar (Eq. 28, left) and *v*_*a*_ of the TPN tuned to the true motion direction in ring 1 (right). The intact network (black) is compared with two control networks, in which either the VIP cells receive no contextual input from the neighboring rings (gray, dashed) or the apical compartments receive no contextual input while the VIP cells keep theirs (blue, dash-dotted). Potentials and currents are given in model units (a.u.). See Appendix D.4 for further details.

#### D.4. Order of events during disambiguation

To resolve the order of events by which a ring disambiguates its estimate, the bar of 20 rings of Fig. 8 and configuration 4 of Fig. 7 were simulated again with the same inputs and initial states. The potentials and input currents of every TPN and of its PV, SOM and VIP cells were recorded every 1 ms (Fig. D.3). In rings 2 to 17 of the bar, the contextual input from the neighboring ring reaches the VIP cell of the TPN tuned to the true motion direction about 210 ms before the ring disambiguates its estimate. At first, the neighboring TPN fires below the dendritic threshold *ρ*_*dend*_ (Eq. 25), so that only the VIP cell receives this input. The VIP cell starts to fire 112 to 156 ms before disambiguation, while the gated contextual input to the apical compartment is still zero. The potential of the SOM cell of the same TPN drops half-way 4 to 16 ms later. The VIP cell of the TPN tuned to the 1-D motion component, in contrast, stays below its output threshold. Without SOM inhibition, the exchange current from the more depolarized basal compartment raises the apical potential from about −0.60 toward *ρ*_*a*_, while the gated apical excitation remains zero. As *v*_*a*_ approaches *v*_*b*_, the exchange current drains less from the basal compartment. This is the main reason why *v*_*b*_ of this TPN exceeds *v*_*b*_ of the TPN tuned to the 1-D motion component 9 to 32 ms before disambiguation. The apical gate opens 8 to 29 ms after disambiguation, and within 200 ms of disambiguation *v*_*b*_ of the winning TPN rises from 0.35 to 0.63. The VIP cell of the next ring starts to fire within 10 ms of this gate opening. In configuration 4, the VIP cell of the winning TPN likewise starts to fire and the potential of its SOM cell drops half-way before the population vector passes the midpoint between the ring’s cue center and the bias direction. Its apical gate opens 28 to 86 ms after this midpoint.

In a control network without contextual input to the VIP cells, no interior ring disambiguates its estimate within 12 s, although the rings next to the bar’s ends still receive gated contextual input in their apical compartments. In a second control network, the apical compartments receive no contextual input, while the VIP cells keep theirs. The two rings next to the bar’s ends then disambiguate their estimates as early as in the intact network. Their winning TPNs, however, fire at a rate of only 0.16 instead of 0.38, which is too weak to switch on the VIP cells of the next rings, and the front stops there. The contextual input thus disambiguates a ring’s estimate through the dis-inhibition of the apical compartment via VIP and SOM cells. The subsequent opening of the apical gate amplifies the winning TPN enough to signal a state of coherence and to pass the evidence on to the next ring.

### Appendix E. Reduced two-point neuron models

#### E.1. Abstraction levels of the two-point neuron model

**Table E.1:** Comparison of the TPN model and its reduction to Mod-E-I units (cf. Fig. 2) with respect to the questions they can address.

|  | TPN | Mod-E-I |
| --- | --- | --- |
| State variables per node | five (Eqs. 1–3) | two (Eqs. E.1, E.4) |
| Fit to the input-output map of the multi-compartment L5Pyr model | two input-dependent regions; 88.6 % | no separate regions; 77.9 % |
| Time course of amplification | resolved (Fig. 5B) | instantaneous* |
| Context mismatching uncertain drive | no effect on the population mode (Fig. D.1) | steers the population mode (Fig. E.2) |
| Control of operating regime | interneuron drive, $g_{ab}, \rho_a$ | choice or gating of $f$ |
| Coincidence signal for learning | apical-to-basal current | algebraic proxy $g_b^{exc} \cdot f(g_a^{exc})$ |
| Cell-type-specific predictions | PV, SOM, VIP | single inhibitory population |
| Large-scale steady-state networks | costly | efficient |

\*The reduction yields a Mod-E-I unit with two ODEs, one for its excitatory and one for its inhibitory cell (Eqs. E.1, E.4). The Mod-E-I model fitted in Fig. 5A is the steady state of this unit (Eq. 22). This steady state needs no numerical integration over time and responds instantaneously to the inputs (Appendix E.2). The rings of Appendix E.3 integrate only $v_E$ over time and take the inhibition from the instantaneous rates. There, too, the modulation $f(g_a^{exc})$ acts on the basal drive without a time course of its own. Other variants integrate the FB signal over time before it amplifies the driving input, in a dis-inhibitory interneuron (Brosch and Neumann, 2014b) or in the apical compartment of the simplified TPN (Fig. 2B; Schmid and Neumann, 2025).

The proposed TPN model describes processing motifs of layer V pyramidal cells and their adjacent interneuron circuitry to explain feedforward-feedback (FF-FB) integration mechanisms from the data of a multi-compartment model (Sect. 4). Here, we show how the TPN model can be structurally simplified to arrive at modulated excitatory-inhibitory (Mod-E-I) models (Brosch and Neumann, 2014a). The two model variants share some properties while differing in others (Tab. E.1). The simplification results in a dimension reduction of the TPN system from five to two dimensions. Consequently, the Mod-E-I models have less explanatory power in fitting the input-output map of the multi-compartment L5Pyr model (see results in Fig. 5A). Yet, they conserve the main processing motifs of TPNs and reduce the computational cost of simulating a network in large-scale investigations. Here, we reduce the TPN model to a simplified version of two structural components, subsuming the apical compartment and its adjacent local inhibitory interneuron circuitry into algebraic terms.

#### Simplification of the apical compartment

The proposed TPN model possesses two sites of integration, the basal and the apical compartment. Explicitly modeling these two sites of integration comes with two implications, i.e., having separate integration streams and modeling bidirectional coupling dynamics.

##### Separate integration streams

Different input streams (FF vs. FB) are treated by separate basal and apical integration functions (Eqs. 1, 2). This separation allows for different streams to have different impacts on the model pyramidal neuron’s computation in terms of asymmetric FF-FB integration (Figs. 1B, 5A, TPN model).

##### Bidirectional coupling

The existence of separate basal and apical compartments allows for modeling bidirectional interaction dynamics between these basal and apical integration functions (Eq. 8). This bidirectional coupling together with the voltage-dependent apical gate (Eq. 6) allows for modeling of basal-apical integration dynamics, such as those of backpropagation-activated calcium spike firing at the level of firing rates (BAC; Figs. 1B, 5B). Such internal cell communication accounts for, e.g., the temporal coincidence detection of feedforward sensory features and top-down contextual field signals (Larkum, 2013).

##### Simplification

Mod-E-I models capture the asymmetric FF-FB integration of separate integration streams without modeling the dynamical apical compartment in its bidirectional coupling with the basal cell compartment (Fig. 5A, Mod-E-I model). Instead, a modulation transfer function (MTF) is described that performs the separate stream integration at the apical tuft instantaneously. As a result, Eqs. 1, 2, 6, 8 simplify to

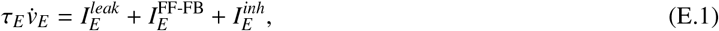

where the term

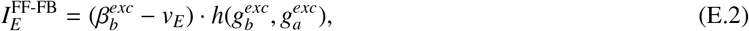

combines basal FF and apical FB conductances according to a function

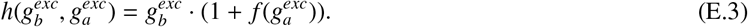

The function *h*(·, ·) now captures the TPN’s asymmetric FF-FB integration in its phenomenology with driving and modulating inputs (cf. Fig. 1B, and Fig. 5A). The basal input 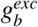 has an impact on the membrane potential even in the absence of apical input 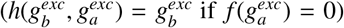 . The apical input, in contrast, only has an impact if it coincides with basal input, due to the multiplicative combination 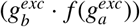. Depending on the model, this apical integration function *f* (·) can, e.g., become a simple summation term (Bouecke et al., 2011; Fig. 2C) or a function that gates contextual cortical FB by thalamic input (Schmid and Neumann, 2025; Fig. 2B).

#### Simplification of the interneuron circuitry

The local circuitry of inhibitory interneurons stabilizes the TPN’s computation and realizes two functions, namely, a pool competition to normalize the excitatory output responses and the control of apical-basal integration.

##### Pool competition

PV cells balance the local drive to an operating regime and perform pool normalization among TPNs (Sect. 3.1.2).

##### Control of apical-basal integration

VIP cells and their interaction with SOM cells (dis-)inhibit the apical compartment and gate the apical-basal integration (Sect. 4 and Fig. 5B,C).

##### Simplification

With the simplification of the apical integration mechanism above, the VIP-SOM system becomes subsumed in the apical integration function *f* (·) (cf. Eq. E.3; see Brosch and Neumann, 2014b for a firing-rate circuit of a cortical column whose feedback-driven dis-inhibition yields such a modulation), while the PV’s pool competition motif remains. As a result, the inhibitory interneuron Eq. 3 reduces to a single, structurally identical inhibitory interneuron equation

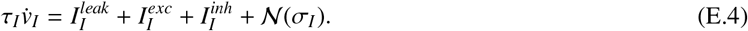

Overall, we arrive at a Mod-E-I model that simplifies the TPN and reduces the system of ODEs from five state variables (Eqs. 1–3) to two (Eqs. E.1, E.4). The above simplifications help contextualize earlier findings in light of the proposed TPN model and provide guidance to match the model’s complexity to the concrete target of investigation. Our investigation shows how the biological detail and rich dynamics can be simplified while still keeping the motifs of asymmetric FF-FB integration and pool competition intact. The Mod-E-I model demonstrably implements these motifs to realize computational strategies for accumulating and disambiguating sensory evidence as in biased competition (cf. Sects. 2.1.1 and 3.1.2).

#### E.2 Simplification of model equations

After performing the simplification steps from the TPN model to the Mod-E-I model (Appendix E.1), finally, the differential equations themselves can be the target of simplification.

##### Numerical integration

To simulate the computational models, the system of ODEs has to be solved to determine its state variables *v*_*i*_(*t*) at every instant in time. The ODE system is typically nonlinear, such that its solution requires computationally expensive numerical integration schemes.

##### Simplification

Often, the exact moment-to-moment variation is not of predominant interest, but rather its functional impact on the resulting computation once the system has arrived at a stable steady state. Mod-E-I models have been found to converge to such stable steady states for suitable parameters (Brosch and Neumann, 2014a). These solutions would require solving the ODE system for its steady state 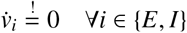. Yet, as the Mod-E-I system is usually nonlinear, the equilibrium solution is not straightforward to obtain, even under static input. Nevertheless, earlier work has shown that approximations to the temporal evolution of the system toward its equilibrium can work quite well. Two relevant variants exist. The first is a fixed-point iteration scheme, which effectively amortizes fine-grained numerical integration of the ODEs by iteratively updating the analytical steady-state equation with current estimates of the state variables (Schmid and Neumann, 2025). The second, further simplified approach performs the processing steps of the model computation in sequence rather than in parallel. It first computes the feedforward integration, then applies the feedback modulation, and finally computes the pool normalization from the previous stage (Brosch and Neumann, 2014a).

#### E.3. Ring networks of Mod-E-I units

To test which network properties the reduction retains, the simulations of the line (Fig. 7, Appendix D.2) and of the bars (Fig. 8) were repeated with rings of Mod-E-I units. Every TPN with its PV, SOM and VIP cells is replaced by an excitatory unit (Eq. E.1) with 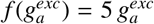. Its inhibition is pooled within the ring from the instantaneous rates 1.3 [*r*_*E*_ − 0.1]_+_ (uniform kernel, *w* = 800*/N*, reversal potential −0.01, gain 10). Its basal conductance sums the FF input and the recurrent excitation (Gaussian kernel, *σ* = *π/*16, peak 4). Its apical conductance sums the external FB input and the contextual projections (weight 1 from each neighbor, 2 at the ends of the line). These pass the dendritic threshold of Eq. 25 at *ρ*_*dend*_ = 0.16, which lies between the outputs of rings with uncertain and with certain FF input. With 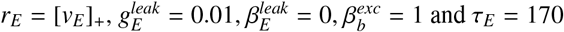 ms, the units relax at their operating point about as fast as the basal compartment of the TPN. Inputs and integration are those of the TPN rings (Tabs. D.1 and A.2). Unlike there, the inputs are not rectified, the potentials start from *v*_*E*_ *∼ U*(0, 0.1), and the bias starts at 0.2 s. The bars run for 0.5 s, and those of 500 and 1000 rings for 4 s.

The Mod-E-I rings reproduce configurations 1 to 4 of the TPN rings (Fig. E.1). Unlike the TPN rings, they also follow a bias that meets certain cues (configuration 5), and ring 0 moves 97 % of the way to the bias direction. They even follow a bias opposite to uncertain cues (configuration 6, Fig. E.2; cf. Fig. D.1), where ring 0 forms a new bump at the bias direction. The multiplicative modulation scales even the weak residual drive at the bias direction. Context that mismatches the FF input therefore steers the population mode of Mod-E-I units, whereas the apical gate of the TPN requires coincidence with basal activity (Tab. E.1). The bars of Mod-E-I rings align as well (Fig. E.3), within 14, 23, 42, 92, 468 and 944 ms for 10 to 1000 rings. This amounts to about 1 ms per ring of bar length, which is much faster than in the TPN network (Fig. 8G), as the apical input of the Mod-E-I unit modulates its basal drive directly. Only the relaxation time of the Mod-E-I unit was set to match that of the basal compartment of the TPN. The propagation speed along the bars was not matched, so that this difference also reflects the parametrization of the two networks.

**Figure E.1:**
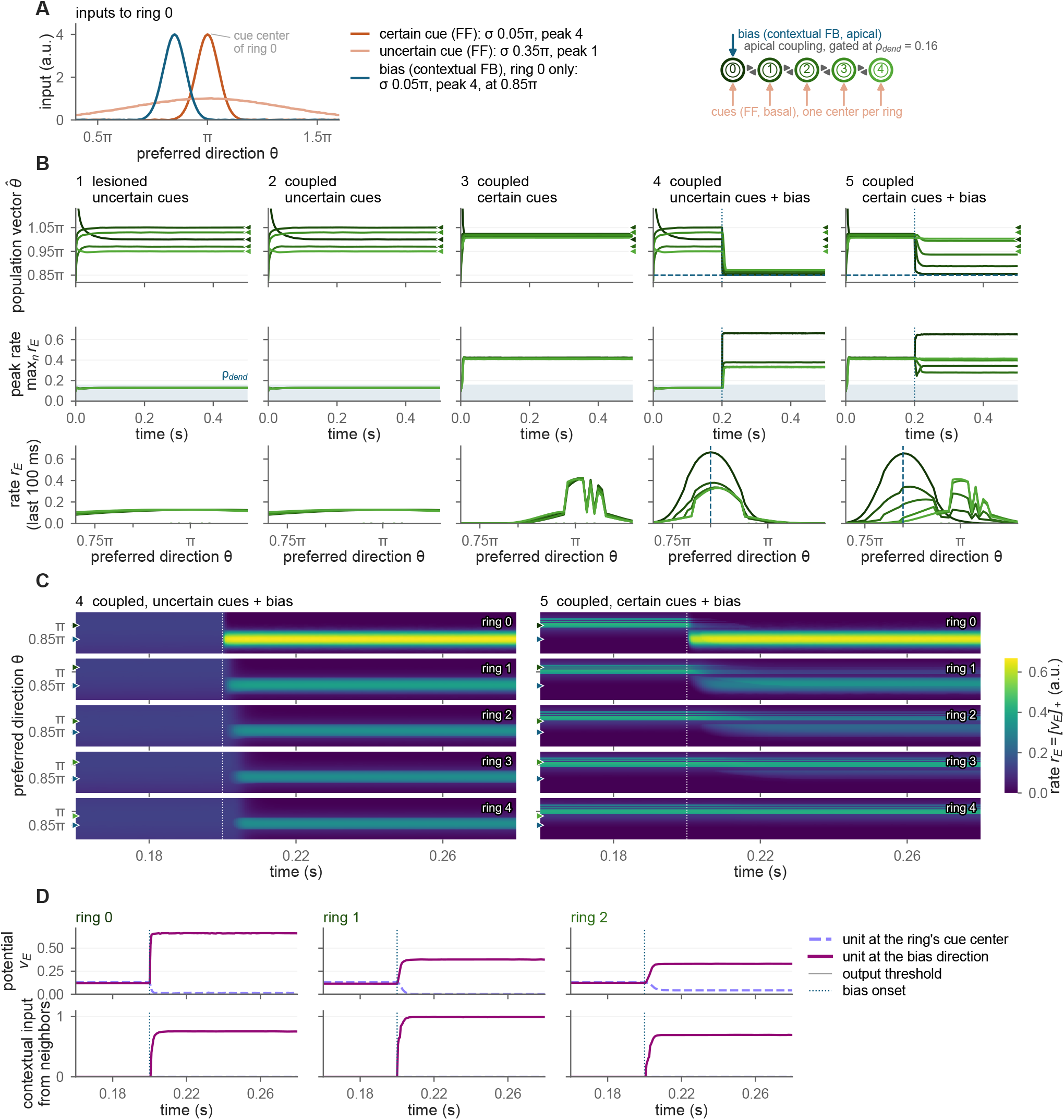
Competition and cooperation in a line of Mod-E-I rings. Panels as in Fig. 7A–D, with the bias from *t* = 0.2 s on. The bottom row of B shows the rate profiles averaged over the last 100 ms of the 0.5-s simulation, and D shows the potential *v*_*E*_ (top) and the contextual input from the neighboring rings (bottom) in configuration 4.

**Figure E.2:**
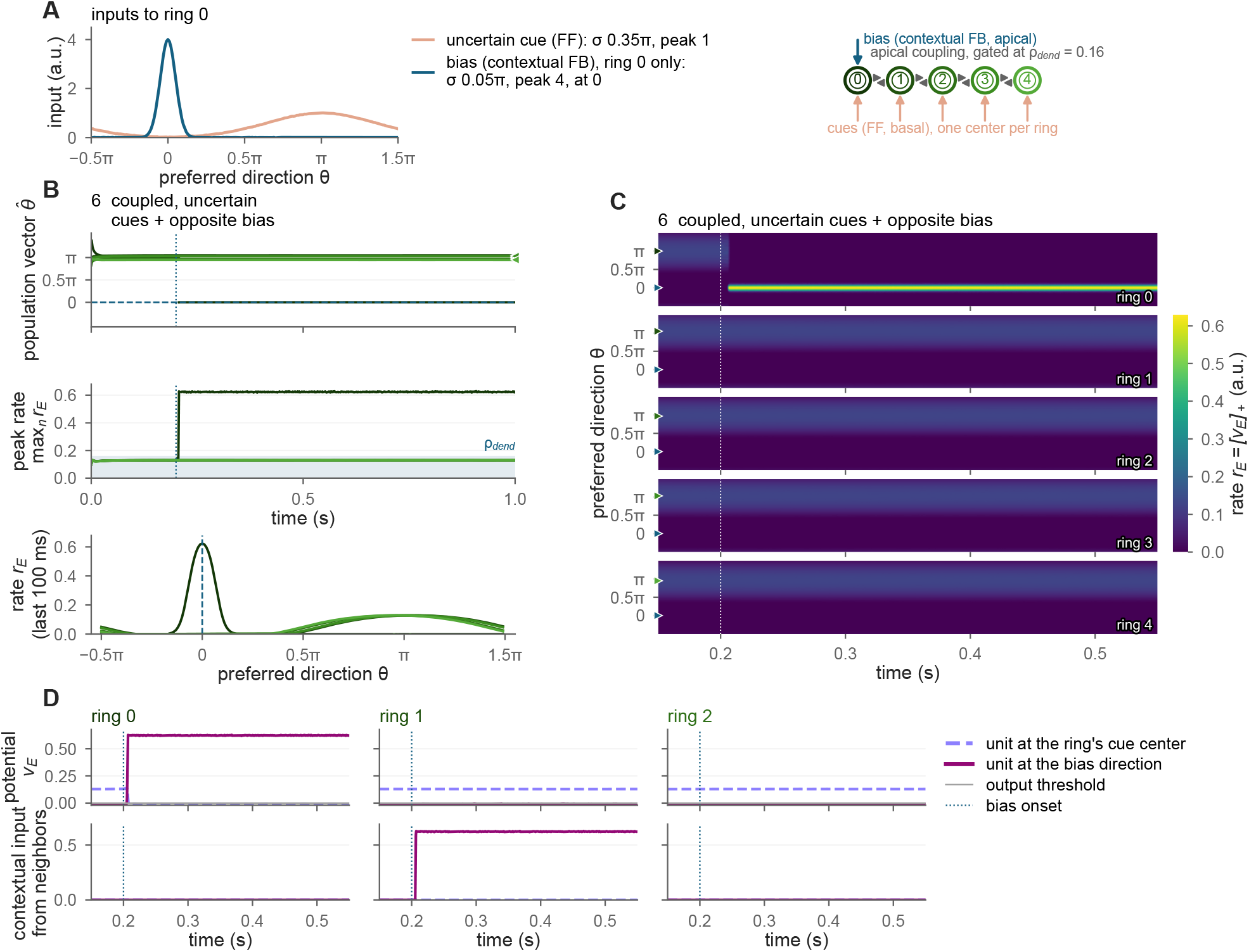
Context opposite to uncertain FF input in a line of Mod-E-I rings. Panels as in Fig. D.1 for configuration 6 of the Mod-E-I rings, with the bias from *t* = 0.2 s on. The bottom row of B shows the rate profiles averaged over the last 100 ms of the 1-s simulation, and D shows the potential *v*_*E*_ (top) and the contextual input from the neighboring rings (bottom) of the unit closest to the ring’s cue center and of the unit closest to the bias direction in rings 0 to 2.

**Figure E.3:**
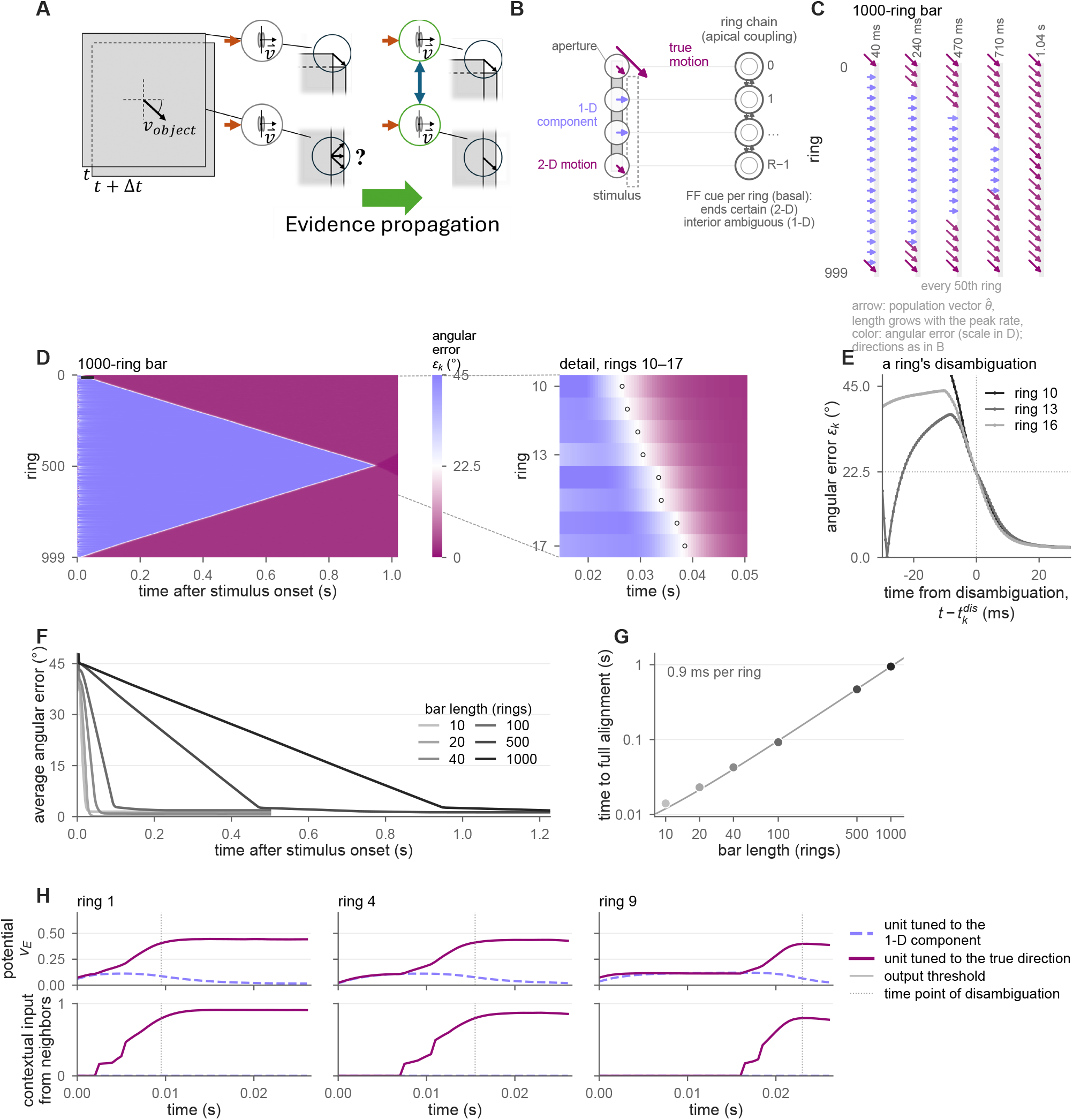
Visual motion integration in a network of Mod-E-I rings. Panels as in Fig. 8A–H for bars of Mod-E-I rings. C to E show a bar of 1000 rings, of which C shows every 50th ring. F and G include the bars of 500 and 1000 rings, with the intact network only in F and logarithmic axes in G. H shows the potential *v*_*E*_ (top) and the contextual input from the neighboring rings (bottom) of the units tuned to the 1-D motion component and to the true motion direction in rings 1, 4 and 9 of a bar of 20 rings.

#### E.4. Task-dependent parallel-sequential and multi-scale evidence integration Model description

**Figure E.4:**
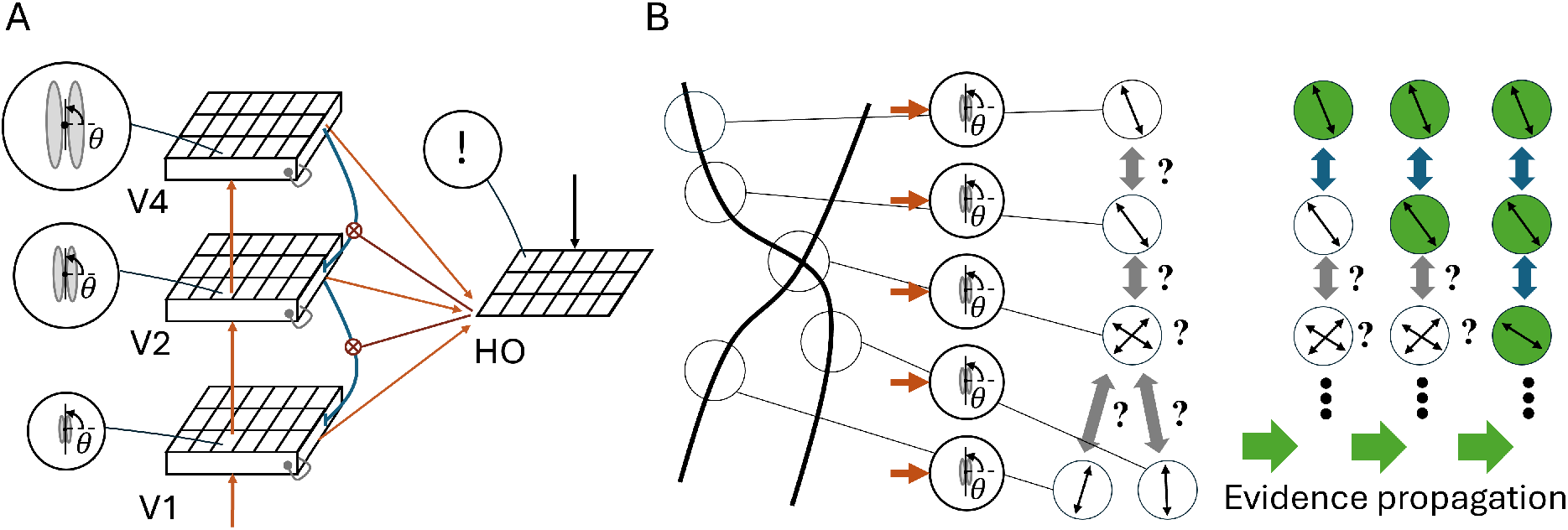
Incremental grouping architecture. (A) Schematic of the architecture of Schmid and Neumann (2025) (cf. their Fig. 3A), which consists of three model visual cortical areas, each with stacked feature maps of simplified TPNs (Fig. 2B), and a higher-order thalamic (HO-Thal) area of point neurons. The model computes oriented contrast information at multiple resolutions across the cortical hierarchy and implements a scale-invariant incremental grouping algorithm. Simplified TPNs together with local inhibitory interneurons perform biased competition and asymmetric feedforward-feedback (FF-FB) integration similar to the more elaborate TPN model proposed here (Fig. 2). Contextual FB from higher to lower cortical stages of compatible oriented contrasts yields an enhancement of the activity of simplified TPNs and signal binding. Thalamic neurons aggregate this enhanced activity from the visual hierarchy and from external (e.g., prefrontal) signals to encode locations of task relevance in a coarse-grained manner. In turn, thalamic projections to the simplified TPNs gate the FF-FB integration process and influence which location can engage in evidence propagation and binding at any given moment in time. See Schmid and Neumann (2025) for details. (B) Incremental grouping in biological neural systems is solved by a decentralized and iterative process of evidence propagation (Roelfsema and Houtkamp, 2011). Initially, with stimulus onset, FF processing extracts evidence about features of the visual stimulus and forms base grouping configurations (circles with orientation arrows). Subsequently, an interaction skeleton is established that expresses compatibilities among local features (bidirectional gray arrows). Afterwards an iterative process takes place, during which compatible neighbors cooperate to propagate task-relevance signals of belonging to the attended target object of interest (blue arrows and green circles). The iterative process adaptively resolves ambiguities dynamically dependent on the already established context of attended elements (crossing element at the center of the configuration).

Schmid and Neumann (2025) present an architecture of simplified TPNs that performs object-based attention in contour tracing by an interplay of corticocortical and thalamocortical loops (Fig. E.4A). The architecture organizes populations of simplified TPNs in a hierarchy of model visual areas V1, V2, and V4 together with an additional higher-order thalamic (HO-Thal) population of point neurons. Model visual areas are mutually connected in an FF-FB manner. The visual hierarchy extracts oriented luminance contrasts at multiple spatial resolutions and forms a scale-space representation. This model presents a realization of the simplified TPN (Fig. 2B). The simplified TPNs are described by two separate compartments integrating FF signals basally and FB signals apically, but possess no bidirectional coupling and perform apical-basal integration via the more abstract FF · (1 + *λ* · FB) modulation transfer function. See Appendix E.1 for details on this simplification. FF-FB integration of the simplified TPNs is additionally gated by input from HO-Thal neurons. In turn, thalamic neurons are driven by task-relevance information originating from across the hierarchy of visual areas and from external input, e.g., from prefrontal areas.

#### Results and insights

The architecture described in Schmid and Neumann (2025) utilizes a network of simplified TPNs to explain parallel-sequential evidence integration in the visual cortex. Such evidence integration capabilities are needed for solving object-based attention tasks commonly investigated by contour tracing paradigms (Fig. E.4B). In contour tracing an iterative binding problem is solved, where local contour elements need to be bound into a coherent object. In experimental tasks, the goal is to determine whether two points on different contour segments belong to the same contour object or not. The computational model explains such iterative binding by attentional enhancement of neural firing rates via the asymmetric FF-FB integration mechanism of the simplified TPN. The dynamics are consistent with findings from neurophysiology and psychophysics (Roelfsema and Houtkamp, 2011). This iterative binding is a form of the evidence propagation scheme as described in Sect. 3. It constitutes a special case in which evidence propagation starts from a single initial location and continues among neighbors. With stimulus onset, an FF sweep extracts a base representation about the feature distribution in the input at multiple scales. Then, context signals about larger neighborhoods are fed back to the more local simplified TPNs in the next lower areas. Crucially, the apical context integration then impacts the basal compartments of only those neurons that also receive coinciding thalamic gating signals. At first, the only signal for opening thalamic gates arises from an attentional seed location provided to thalamic neurons from a driving (e.g., prefrontal) source. This location forms the initial seed point for starting an automated iterative process of FF-FB propagation and integration by an interplay of thalamocortical and corticocortical projection loops. Neurons at the seeded location are allowed to integrate contextual evidence from areas higher up in the hierarchy. The local biased competition, in turn, rewards winning simplified TPNs with enhanced activity. This enhanced activity is again back-projected to thalamic neurons at neighboring locations. These thalamic neurons then also open their gating signals and the FF-FB integration loop proceeds again to label neighboring simplified TPNs by enhanced firing rates until the process converges.

Importantly, the described spreading of attentional modulation establishes a global object representation by sequentially binding local contour elements composed of associated oriented feature items. This iterative process also allows for solving demanding perceptual situations, where the input alone admits more than one interpretation and the intended one requires internal context (Fig. E.5C). Depending on the initially attended location, either the horizontal or the vertical line at the intersection needs to be integrated into the object representation. This requires the iterative integration process to be steerable by task demands.

The dependency of the FF-FB integration process on thalamic neurons allows for flexibly steering the contextual integration by such task-related demands. Depending on the apical gating signal, the representation binds either the one or the other contour object across the crossing and thus respects the task-relevant context in the FF-FB integration process.

These results relate to the findings on the TPN in the main text. First, the coherent labeling of the attended contour corresponds to the global coherence that the networks of TPN rings reach by sequential propagation (Sect. 5). There, the apical amplification of a disambiguated ring passes the evidence on to the next ring, and the time to full alignment grows with bar length (Fig. 8F,G). Likewise, the onset latencies of the response enhancement along a traced contour increase with the distance from the attentional seed in the simplified TPN (Schmid and Neumann, 2025). Second, the higher-order thalamic input of the simplified TPN gates apical-basal integration directly, and Schmid and Neumann (2025) suggested that a more detailed model might mediate this control via apical (dis-)inhibitory circuitry. In the TPN, the VIP-SOM circuit provides such a route and sets the operating regime of apical-basal integration (Fig. 5C). Third, the same circuit also gates learning, because a coincidence tag forms only where FF input coincides with an open apical gate (Appendix C.1). In the flanker task, each TPN thereby learns a stimulus pattern together with the context that co-occurs with it (Fig. 6B). Tonic input to the VIP cells switches this learning on and off (Fig. 6E). A task-dependent gate on apical-basal integration may therefore select both which features are bound into an object and which features are learned in their context.

**Figure E.5:**
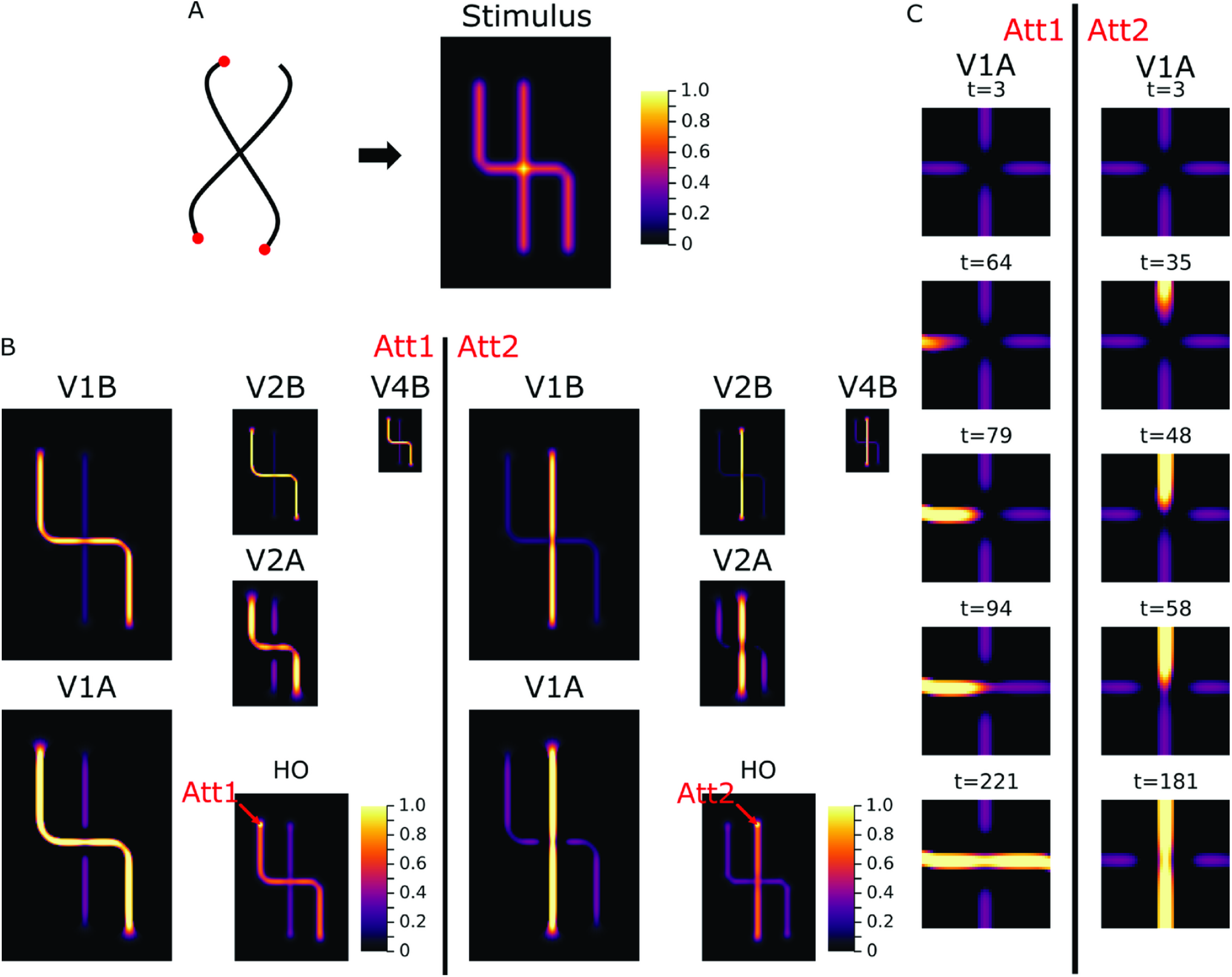
Task-dependent evidence propagation. The model proposed by Schmid and Neumann (2025) dynamically implements a task-dependent evidence propagation algorithm to perform incremental grouping tasks (cf. Fig. E.4). (A) An ambiguous input stimulus consisting of two intersecting curves is fed to the model. (B) Resulting representation after convergence of the iterative process. One of the curves becomes the attended object given a localized starting point. The representation attentively labels retinotopic locations along the curve’s outline by forming a coherent state of amplified responses of the simplified TPNs across the populations. Which of the two curves is attended depends on a task relevance signal provided to the HO population (Att1 vs. Att2; left vs. right column, respectively). The location of one of the lines’ ends is seeded with a task relevance signal. Afterwards an iterative process of evidence propagation takes place (cf. Fig. E.4B). (C) The evidence propagation unfolds over time and dynamically disambiguates between rivaling patterns of connectedness at the intersection point (cf. Fig. E.4B). After stimulus onset, the model extracts local contour information across the hierarchy of visual cortical areas. Apical compartments of the simplified TPNs integrate the compatible contextual FB and represent an interaction skeleton. At the intersection point, the crossing rivaling contours provide ambiguous information and the interaction skeleton formation is locally suppressed. Once the incremental grouping process arrives in the vicinity of the intersection point, the apical representation starts to change. This enhanced apical context helps to resolve the ambiguity at the intersection in favor of collinear contour segments and the evidence propagation continues. See Schmid and Neumann (2025) for details. Heatmap values are normalized to the maximum across populations. VxB and VxA denote the basal and apical compartment activity in areas V1 to V4, and HO the higher-order thalamic area. Panels reproduced from Schmid and Neumann (2025) under the Creative Commons Attribution 4.0 International License (https://creativecommons.org/licenses/by/4.0/).

## Footnotes

1 For ordered populations of cells, such as for retinotopically organized areas in visual information processing, such pattern matching of input can then be expressed as a convolution filtering operation (*I * K*)_*x*_ in domain *x* with input *I*, a kernel *K*, and convolution operator *\**.

2 The leak currents relax the membrane potential toward the reversal potential 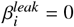, below which the reversal potentials of inhibitory inputs 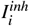 lie, such that inhibitory inputs hyperpolarize the cell.

3 For example, in visual information processing a model population would correspond to a cortical hypercolumn that encodes the direction of local motion at a certain retinotopic location, or, for a ring over [0, *π*), oriented luminance contrasts.

4 Data available from Shai et al. (2015) via https://github.com/ModelDBRepository/180373.

5 See Appendix A.1 for these time constants and Appendix A.3 for a TPN variant that passes sodium- and calcium-like signals between its compartments.

6 Without coincidence tags, the efficacies do not become selective, and the accuracy that an individual network reaches is set by the selectivity of its random initial basal efficacies.

7 Details on time constants and an extended model with separate sodium- and calcium-like exchange terms can be found in Appendices A.1 and A.3.

8 Interestingly, despite its amplifying impact on processing, the apical-to-basal influx component for tag formation (Eq. 15) is described by a rectified difference between the apical and basal potentials, which could be viewed as an apical-basal error term.

9 Learning in large-scale networks of reduced models along these lines is the subject of ongoing work.

## Notes

### Competing Interest Statement

The authors have declared no competing interest.

### Summary of Updates

Revised after peer review. The main changes are: - The two-point neuron (TPN) model now covers both processing and learning. Apical-basal coincidence tags TPNs for a local four-factor learning rule driven by reward, and VIP dis-inhibition gates whether learning takes place. A pool of TPNs learns to use flanker context to decide ambiguous orientations (new Section 4.2, Figures 3 and 6, Appendix C). - New network results with the full TPN. Networks of TPN ring populations disambiguate the motion of bars by propagating evidence at constant speed, which fails when the basal and apical compartments are decoupled (new Section 5.2, Figure 8, Appendix D). - The manuscript was restructured so that the Methods precede the Results. An Appendix in five parts gives further details of the parameters, fits and simulations. - The earlier large-scale simulations with simplified models, which had been published before, were removed from the main text. Appendix E summarizes part of them. - We state more precisely which consequences of backpropagation-activated calcium spike (BAC) firing the model captures and on which time scale. - The Discussion was rewritten. It compares the TPN critically with its reduced versions (Mod-E-I models), supported by new simulations of Mod-E-I ring networks, and with other TPN models, which a new overview figure places by their level of description. - Abstract, Highlights, Introduction and graphical abstract were updated. Figure 1 was partly recolored and redrawn, Figure 5 redrawn with new simulations, and Figure 7 shows new simulations with five input configurations.

## References

Adeel, A., Franco, M., Raza, M., Ahmed, K., 2026. Context-sensitive neocortical neurons transform the effectiveness and efficiency of neural information processing. arXiv preprint arXiv:2207.07338v8. doi:10.48550/arXiv.2207.07338.

Adeel, A., Muzaffar, J., Zia, F., Ahmed, K., Raza, M., Chaudary, E., Riaz, T.B., Saeed, A., 2025. Cooperation is all you need. arXiv preprint arXiv:2305.10449v3. doi:10.48550/arXiv.2305.10449.

Adeli, H., Ahn, S., Kriegeskorte, N., Zelinsky, G., 2023. Affinity-based attention in self-supervised transformers predicts dynamics of object grouping in humans. arXiv preprint arXiv:2306.00294. doi:10.48550/arXiv.2306.00294.

Adelson, E.H., Movshon, J.A., 1982. Phenomenal coherence of moving visual patterns. Nature 300, 523–525. doi:10.1038/300523a0.

Adesnik, H., Bruns, W., Taniguchi, H., Huang, Z.J., Scanziani, M., 2012. A neural circuit for spatial summation in visual cortex. Nature 490, 226–231. doi:10.1038/nature11526.

Amari, S., 1977. Dynamics of pattern formation in lateral-inhibition type neural fields. Biological Cybernetics 27, 77–87. doi:10.1007/BF00337259.

Anastasiades, P.G., Collins, D.P., Carter, A.G., 2021. Mediodorsal and ventromedial thalamus engage distinct L1 circuits in the prefrontal cortex. Neuron 109, 314–330.e4. doi:10.1016/j.neuron.2020.10.031.

Antic, S.D., Zhou, W.L., Moore, A.R., Short, S.M., Ikonomu, K.D., 2010. The decade of the dendritic NMDA spike. Journal of Neuroscience Research 88, 2991–3001. doi:10.1002/jnr.22444.

Aru, J., Siclari, F., Phillips, W.A., Storm, J.F., 2020. Apical drive—A cellular mechanism of dreaming? Neuroscience and Biobehavioral Reviews 119, 440–455. doi:10.1016/j.neubiorev.2020.09.018.

Aru, J., Suzuki, M., Rutiku, R., Larkum, M.E., Bachmann, T., 2019. Coupling the state and contents of consciousness. Frontiers in Systems Neuroscience 13, 43. doi:10.3389/fnsys.2019.00043.

Baronig, M., Legenstein, R., 2024. Context association in pyramidal neurons through local synaptic plasticity in apical dendrites. Frontiers in Neuroscience 17, 1276706. doi:10.3389/fnins.2023.1276706.

Bayerl, P., Neumann, H., 2004. Disambiguating visual motion through contextual feedback modulation. Neural Computation 16, 2041–2066. doi:10.1162/0899766041732404.

Beaulieu-Laroche, L., Toloza, E.H.S., Brown, N.J., Harnett, M.T., 2019. Widespread and highly correlated somatodendritic activity in cortical layer 5 neurons. Neuron 103, 235–241.e4. doi:10.1016/j.neuron.2019.05.014.

Ben-Yishai, R., Bar-Or, R.L., Sompolinsky, H., 1995. Theory of orientation tuning in visual cortex. Proceedings of the National Academy of Sciences 92, 3844–3848. doi:10.1073/pnas.92.9.3844.

Bittner, K.C., Milstein, A.D., Grienberger, C., Romani, S., Magee, J.C., 2017. Behavioral time scale synaptic plasticity underlies CA1 place fields. Science 357, 1033–1036. doi:10.1126/science.aan3846.

Born, R.T., Bradley, D.C., 2005. Structure and function of visual area MT. Annual Review of Neuroscience 28, 157–189. doi:10.1146/annurev.neuro.26.041002.131052.

Born, R.T., Pack, C.C., Ponce, C.R., Yi, S., 2006. Temporal evolution of 2-dimensional direction signals used to guide eye movements. Journal of Neurophysiology 95, 284–300. doi:10.1152/jn.01329.2004.

Bortone, D.S., Olsen, S.R., Scanziani, M., 2014. Translaminar inhibitory cells recruited by layer 6 corticothalamic neurons suppress visual cortex. Neuron 82, 474–485. doi:10.1016/j.neuron.2014.02.021.

Bouecke, J.D., Tlapale, E., Kornprobst, P., Neumann, H., 2011. Neural mechanisms of motion detection, integration, and segregation: From biology to artificial image processing systems. EURASIP Journal on Advances in Signal Processing 2011, 781561. doi:10.1155/2011/781561.

Bradbury, J., Frostig, R., Hawkins, P., Johnson, M.J., Katariya, Y., Leary, C., Maclaurin, D., Necula, G., Paszke, A., VanderPlas, J., Wanderman-Milne, S., Zhang, Q., 2018. JAX: Composable transformations of Python+NumPy programs. https://github.com/jax-ml/jax.

Branch, M.A., Coleman, T.F., Li, Y., 1999. A subspace, interior, and conjugate gradient method for large-scale bound-constrained minimization problems. SIAM Journal on Scientific Computing 21, 1–23. doi:10.1137/S1064827595289108.

Britten, K.H., Shadlen, M.N., Newsome, W.T., Movshon, J.A., 1992. The analysis of visual motion: A comparison of neuronal and psychophysical performance. The Journal of Neuroscience 12, 4745–4765. doi:10.1523/JNEUR0SCI.12-12-04745.1992.

Brosch, T., Neumann, H., 2014a. Computing with a canonical neural circuits model with pool normalization and modulating feedback. Neural Computation 26, 2735–2789. doi:10.1162/NEC0_a_00675.

Brosch, T., Neumann, H., 2014b. Interaction of feedforward and feedback streams in visual cortex in a firing-rate model of columnar computations. Neural Networks 54, 11–16. doi:10.1016/j.neunet.2014.02.005.

Brosch, T., Neumann, H., Roelfsema, P.R., 2015. Reinforcement learning of linking and tracing contours in recurrent neural networks. PLOS Computational Biology 11, e1004489. doi:10.1371/journal.pcbi.1004489.

Brown, J., Bullock, D., Grossberg, S., 1999. How the basal ganglia use parallel excitatory and inhibitory learning pathways to selectively respond to unexpected rewarding cues. The Journal of Neuroscience 19, 10502–10511. doi:10.1523/JNEUR0SCI.19-23-10502.1999.

Campagnola, L., Seeman, S.C., Chartrand, T., Kim, L., Hoggarth, A., Gamlin, C., Ito, S., Trinh, J., Davoudian, P., Radaelli, C., Kim, M.H., Hage, T., Braun, T., Alfiler, L., Andrade, J., Bohn, P., Dalley, R., Henry, A., Kebede, S., Mukora, A., Sandman, D., Williams, G., Larsen, R., Teeter, C., Daigle, T.L., Berry, K., Dotson, N., Enstrom, R., Gorham, M., Hupp, M., Dingman Lee, S., Ngo, K., Nicovich, P.R., Potekhina, L., Ransford, S., Gary, A., Goldy, J., McMillen, D., Pham, T., Tieu, M., Siverts, L., Walker, M., Farrell, C., Schroedter, M., Slaughterbeck, C., Cobb, C., Ellenbogen, R., Gwinn, R.P., Keene, C.D., Ko, A.L., Ojemann, J.G., Silbergeld, D.L., Carey, D., Casper, T., Crichton, K., Clark, M., Dee, N., Ellingwood, L., Gloe, J., Kroll, M., Sulc, J., Tung, H., Wadhwani, K., Brouner, K., Egdorf, T., Maxwell, M., McGraw, M., Pom, C.A., Ruiz, A., Bomben, J., Feng, D., Hejazinia, N., Shi, S., Szafer, A., Wakeman, W., Phillips, J., Bernard, A., Esposito, L., D’Orazi, F.D., Sunkin, S., Smith, K., Tasic, B., Arkhipov, A., Sorensen, S., Lein, E., Koch, C., Murphy, G., Zeng, H., Jarsky, T., 2022. Local connectivity and synaptic dynamics in mouse and human neocortex. Science 375, eabj5861. doi:10.1126/science.abj5861.

Capone, C., Lupo, C., Muratore, P., Paolucci, P.S., 2023. Beyond spiking networks: The computational advantages of dendritic amplification and input segregation. Proceedings of the National Academy of Sciences 120, e2220743120. doi:10.1073/pnas.2220743120.

Carandini, M., Heeger, D.J., 2012. Normalization as a canonical neural computation. Nature Reviews Neuroscience 13, 51–62. doi:10.1038/nrn3136.

Chen, Y., Douglas, H., Medina, B.J., Olarinre, M., Siegle, J.H., Kass, R.E., 2022. Population burst propagation across interacting areas of the brain. Journal of Neurophysiology 128, 1578–1592. doi:10.1152/jn.00066.2022.

Chilton, J., 2020. Brain outline. Zenodo, 10.5281/zenodo.3925989. Licensed under CC BY 4.0, https://creativecommons.org/licenses/by/4.0/.

Cossell, L., Iacaruso, M.F., Muir, D.R., Houlton, R., Sader, E.N., Ko, H., Hofer, S.B., Mrsic-Flogel, T.D., 2015. Functional organization of excitatory synaptic strength in primary visual cortex. Nature 518, 399–403. doi:10.1038/nature14182.

Darlow, L., Regan, C., Risi, S., Seely, J., Jones, L., 2025. Continuous Thought Machines, in: Advances in Neural Information Processing Systems 38, pp. 24432–24478. doi:10.52202/085713-0728.

Desimone, R., Duncan, J., 1995. Neural mechanisms of selective visual attention. Annual Review of Neuroscience 18, 193–222. doi:10.1146/annurev.ne.18.030195.001205.

Dormand, J.R., Prince, P.J., 1980. A family of embedded Runge-Kutta formulae. Journal of Computational and Applied Mathematics 6, 19–26. doi:10.1016/0771-050X(80)90013-3.

Doron, G., Shin, J.N., Takahashi, N., Drüke, M., Bocklisch, C., Skenderi, S., de Mont, L., Toumazou, M., Ledderose, J., Brecht, M., Naud, R., Larkum, M.E., 2020. Perirhinal input to neocortical layer 1 controls learning. Science 370, eaaz3136. doi:10.1126/science.aaz3136.

Felleman, D.J., Van Essen, D.C., 1991. Distributed hierarchical processing in the primate cerebral cortex. Cerebral Cortex 1, 1–47. doi:10.1093/cercor/1.1.1.

Francioni, V., Padamsey, Z., Rochefort, N.L., 2019. High and asymmetric somato-dendritic coupling of V1 layer 5 neurons independent of visual stimulation and locomotion. eLife 8, e49145. doi:10.7554/eLife.49145.

Francioni, V., Tang, V.D., Toloza, E.H.S., Ding, Z., Brown, N.J., Harnett, M.T., 2026. Vectorized instructive signals in cortical dendrites. Nature 652, 1254–1263. doi:10.1038/s41586-026-10190-7.

Frémaux, N., Gerstner, W., 2016. Neuromodulated spike-timing-dependent plasticity, and theory of three-factor learning rules. Frontiers in Neural Circuits 9, 85. doi:10.3389/fncir.2015.00085.

Friedenberger, Z., Harkin, E., Tóth, K., Naud, R., 2023. Silences, spikes and bursts: Three-part knot of the neural code. The Journal of Physiology 601, 5165–5193. doi:10.1113/JP281510.

Furutachi, S., Franklin, A.D., Aldea, A.M., Mrsic-Flogel, T.D., Hofer, S.B., 2024. Cooperative thalamocortical circuit mechanism for sensory prediction errors. Nature 633, 398–406. doi:10.1038/s41586-024-07851-w.

Gambino, F., Pagès, S., Kehayas, V., Baptista, D., Tatti, R., Carleton, A., Holtmaat, A., 2014. Sensory-evoked LTP driven by dendritic plateau potentials in vivo. Nature 515, 116–119. doi:10.1038/nature13664.

Georgopoulos, A.P., Schwartz, A.B., Kettner, R.E., 1986. Neuronal population coding of movement direction. Science 233, 1416–1419. doi:10.1126/science.3749885.

Gerstner, W., Lehmann, M., Liakoni, V., Corneil, D., Brea, J., 2018. Eligibility traces and plasticity on behavioral time scales: Experimental support of neoHebbian three-factor learning rules. Frontiers in Neural Circuits 12, 53. doi:10.3389/fncir.2018.00053.

Gilbert, C.D., Li, W., 2013. Top-down influences on visual processing. Nature Reviews Neuroscience 14, 350–363. doi:10.1038/nrn3476.

Graham, B.P., Kay, J.W., 2026. Inhibitory and neuromodulatory control of two-pathway signal integration in a model neocortical pyramidal cell. Current Research in Neurobiology 11, 100166. doi:10.1016/j.crneur.2026.100166.

Graham, B.P., Kay, J.W., Phillips, W.A., 2025. Context-sensitive processing in a model neocortical pyramidal cell with two sites of input integration. Neural Computation 37, 588–634. doi:10.1162/neco_a_01739.

Greff, K., van Steenkiste, S., Schmidhuber, J., 2020. On the binding problem in artificial neural networks. arXiv preprint arXiv:2012.05208. doi:10.48550/arXiv.2012.05208.

Grossberg, S., 1973. Contour enhancement, short term memory, and constancies in reverberating neural networks. Studies in Applied Mathematics 52, 213–257. doi:10.1002/sapm1973523213.

Grossberg, S., 1976. Adaptive pattern classification and universal recoding: I. Parallel development and coding of neural feature detectors. Biological Cybernetics 23, 121–134. doi:10.1007/BF00344744.

Grossberg, S., Mingolla, E., 1985. Neural dynamics of form perception: Boundary completion, illusory figures, and neon color spreading. Psychological Review 92, 173–211. doi:10.1037/0033-295X.92.2.173.

Halassa, M.M. (Ed.), 2022. The Thalamus. Cambridge University Press, Cambridge. doi:10.1017/9781108674287.

Halassa, M.M., Kastner, S., 2017. Thalamic functions in distributed cognitive control. Nature Neuroscience 20, 1669–1679. doi:10.1038/s41593-017-0020-1.

Hangya, B., Pi, H.J., Kvitsiani, D., Ranade, S.P., Kepecs, A., 2014. From circuit motifs to computations: Mapping the behavioral repertoire of cortical interneurons. Current Opinion in Neurobiology 26, 117–124. doi:10.1016/j.conb.2014.01.007.

Harris, C.R., Millman, K.J., van der Walt, S.J., Gommers, R., Virtanen, P., Cournapeau, D., Wieser, E., Taylor, J., Berg, S., Smith, N.J., et al., 2020. Array programming with NumPy. Nature 585, 357–362. doi:10.1038/s41586-020-2649-2.

Harris, K.D., Mrsic-Flogel, T.D., 2013. Cortical connectivity and sensory coding. Nature 503, 51–58. doi:10.1038/nature12654.

Hay, E., Hill, S., Schürmann, F., Markram, H., Segev, I., 2011. Models of neocortical layer 5b pyramidal cells capturing a wide range of dendritic and perisomatic active properties. PLOS Computational Biology 7, e1002107. doi:10.1371/journal.pcbi.1002107.

Hensch, T.K., 2005. Critical period plasticity in local cortical circuits. Nature Reviews Neuroscience 6, 877–888. doi:10.1038/nrn1787.

Hertäg, L., Sprekeler, H., 2019. Amplifying the redistribution of somato-dendritic inhibition by the interplay of three interneuron types. PLOS Computational Biology 15, e1006999. doi:10.1371/journal.pcbi.1006999.

Hertäg, L., Sprekeler, H., 2020. Learning prediction error neurons in a canonical interneuron circuit. eLife 9, e57541. doi:10.7554/eLife.57541.

Hunter, J.D., 2007. Matplotlib: A 2D graphics environment. Computing in Science & Engineering 9, 90–95. doi:10.1109/MCSE.2007.55.

Hupé, J.M., James, A.C., Payne, B.R., Lomber, S.G., Girard, P., Bullier, J., 1998. Cortical feedback improves discrimination between figure and background by V1, V2 and V3 neurons. Nature 394, 784–787. doi:10.1038/29537.

Islah, N., Etter, G., Tugsbayar, M., Gurbuz, B.T., Richards, B., Muller, E.B., 2025. Learning to combine top-down context and feed-forward representations under ambiguity with apical and basal dendrites. Cerebral Cortex 35, bhaf134. doi:10.1093/cercor/bhaf134.

Iyer, A., Grewal, K., Velu, A., Souza, L.O., Forest, J., Ahmad, S., 2022. Avoiding catastrophe: Active dendrites enable multi-task learning in dynamic environments. Frontiers in Neurorobotics 16, 846219. doi:10.3389/fnbot.2022.846219.

Izhikevich, E.M., 2007. Solving the distal reward problem through linkage of STDP and dopamine signaling. Cerebral Cortex 17, 2443–2452. doi:10.1093/cercor/bhl152.

Jordan, R., Keller, G.B., 2020. Opposing influence of top-down and bottom-up input on excitatory layer 2/3 neurons in mouse primary visual cortex. Neuron 108, 1194–1206.e5. doi:10.1016/j.neuron.2020.09.024.

Kampa, B.M., Letzkus, J.J., Stuart, G.J., 2006. Requirement of dendritic calcium spikes for induction of spike-timing-dependent synaptic plasticity. The Journal of Physiology 574, 283–290. doi:10.1113/jphysiol.2006.111062.

Kapadia, M.K., Ito, M., Gilbert, C.D., Westheimer, G., 1995. Improvement in visual sensitivity by changes in local context: Parallel studies in human observers and in V1 of alert monkeys. Neuron 15, 843–856. doi:10.1016/0896–6273(95)90175-2.

Kaup, K.K., Hidalgo Jiménez, J., Aru, J., 2026. Cellular mechanisms of serotonergic psychedelics – apical hypercontextualisation. Neuroscience and Biobehavioral Reviews 189, 106876. doi:10.1016/j.neubiorev.2026.106876.

Kay, J., Phillips, W.A., 1997. Activation functions, computational goals, and learning rules for local processors with contextual guidance. Neural Computation 9, 895–910. doi:10.1162/neco.1997.9.4.895.

Kay, J.W., Graham, B.P., 2026. Modulation and integration in a local processor having input from two distinct sources. arXiv preprint arXiv:2610.05360. doi:10.48550/arXiv.2610.05360.

Kay, J.W., Phillips, W.A., 2011. Coherent Infomax as a computational goal for neural systems. Bulletin of Mathematical Biology 73, 344–372. doi:10.1007/s11538-010-9564-x.

Kay, J.W., Phillips, W.A., 2020. Contextual modulation in mammalian neocortex is asymmetric. Symmetry 12, 815. doi:10.3390/sym12050815.

Kerlin, A.M., Andermann, M.L., Berezovskii, V.K., Reid, R.C., 2010. Broadly tuned response properties of diverse inhibitory neuron subtypes in mouse visual cortex. Neuron 67, 858–871. doi:10.1016/j.neuron.2010.08.002.

Khan, A.G., Hofer, S.B., 2018. Contextual signals in visual cortex. Current Opinion in Neurobiology 52, 131–138. doi:10.1016/j.conb.2018.05.003.

Khona, M., Fiete, I.R., 2022. Attractor and integrator networks in the brain. Nature Reviews Neuroscience 23, 744–766. doi:10.1038/s41583-022-00642-0.

Kidger, P., 2021. On Neural Differential Equations. Ph.D. thesis. University of Oxford. arXiv:2202.02435.

Kirchberger, L., Mukherjee, S., Schnabel, U.H., van Beest, E.H., Barsegyan, A., Levelt, C.N., Heimel, J.A., Lorteije, J.A.M., van der Togt, C., Self, M.W., Roelfsema, P.R., 2021. The essential role of recurrent processing for figure-ground perception in mice. Science Advances 7, eabe1833. doi:10.1126/sciadv.abe1833.

Ko, H., Hofer, S.B., Pichler, B., Buchanan, K.A., Sjöström, P.J., Mrsic-Flogel, T.D., 2011. Functional specificity of local synaptic connections in neocortical networks. Nature 473, 87–91. doi:10.1038/nature09880.

Koch, C., Rapp, M., Segev, I., 1996. A brief history of time (constants). Cerebral Cortex 6, 93–101. doi:10.1093/cercor/6.2.93.

Lam, S.K., Pitrou, A., Seibert, S., 2015. Numba: A LLVM-based Python JIT compiler, in: Proceedings of the Second Workshop on the LLVM Compiler Infrastructure in HPC, ACM. pp. 1–6. doi:10.1145/2833157.2833162.

Lamme, V.A.F., Roelfsema, P.R., 2000. The distinct modes of vision offered by feedforward and recurrent processing. Trends in Neurosciences 23, 571–579. doi:10.1016/S0166-2236(00)01657-X.

Lamme, V.A.F., Zipser, K., Spekreijse, H., 1998. Figure-ground activity in primary visual cortex is suppressed by anesthesia. Proceedings of the National Academy of Sciences 95, 3263–3268. doi:10.1073/pnas.95.6.3263.

Larkum, M., 2013. A cellular mechanism for cortical associations: An organizing principle for the cerebral cortex. Trends in Neurosciences 36, 141–151. doi:10.1016/j.tins.2012.11.006.

Larkum, M.E., Nevian, T., Sandler, M., Polsky, A., Schiller, J., 2009. Synaptic integration in tuft dendrites of layer 5 pyramidal neurons: A new unifying principle. Science 325, 756–760. doi:10.1126/science.1171958.

Larkum, M.E., Senn, W., Lüscher, H.R., 2004. Top-down dendritic input increases the gain of layer 5 pyramidal neurons. Cerebral Cortex 14, 1059–1070. doi:10.1093/cercor/bhh065.

Larkum, M.E., Zhu, J.J., Sakmann, B., 1999. A new cellular mechanism for coupling inputs arriving at different cortical layers. Nature 398, 338–341. doi:10.1038/18686.

Larsen, B.W., Druckmann, S., 2022. Towards a more general understanding of the algorithmic utility of recurrent connections. PLOS Computational Biology 18, e1010227. doi:10.1371/journal.pcbi.1010227.

Linsley, D., Kim, J., Veerabadran, V., Windolf, C., Serre, T., 2018. Learning long-range spatial dependencies with horizontal gated recurrent units, in: Bengio, S., Wallach, H., Larochelle, H., Grauman, K., Cesa-Bianchi, N., Garnett, R. (Eds.), Advances in Neural Information Processing Systems 31, Curran Associates, Inc. URL: https://proceedings.neurips.cc/paper_files/paper/2018/file/ec8956637a99787bd197eacd77acce5e-Paper.pdf.

Löhr, M.P.R., Schmid, D., Neumann, H., 2019. Motion integration and disambiguation by spiking V1-MT-MSTl feedforward-feedback interaction, in: 2019 International Joint Conference on Neural Networks (IJCNN), IEEE, Budapest, Hungary. pp. 1–9. doi:10.1109/IJCNN.2019.8852029.

Marvan, T., Phillips, W.A., 2024. Cellular mechanisms of cooperative context-sensitive predictive inference. Current Research in Neurobiology 6, 100129. doi:10.1016/j.crneur.2024.100129.

Maryada De Luca, C., Rubino, A., Wen, C., Cartiglia, M., Fodorut, I.I., Payvand, M., Indiveri, G., 2025. A canonical cortical electronic circuit for neuromorphic intelligence. bioRxiv preprint 2025.03.28.646019. doi:10.1101/2025.03.28.646019.

Mediano, P.A.M., Rosas, F.E., Luppi, A.I., Carhart-Harris, R.L., Bor, D., Seth, A.K., Barrett, A.B., 2025. Toward a unified taxonomy of information dynamics via Integrated Information Decomposition. Proceedings of the National Academy of Sciences 122, e2423297122. doi:10.1073/pnas.2423297122.

Mollard, S., Wacongne, C., Bohte, S.M., Roelfsema, P.R., 2024. Recurrent neural networks that learn multistep visual routines with reinforcement learning. PLOS Computational Biology 20, e1012030. doi:10.1371/journal.pcbi.1012030.

Munn, B.R., Müller, E.J., Aru, J., Whyte, C.J., Gidon, A., Larkum, M.E., Shine, J.M., 2023. A thalamocortical substrate for integrated information via critical synchronous bursting. Proceedings of the National Academy of Sciences 120, e2308670120. doi:10.1073/pnas.2308670120.

Murayama, M., Pérez-Garci, E., Nevian, T., Bock, T., Senn, W., Larkum, M.E., 2009. Dendritic encoding of sensory stimuli controlled by deep cortical interneurons. Nature 457, 1137–1141. doi:10.1038/nature07663.

Naud, R., Sprekeler, H., 2018. Sparse bursts optimize information transmission in a multiplexed neural code. Proceedings of the National Academy of Sciences 115, E6329–E6338. doi:10.1073/pnas.1720995115.

Naumann, L.B., Hertäg, L., Müller, J., Letzkus, J.J., Sprekeler, H., 2025. Layer-specific control of inhibition by NDNF interneurons. Proceedings of the National Academy of Sciences 122, e2408966122. doi:10.1073/pnas.2408966122.

Nelder, J.A., Mead, R., 1965. A simplex method for function minimization. The Computer Journal 7, 308–313. doi:10.1093/comjnl/7.4.308.

Newsome, W.T., Britten, K.H., Movshon, J.A., 1989. Neuronal correlates of a perceptual decision. Nature 341, 52–54. doi:10.1038/341052a0.

Niell, C.M., Scanziani, M., 2021. How cortical circuits implement cortical computations: Mouse visual cortex as a model. Annual Review of Neuroscience 44, 517–546. doi:10.1146/annurev-neuro-102320-085825.

Pack, C.C., Berezovskii, V.K., Born, R.T., 2001. Dynamic properties of neurons in cortical area MT in alert and anaesthetized macaque monkeys. Nature 414, 905–908. doi:10.1038/414905a.

Pack, C.C., Born, R.T., 2001. Temporal dynamics of a neural solution to the aperture problem in visual area MT of macaque brain. Nature 409, 1040–1042. doi:10.1038/35059085.

Pastorelli, E., Yegenoglu, A., Kolodziej, N., Wybo, W., Simula, F., Diaz-Pier, S., Storm, J.F., Paolucci, P.S., 2025. Simplified two-compartment neuron with calcium dynamics capturing brain-state specific apical-amplification, -isolation and -drive. Frontiers in Computational Neuroscience 19, 1566196. doi:10.3389/fncom.2025.1566196.

Payeur, A., Guerguiev, J., Zenke, F., Richards, B.A., Naud, R., 2021. Burst-dependent synaptic plasticity can coordinate learning in hierarchical circuits. Nature Neuroscience 24, 1010–1019. doi:10.1038/s41593-021-00857-x.

Pfeffer, C.K., Xue, M., He, M., Huang, Z.J., Scanziani, M., 2013. Inhibition of inhibition in visual cortex: The logic of connections between molecularly distinct interneurons. Nature Neuroscience 16, 1068–1076. doi:10.1038/nn.3446.

Phillips, W.A., 2023. The Cooperative Neuron: Cellular Foundations of Mental Life. Oxford University Press, Oxford. doi:10.1093/oso/9780198876984.001.0001.

Phillips, W.A., Kay, J., Smyth, D., 1995. The discovery of structure by multi-stream networks of local processors with contextual guidance. Network: Computation in Neural Systems 6, 225–246. doi:10.1088/0954-898X_6_2_005.

Pi, H.J., Hangya, B., Kvitsiani, D., Sanders, J.I., Huang, Z.J., Kepecs, A., 2013. Cortical interneurons that specialize in disinhibitory control. Nature 503, 521–524. doi:10.1038/nature12676.

Polat, U., Mizobe, K., Pettet, M.W., Kasamatsu, T., Norcia, A.M., 1998. Collinear stimuli regulate visual responses depending on cell’s contrast threshold. Nature 391, 580–584. doi:10.1038/35372.

Polat, U., Sagi, D., 1993. Lateral interactions between spatial channels: Suppression and facilitation revealed by lateral masking experiments. Vision Research 33, 993–999. doi:10.1016/0042-6989(93)90081-7.

Purves, D., Augustine, G.J., Fitzpatrick, D., Hall, W.C., LaMantia, A.S., Mooney, R.D., Platt, M.L., White, L.E. (Eds.), 2018. Neuroscience. 6th ed., Sinauer Associates /Oxford University Press, New York.

Roelfsema, P.R., 2023. Solving the binding problem: Assemblies form when neurons enhance their firing rate—they don’t need to oscillate or synchronize. Neuron 111, 1003–1019. doi:10.1016/j.neuron.2023.03.016.

Roelfsema, P.R., Holtmaat, A., 2018. Control of synaptic plasticity in deep cortical networks. Nature Reviews Neuroscience 19, 166–180. doi:10.1038/nrn.2018.6.

Roelfsema, P.R., Houtkamp, R., 2011. Incremental grouping of image elements in vision. Attention, Perception, & Psychophysics 73, 2542–2572. doi:10.3758/s13414-011-0200-0.

Roelfsema, P.R., van Ooyen, A., 2005. Attention-gated reinforcement learning of internal representations for classification. Neural Computation 17, 2176–2214. doi:10.1162/0899766054615699.

Sacramento, J., Ponte Costa, R., Bengio, Y., Senn, W., 2018. Dendritic cortical microcircuits approximate the backpropagation algorithm, in: Bengio, S., Wallach, H., Larochelle, H., Grauman, K., Cesa-Bianchi, N., Garnett, R. (Eds.), Advances in Neural Information Processing Systems 31, Curran Associates, Inc. URL: https://proceedings.neurips.cc/paper_files/paper/2018/file/1dc3a89d0d440ba31729b0ba74b93a33-Paper.pdf.

Salehi, S., Lei, J., Benjamin, A.S., Müller, K.R., Kording, K.P., 2026. Modeling attention and binding in the brain through bidirectional recurrent gating. Nature Communications 17, 4072. doi:10.1038/s41467-026-72146-9.

Schmid, D., Jarvers, C., Neumann, H., 2023. Canonical circuit computations for computer vision. Biological Cybernetics 117, 299–329. doi:10.1007/s00422-023-00966-9.

Schmid, D., Neumann, H., 2025. A model of thalamo-cortical interaction for incremental binding in mental contourtracing. PLOS Computational Biology 21, e1012835. doi:10.1371/journal.pcbi.1012835.

Schultz, W., Dayan, P., Montague, P.R., 1997. A neural substrate of prediction and reward. Science 275, 1593–1599. doi:10.1126/science.275.5306.1593.

Schuman, B., Machold, R.P., Hashikawa, Y., Fuzik, J., Fishell, G.J., Rudy, B., 2019. Four unique interneuron populations reside in neocortical layer 1. The Journal of Neuroscience 39, 125–139. doi:10.1523/JNEUR0SCI.1613-18.2018.

Shai, A.S., Anastassiou, C.A., Larkum, M.E., Koch, C., 2015. Physiology of layer 5 pyramidal neurons in mouse primary visual cortex: Coincidence detection through bursting. PLOS Computational Biology 11, e1004090. doi:10.1371/journal.pcbi.1004090.

Silverstein, S.M., Zolnik, T., Larkum, M., Thompson, J.L., 2026. Apical integration theory provides a neural basis for understanding disorganization and consciousness in the schizophrenias. Current Research in Neurobiology 11, 100165. doi:10.1016/j.crneur.2026.100165.

Smith, M.A., Majaj, N.J., Movshon, J.A., 2005. Dynamics of motion signaling by neurons in macaque area MT. Nature Neuroscience 8, 220–228. doi:10.1038/nn1382.

Storm, J.F., Klink, P.C., Aru, J., Senn, W., Goebel, R., Pigorini, A., Avanzini, P., Vanduffel, W., Roelfsema, P.R., Massimini, M., Larkum, M.E., Pennartz, C.M.A., 2024. An integrative, multiscale view on neural theories of consciousness. Neuron 112, 1531–1552. doi:10.1016/j.neuron.2024.02.004.

Strang, G., 1968. On the construction and comparison of difference schemes. SIAM Journal on Numerical Analysis 5, 506–517. doi:10.1137/0705041.

Suzuki, M., Larkum, M.E., 2020. General anesthesia decouples cortical pyramidal neurons. Cell 180, 666–676.e13. doi:10.1016/j.cell.2020.01.024.

Takahashi, H., Magee, J.C., 2009. Pathway interactions and synaptic plasticity in the dendritic tuft regions of CA1 pyramidal neurons. Neuron 62, 102–111. doi:10.1016/j.neuron.2009.03.007.

Takahashi, N., Ebner, C., Sigl-Glöckner, J., Moberg, S., Nierwetberg, S., Larkum, M.E., 2020. Active dendritic currents gate descending cortical outputs in perception. Nature Neuroscience 23, 1277–1285. doi:10.1038/s41593-020-0677-8.

Takahashi, N., Oertner, T.G., Hegemann, P., Larkum, M.E., 2016. Active cortical dendrites modulate perception. Science 354, 1587–1590. doi:10.1126/science.aah6066.

Tay, Y., Dehghani, M., Abnar, S., Shen, Y., Bahri, D., Pham, P., Rao, J., Yang, L., Ruder, S., Metzler, D., 2021. Long Range Arena: A benchmark for efficient transformers, in: International Conference on Learning Representations (ICLR). URL: https://openreview.net/forum?id=qVyeW-grC2k.

Tremblay, R., Lee, S., Rudy, B., 2016. GABAergic interneurons in the neocortex: From cellular properties to circuits. Neuron 91, 260–292. doi:10.1016/j.neuron.2016.06.033.

Tsui, J.M.G., Hunter, J.N., Born, R.T., Pack, C.C., 2010. The role of V1 surround suppression in MT motion integration. Journal of Neurophysiology 103, 3123–3138. doi:10.1152/jn.00654.2009.

Virtanen, P., Gommers, R., Oliphant, T.E., Haberland, M., Reddy, T., Cournapeau, D., Burovski, E., Peterson, P., Weckesser, W., Bright, J., et al., 2020. SciPy 1.0: Fundamental algorithms for scientific computing in Python. Nature Methods 17, 261–272. doi:10.1038/s41592-019-0686-2.

Wagatsuma, N., Terada, Y., Okuno, H., Ageta-Ishihara, N., 2025. Local connections among excitatory neurons underlie characteristics of enriched environment exposure-induced neuronal response modulation in layers 2/3 of the mouse V1. Frontiers in Systems Neuroscience 19, 1525717. doi:10.3389/fnsys.2025.1525717.

Wang, X.J., Tegnér, J., Constantinidis, C., Goldman-Rakic, P.S., 2004. Division of labor among distinct subtypes of inhibitory neurons in a cortical microcircuit of working memory. Proceedings of the National Academy of Sciences 101, 1368–1373. doi:10.1073/pnas.0305337101.

Whyte, C.J., Müller, E.J., Aru, J., Larkum, M., John, Y., Munn, B.R., Shine, J.M., 2025. A burst-dependent thalamocortical substrate for perceptual awareness. PLOS Computational Biology 21, e1012951. doi:10.1371/journal.pcbi.1012951.

Wibral, M., Priesemann, V., Kay, J.W., Lizier, J.T., Phillips, W.A., 2017. Partial information decomposition as a unified approach to the specification of neural goal functions. Brain and Cognition 112, 25–38. doi:10.1016/j.bandc.2015.09.004.

Williams, L.E., Holtmaat, A., 2019. Higher-order thalamocortical inputs gate synaptic long-term potentiation via disinhibition. Neuron 101, 91–102.e4. doi:10.1016/j.neuron.2018.10.049.

Williams, P.L., Beer, R.D., 2010. Nonnegative decomposition of multivariate information. arXiv preprint arXiv:1004.2515. doi:10.48550/arXiv.1004.2515.

Williams, R.J., 1992. Simple statistical gradient-following algorithms for connectionist reinforcement learning. Machine Learning 8, 229–256. doi:10.1007/BF00992696.

Wilmerding, L.K., Gourianova, F., Grienberger, C., 2026. Instructive signals drive learning through behavioral timescale synaptic plasticity in the hippocampus and beyond. Current Research in Neurobiology 11, 100167. doi:10.1016/j.crneur.2026.100167.

Wilmes, K.A., Clopath, C., 2019. Inhibitory microcircuits for top-down plasticity of sensory representations. Nature Communications 10, 5055. doi:10.1038/s41467-019-12972-2.

Wybo, W.A.M., Tsai, M.C., Tran, V.A.K., Illing, B., Jordan, J., Morrison, A., Senn, W., 2023. NMDA-driven dendritic modulation enables multitask representation learning in hierarchical sensory processing pathways. Proceedings of the National Academy of Sciences 120, e2300558120. doi:10.1073/pnas.2300558120.

Yang, G.R., Murray, J.D., Wang, X.J., 2016. A dendritic disinhibitory circuit mechanism for pathway-specific gating. Nature Communications 7, 12815. doi:10.1038/ncomms12815.

Zhang, S., Xu, M., Kamigaki, T., Hoang Do, J.P., Chang, W.C., Jenvay, S., Miyamichi, K., Luo, L., Dan, Y., 2014. Long-range and local circuits for top-down modulation of visual cortex processing. Science 345, 660–665. doi:10.1126/science.1254126.s

